# Co-substrate pools can constrain and regulate pathway fluxes in cell metabolism

**DOI:** 10.1101/2022.09.05.506656

**Authors:** Robert West, Hadrien Delattre, Elad Noor, Elisenda Feliu, Orkun S Soyer

## Abstract

Cycling of co-substrates, whereby a metabolite is converted among alternate forms via different reactions, is ubiquitous in metabolism. Several cycled co-substrates are well known as energy and electron carriers (e.g. ATP and NAD(P)H), but there are also other metabolites that act as cycled co-substrates in different parts of central metabolism. Here, we develop a mathematical framework to analyse the effect of co-substrate cycling on metabolic flux. In the cases of a single reaction and linear pathways, we find that co-substrate cycling imposes an additional flux limit on a reaction, distinct to the limit imposed by the kinetics of the primary enzyme catalysing that reaction. Using analytical methods, we show that this additional limit is a function of the total pool size and turnover rate of the cycled co-substrate. Expanding from this insight and using simulations, we show that regulation of co-substrate pool size can allow regulation of flux dynamics in branched and coupled pathways. To support theses theoretical insights, we analysed existing flux measurements and enzyme levels from the central carbon metabolism and identified several reactions that could be limited by co-substrate cycling. We discuss how the limitations imposed by co-substrate cycling provide experimentally testable hypotheses on specific metabolic phenotypes. We conclude that measuring and controlling co-substrate pools is crucial for understanding and engineering the dynamics of metabolism.

## INTRODUCTION

Dynamics of cell metabolism directly influences individual and population-level cellular responses. Examples include metabolic oscillations underpinning the cell cycle (1,2) and metabolic shifts from respiration to fermentation, observed in cancer phenotypes (3-5) and cell-to-cell cross-feeding (6-8). Predicting or conceptualising these physiological responses using dynamical models, however, is difficult due to the large size and high connectivity of cellular metabolism. Despite this complexity, cellular metabolism might feature simplifying ‘design principles’ that determine the overall dynamics.

There is ongoing interest in finding such simplifying principles. Early studies developed a theory of metabolic pathway structure, concerning the position of ATP generating steps in a linear pathway, under the assumption of pathway flux optimisation with limited enzyme production capacity (9). This theory predicted a trade-off between pathway flux and yield (net ATP generation) (10), which is used to explain the emergence of different metabolic phenotypes (11). In related studies, several specific models pertaining to enzyme allocation and optimality have been developed to explain the structure of different metabolic pathways (12), and the metabolic shifting from respiration to fermentative pathways under increasing glycolysis rates (8, 13, 14).

Another conceptual framework emphasized the importance of co-substrate cycling, rather than net production (e.g. of ATP), as a key to understanding metabolic systems (15). This framework is linked to the idea of considering the supply and demand structures around specific metabolites as regulatory blocks within metabolism (16). For example, the total pool of ATP and its derivates (the ‘energy charge’) is suggested as a key determinant of physiological cell states (17). Inspired by these ideas, early theoretical studies have shown that metabolic systems featuring metabolite cycling together with allosteric regulation can introduce switch-like and bistable dynamics (18, 19), and that metabolite cycling motifs introduce total co-substrate level as an additional control element in metabolic control analysis (20, 21). Specific analyses of ATP cycling in the glycolysis pathway, sometimes referred to as a ‘turbo-design’, and metabolite cycling with autocatalysis, as seen for example in glyoxylate cycle, have shown that these features constrain pathway fluxes (22-27). Taken together, these studies indicate that metabolite cycling, in general, and co-substrate cycling specifically, could provide a key ‘design feature’ in cell metabolism, imposing certain constraints or dynamical properties to it.

Towards better understanding the role of co-substrate cycling in cell metabolism dynamics, we undertook here an analytical and simulation-based mathematical study together with analyses of measured fluxes. We created models of enzymatic reaction systems featuring co-substrate cycling, abstracted from real metabolic systems such as glycolysis, nitrogen-assimilation, and central carbon metabolism. We found that co-substrate cycling introduces a fundamental constraint on reaction flux. In the case of single reaction and short linear pathways, we were able to derive a mathematical expression of the constraint, showing that it relates to the pool size and turnover rate of the co-substrate. Analysing measured fluxes, we find that several of the co-substrate featuring reactions in central carbon metabolism carry lower fluxes than expected from the kinetics of their primary enzymes, suggesting that these reactions might be limited by co-substrate cycling. In addition to its possible constraining role, we show that co-substrate cycling can also act as a regulatory element, where control of co-substrate pool size can allow control of flux dynamics across connected or branching pathways. Together, these findings show that co-substrate cycling can act both as a constraint and a regulatory element in cellular metabolism. The resulting theory provides testable hypotheses on how to manipulate metabolic fluxes and cell physiology through the control of co-substrate pool sizes and turnover dynamics and can be expanded to explain dynamic measurements of metabolite concentrations in different perturbation experiments.

## RESULTS AND DISCUSSION

### Co-substrate cycling is a ubiquitous motif in metabolism

Certain metabolites can be consumed and reproduced via different reactions in the cell, thereby resulting in their ‘cycling’ (Fig. 1A). This cycling creates interconnections within metabolism, spanning either multiple reactions in a single, linear pathway, or multiple pathways that are independent or are branching from common metabolites. For example, in glycolysis, ATP is consumed in reactions mediated by the enzymes glucose hexokinase and phosphofructokinase, and is produced by the downstream reactions mediated by phosphoglycerate and pyruvate kinase (Fig. S1A). In the nitrogen assimilation pathway, the NAD^+^ / NADH pair is cycled by the enzymes glutamine oxoglutarate aminotransferase and glutamate dehydrogenase (Fig. S1B). Many other cycling motifs can be identified, involving either metabolites from the central carbon metabolism or metabolites that are usually referred to as co-substrates. Examples for the latter include NADPH, FADH^2^, GTP, and Acetyl-CoA and their corresponding alternate forms, while examples for the former include the tetrahydrofolate (THF) / 5,10-Methylene-THF and glutamate / α -oxoglutarate (akg) pairs involved in one-carbon transfer and in aminoacid biosynthesis pathways, respectively (Fig. S1C & D). For some of these metabolites, their cycling can connect many reactions in the metabolic network. Taking ATP (NADH) as an example, there are 265 (118) and 833 (601) reactions linked to the cycling of this metabolite in the genome-scale metabolic models of *Escherichia coli* and human respectively (models iJO1366 (28) and Recon3d (29)).

**Figure 1.**
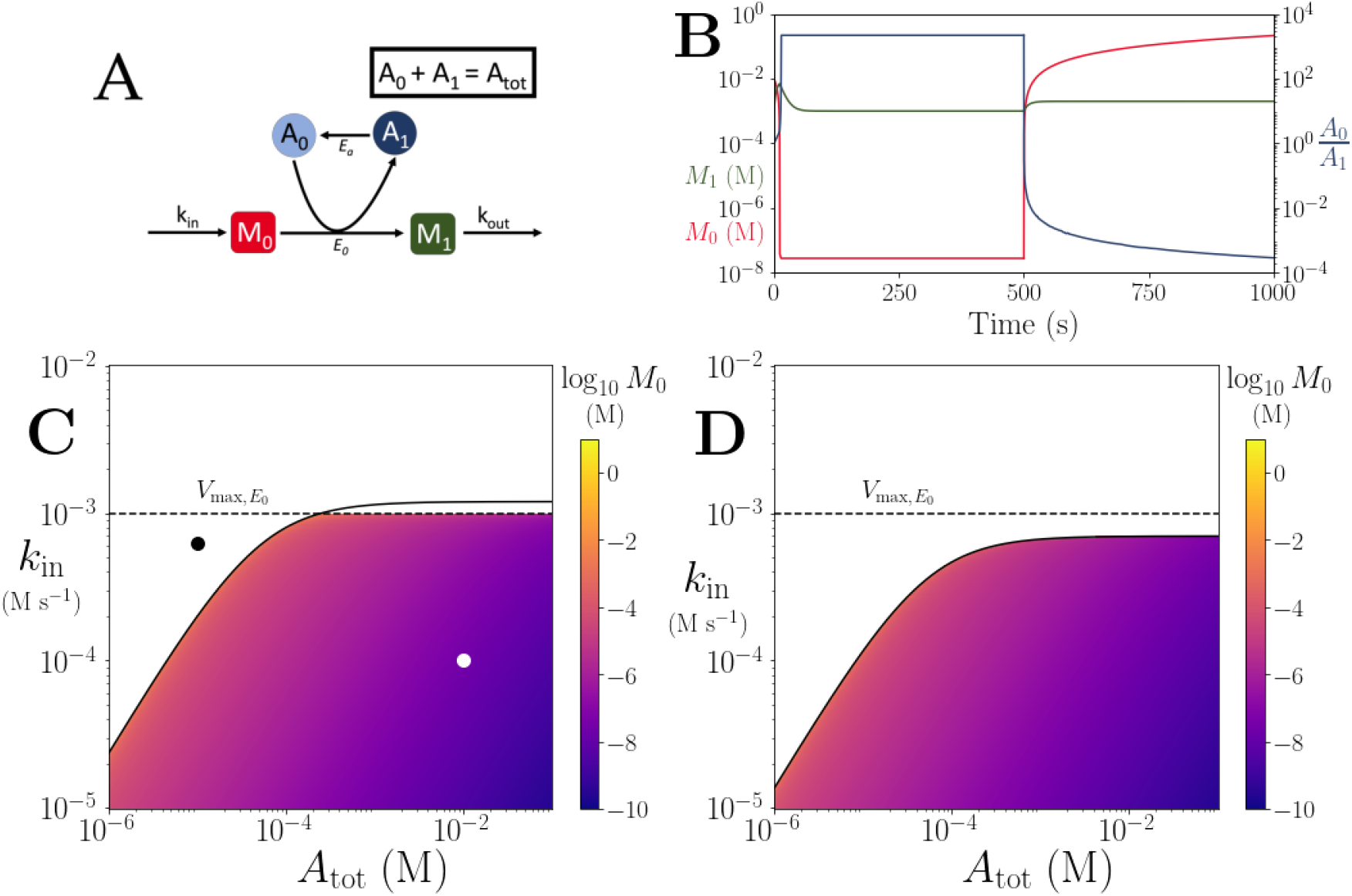
**(A)** Cartoon representation of a single irreversible reaction with co-substrate cycling (see *SI* for other reaction schemes). The co-substrate is considered to have two forms *A*_*0*_ and *A*_*1*_. **(B)** Concentrations of *M*_0_ (red) and *M*_1_ (green) and *A*_0_/*A*_1_ ratio (blue) as a function of time. At t = 500, the parameters are switched from the white dot in panel (C) (where a steady state exists) to the black dot (where we see continual build-up of *M*_0_ and decline of *A*_0_ without steady state). **(C & D)** Heatmap of the steady state concentration of *M*_*0*_ as a function of the total co-substrate pool size (*A*_*tot*_) and inflow flux (*k*_in_). White area shows the region where there is no steady state. On both panels, the dashed line indicates the limitation from the primary enzyme, *k*_*in*_ < *V*_*max,E0*_, and the solid line indicates the limitation from co-substrate cycling, *k*_*in*_ < *A*_*tot*_ *V*_*max,EA*_ */ (K*_*m,EA*_ *+ A*_*tot*_). In panel (C), there is a range of *A*_*tot*_ values for which the first limitation is more severe than the second. In contrast, in panel (D), the second limitation is always more severe than the first. In (B & C) the parameters used for the primary enzyme (for the reaction converting *M*_*0*_ into *M*_*1*_) are picked from within a physiological range (see *Supplementary File* 1) and are set to: *E*_tot_ = 0.01mM, *k*_cat_ = 100 s^− 1^, *K*_*m,E0*_ = *K*_*m,EA*_ =50μ M, while *k*_out_ is set to 0.1s^− 1^. The *E*_tot_ and *k*_cat_ for the co-substrate cycling enzyme are 1.2 times those for the primary enzyme. In panel (D) the parameters are the same except for and *E*_tot_ and *k*_cat_ for the co-substrate cycling enzyme, which are set to 0.7 times those for the primary enzyme.

### Cycled co-substrates can act as ‘conserved moieties’ for metabolic flux dynamics

Cycling of co-substrate results in their turnover across their different forms e.g., NAD^+^ and NADH. The total pool-size involving all the different forms of a cycled metabolite, however, can approach a constant value at steady state. In other words, the total concentration of a cycled metabolite across its different forms at steady state would be given by a constant defined by the ratio of the influx and outflux rates (see *Supplementary Information* (*SI*), section 2 and 3). In other words, the cycled metabolite would become a ‘conserved moiety’ for the rest of the metabolic system and can have a constant ‘pool size’. Supporting this, temporal measurement of specific co-substrate pool sizes shows that ATP and GTP pools are constant under stable metabolic conditions, but can rapidly change in response to external perturbations, possibly through inter-conversions among pools rather than through biosynthesis (30).

### Co-substrate cycling introduces a limitation on reaction flux

To explore the effect of co-substrate cycling on pathway fluxes, we first consider a didactic case of a single reaction. This reaction converts an arbitrary metabolite *M*_*0*_ to *M*_*1*_ and involves co-substrate cycling (Fig. 1A). For co-substrate cycling, we consider additional ‘background’ enzymatic reactions that are independent of *M*_*0*_ and can also convert the co-substrate (denoted *E*_*A*_ on Fig. 1A). We use either irreversible or reversible enzyme dynamics to build an ordinary differential equation (ODE) kinetic model for this reaction system and solve for its steady states analytically (see *Methods* and *SI*, section 3). In the case of using irreversible enzyme kinetics, we obtain that the steady state concentration of the two metabolites, *M*_*0*_ and *M*_*1*_ (denoted as *m*_*0*_ and *m*_*1*_) are given by:

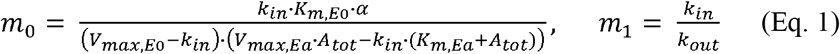

where *k*_*in*_ and *k*_*out*_ denote the rate of in-flux of *M*_*0*_, and out-flux of *M*_*1*_, either in-and-out of the cell or from other pathways, and *A*_*tot*_ denotes the total pool size of the cycled metabolite (with the different forms of the cycled metabolite indicated as *A*_*0*_ and *A*_*1*_ in Fig. 1A). The term α is a positive expression comprising *A*_*tot*_, and the kinetic parameters of the enzymes in the model (see *SI*). The parameters *V*_*max,Ea*_ and *V*_*max,E0*_ are the maximal rates (i.e. *V*_*max*_ = *k*_*cat*_. *E*_*tot*_) for the enzymes catalysing the conversion of *A*_*0*_ and *M*_*0*_ into *A*_*1*_ and *M*_*1*_ (enzyme *E*_*0*_), and the turnover of *A*_*1*_ into *A*_*0*_ (enzyme *E*_*a*_), respectively, while the parameters *K*_*m,Ea*_ and *K*_*m,E0*_ are the individual or combined Michaelis-Menten coefficients for these enzymes’ substrates (i.e. for *A*_*0*_ and *M*_*0*_ and *A*_*1*_, respectively). The steady states for the model with all enzymatic conversions being reversible, and for a model with degradation and synthesis of *A*_*0*_ and *A*_*1*_, are given in the *SI*. The steady state solutions of these alternative models are structurally akin to Eq. 1, and do not alter the qualitative conclusions we make in what follows.

A key property of Eq. 1 is that it contains terms in the denominator that involve a subtraction. The presence of these terms introduces a limit on the parameter values for the system to attain a positive steady state. Specifically, we obtain the following conditions for positive steady states to exist:

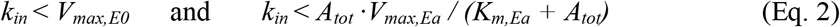

Additionally, the ‘shape’ of Eq. 1 indicates a ‘threshold effect’ on the steady state value of *m*_*0*_, where it would rise towards infinity as *k*_*in*_ increases towards the lower among the limits given in Eq. 2 (see Fig. 1B).

Why does Eq. 1 show this specific form, leading to these limits? We find that this is a direct consequence of the steady state condition, where metabolite production and consumption rates need to be the same at steady state. In the case of co-substrate cycling, the production rate of *M*_*0*_ is given by *k*_*in*_, while its consumption rate is a function of the concentration of *A*_*0*_ and the *V*_*max,E0*_. The concentration of *A*_*0*_ is determined by its re-generation rate (which is a function of *K*_*m,EA*_ and *V*_*max,Ea*_) and the pool size (*A*_*tot*_). This explains the inequalities given in Eq. 2 and shows that a cycled co-substrate, when acting as a conserved moiety, creates the same type of limitation (mathematically speaking) on the flux of a reaction it is involved in, as that imposed by the enzyme catalysing that reaction (*E*_*0*_ in this example) (see Fig. 1C&D). We also show that considering the system shown in Fig. 1A as an enzymatic reaction without co-substrate cycling leads to only the constraint *k*_*in*_ *< V*_*max,E0*_, while when considering it as a non-enzymatic reaction with co-substrate cycling only, the constraint *k*_*in*_ < *A*_*tot*_ · *V*_*max,Ea*_ */ (K*_*m,Ea*_ *+ A*_*tot*_) becomes the sole limitation on the system (see *SI*, section 3). In other words, the two limitations act independently.

To conclude this section, we re-iterate its main result. The flux of a reaction involving co-substrate cycling is limited either by the kinetics of the primary enzyme mediating that reaction, or by the turnover rate of the co-substrate. The latter is determined by the co-substrate pool size and the kinetics of the enzyme(s) mediating its turnover.

### Co-substrate cycling causes a flux limit on linear metabolic pathways

We next considered a generalised, linear pathway model with *n*+1 metabolites and arbitrary locations of reactions for co-substrate cycling, for example as seen in upper glycolysis (Fig. S1). In this model, we only consider intra-pathway metabolite cycling, i.e. the co-substrate is consumed and re-generated solely by the reactions of the pathway. Here, we show results for this model with 5 metabolites as an illustration (Fig. 2A), while the general case is presented in the *SI* section 4.

**Figure 2.**
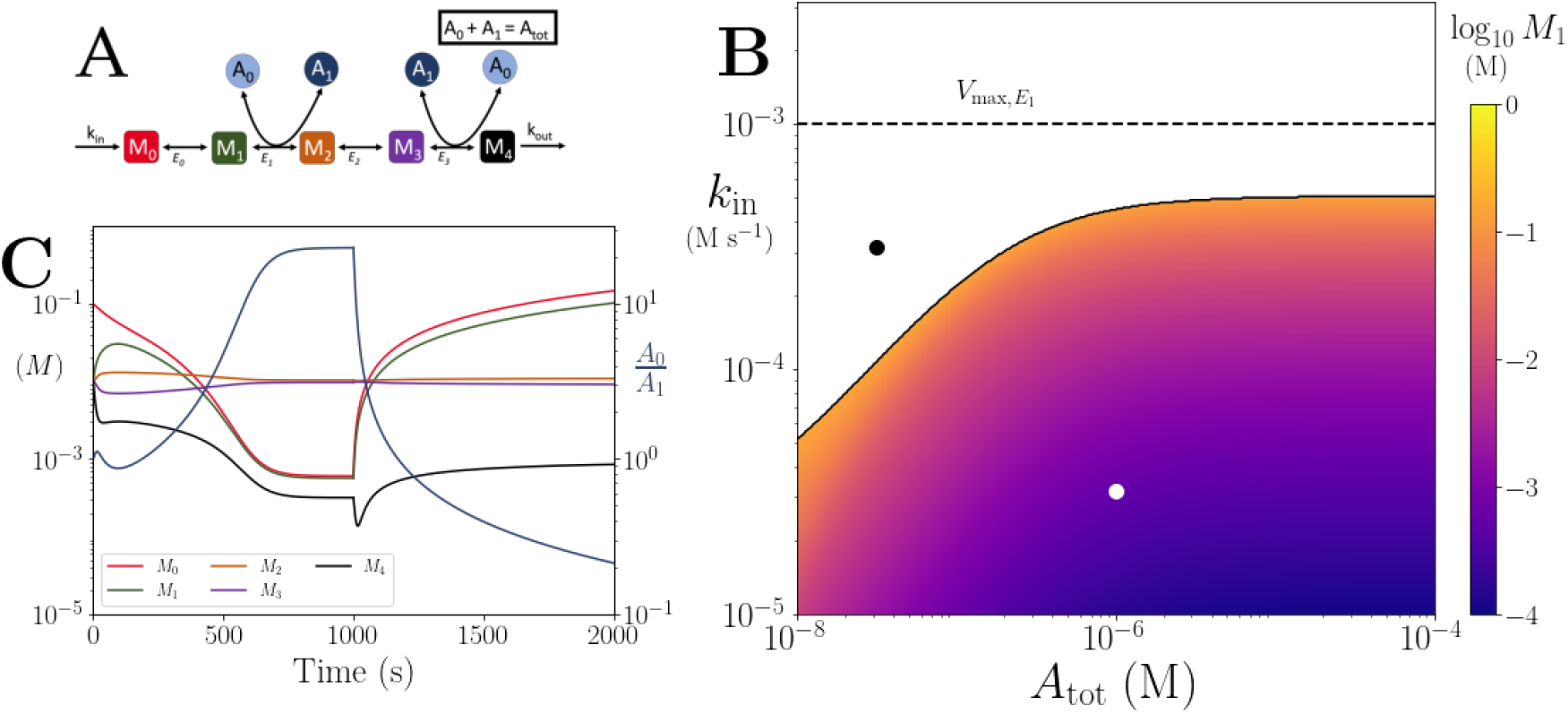
**(A)** Cartoon representation of a chain of reversible reactions with co-substrate cycling occurring solely inter-pathway. The co-substrate is considered to have two forms *A*_*0*_ and *A*_*1*_. **(B)** Heatmap of the steady state concentration of *M*_*0*_ as a function of the total metabolite pool size (*A*_*tot*_) and inflow rate constant (*k*_in_). White area shows the region where there is no steady state. The dashed and solid lines indicate the limitations arising from primary enzyme (*E*_*1*_ in this case) and co-substrate cycling, respectively, as in Fig. 1. **(C)** Concentrations of *M*_0-4_, and *A*_0_/*A*_1_ ratio as a function of time (with colors as indicated in the inset). At t = 1000, the parameters are switched from the white dot in panel (B) (where a steady state exists) to the black dot (where we see build-up of all substrates that are produced before the first co-substrate cycling reaction, and continued decline of *A*_0_). The parameters used are picked from within a physiological range (see *Supplementary File* 1) and are set to: *E*_tot_ = 0.01mM, *k*_cat_ = 100 s^− 1^, *K*_*m*_ =50μ M, for all reactions, and *k*_out_ = 0.1s^− 1^.

We find the same kind of threshold dynamics as in the single reaction case. When *k*_*in*_ is above a threshold value, the metabolite *M*_*0*_ accumulates towards infinity and the system does not have a steady state (Fig. 2B). A numerical analysis, as well as our analytical solution, reveals that the accumulation of metabolites applies to all metabolites upstream of the first reaction with co-substrate cycling (Fig. 2C and *SI* section 4). Additionally, metabolites downstream of the cycling reaction accumulate to a steady state level that does not depend on *k*_*in*_ (Fig. 2C and Fig. S2). In other words, pathway output cannot be increased further by increasing *k*_*in*_ beyond the threshold. Finally, as *k*_*in*_ increases, the cycled metabolite pool shifts towards one form and the ratio of the two forms approaches zero (Fig. 2C).

An analytical expression for the threshold for *k*_*in*_, like shown in Eq. 2, could not be derived for linear pathways with *n* > 3, but our analytical study indicates that (i) the threshold is always linked to *A*_*tot*_ and enzyme kinetic parameters, and (ii) the concentration of all metabolites upstream (downstream) to the reaction coupled to metabolite cycling will accumulate towards infinity (a fixed value) as *k*_*in*_ approaches the threshold (see *SI* section 4). In Figure 2, we illustrate these dynamics with simulations for a system with *n*=4.

We also considered several variants of this generalised linear pathway model, corresponding to biologically relevant cases as shown in Fig. S1. These included (i) intra-pathway cycling of two different metabolites, as seen with ATP and NADH in combined upper glycolysis and fermentation pathways (Fig. S3, *SI* section 5), (ii) different stoichiometries for consumption and re-generation reactions of the cycled metabolite, as seen in upper glycolysis (Fig. S4, *SI* section 6), and (iii) cycling of one metabolite interlinked with that of another, as seen in nitrogen assimilation (Fig. S5, *SI* section 7). The results in the *SI* confirm that all these cases display similar threshold dynamics, where the threshold point is a function of the co-substrate pool size and the enzyme kinetics.

### Cycled metabolite related limit could be relevant for specific reactions from central metabolism

Based on flux values that are either experimentally measured or predicted by flux balance analysis (FBA), many reactions from the central carbon metabolism are shown to have lower flux than expected from the kinetics of their immediate enzymes (31). In other words, these reactions carry fluxes below the first limit identified above in Eq. 2. While substrate limitation and thermodynamic effects can partially explain such lower flux in some cases (31), the presented theory suggests that limitation due to co-substrate turnover could also be a contributing factor.

To explore this possibility, we re-analysed the flux values compiled previously (31, 32) and focussed solely on reactions that are linked to ATP, NADH, or NADPH pools (see *Methods* and *Supplementary File 1*). The resulting dataset contained fluxes, substrate concentrations, and enzyme levels for 45 different reactions determined under 7 different conditions along with turnover numbers and kinetic constants of the corresponding enzymes. In total, we gathered 49 combinations of enzyme-flux-*k*_*cat*_ values with full experimental data and 259 combinations with only FBA-predicted flux values. We compared the flux values that would be expected from the primary enzyme limit identified above, under all conditions analysed (Fig. 3A), and in addition checked whether the saturation effect of the primary substrate could explain the difference (Fig. 3B). We found that in both cases, about 80% of these reactions carry flux lower than what is expected from enzyme kinetics (Fig. S6), suggesting that the limits imposed by co-factor dynamics might be constraining the flux further. The low number of the cases where the flux exceeds the limit might be due to uncertainties in measurement of flux, enzyme or substrate level.

**Figure 3.**
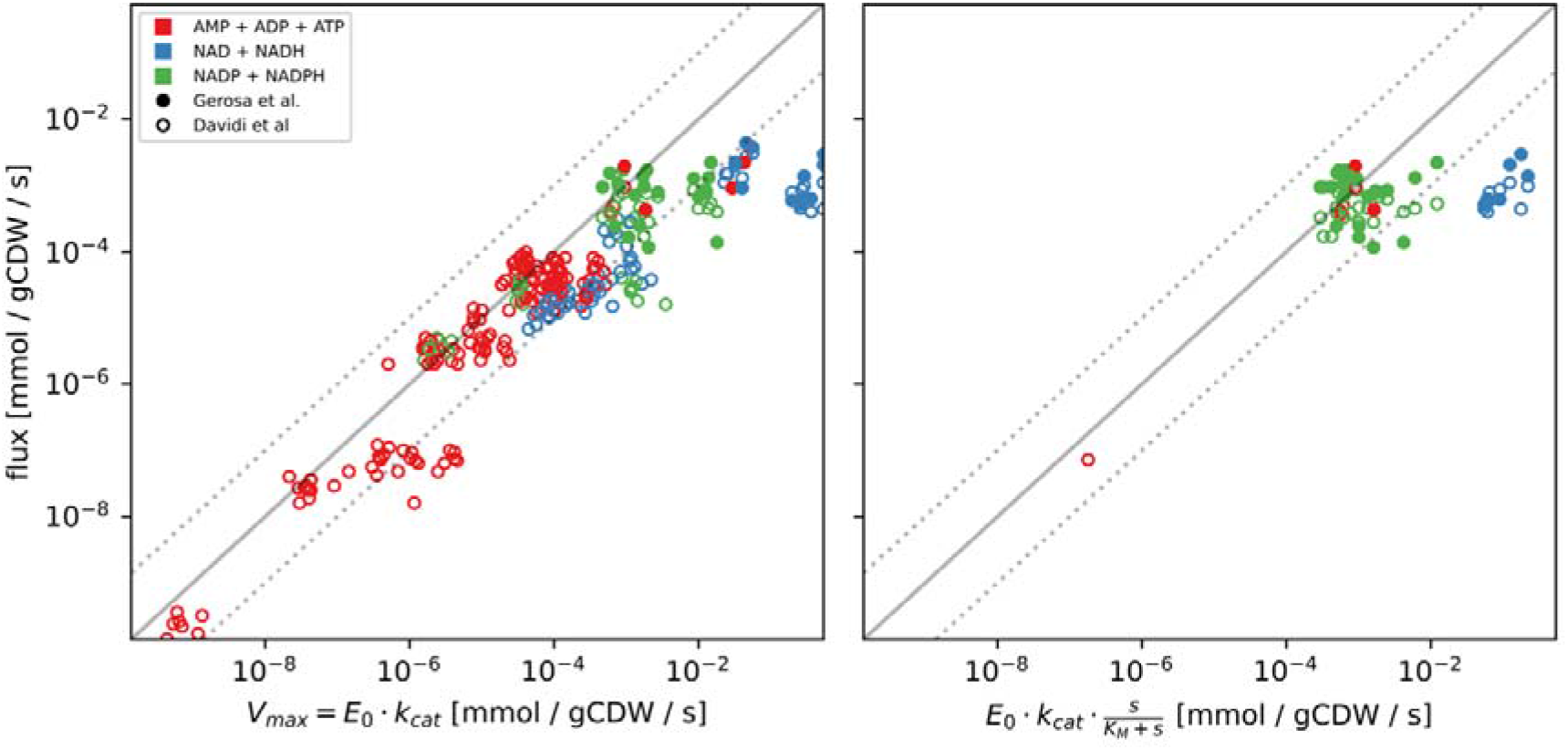
**(A)** Measured and FBA-predicted flux values (from (31, 32)) plotted against the calculated primary enzyme kinetic threshold (first part of Eq. 1). Notice that there are 7 points for each reaction, corresponding to the different experimental conditions under which measurements or FBA modelling was done (see *Supplementary File* S1 for data, along with reaction names and metabolites involved). **(B)** Measured flux values (from (31, 32)) plotted against the calculated primary enzyme kinetic threshold (first part of Eq. 1) adjusted by substrate affinity of the enzyme. Note that the flux data shown here is a subset of the flux data presented in (A), focusing only on those where the main substrate concentration was experimentally measured and the relevant *K*_*m*_ is known. For both panels, the solid line indicates the equivalence of the two values and the dashed lines indicate 10% interval on this, as a guide to the eye. Point color indicates the nature of co-substrate involved and fill state indicates the data source (as shown on the inset).

To further support the hypothesis that co-substrate turnover dynamics contribute to the flux limitation, we checked the relation between fluxes and co-substrate pool sizes, which change among different conditions. For both measured and FBA-predicted fluxes, we find that several reactions show significant correlation between flux and co-substrate pool size (see Table S1, *SI* section 8). In the case of FBA-predicted fluxes, however, we note that these results can be confounded due to additional, flux-to-flux correlations and correlations between pool sizes and growth rate. Among reactions with measured fluxes, the two reactions with high correlation to pool size are those mediated by malate dehydrogenase (*mdh*), linked with NADH pool, and phosphoglycerate kinase (*pgk*), linked with the ATP pool.

### Co-substrate cycling allows regulation of branch point fluxes

In addition to its possible constraining effects on fluxes, we wondered if co-substrate dynamics can offer a regulatory element in cellular metabolism. In particular, co-substrate cycling can commonly interconnect two independent pathways, or pathways branching from the same upstream metabolite, where it could influence flux distributions among those pathways. To explore this idea, we considered a model of a branching pathway, with each branch involving a different co-substrate, *A* and *B* (Fig. 4A and *SI* section 8). This scenario is seen in synthesis of certain amino acids that start from a common precursor but utilise NADH or NADPH, for example Serine and Threonine.

**Figure 4:**
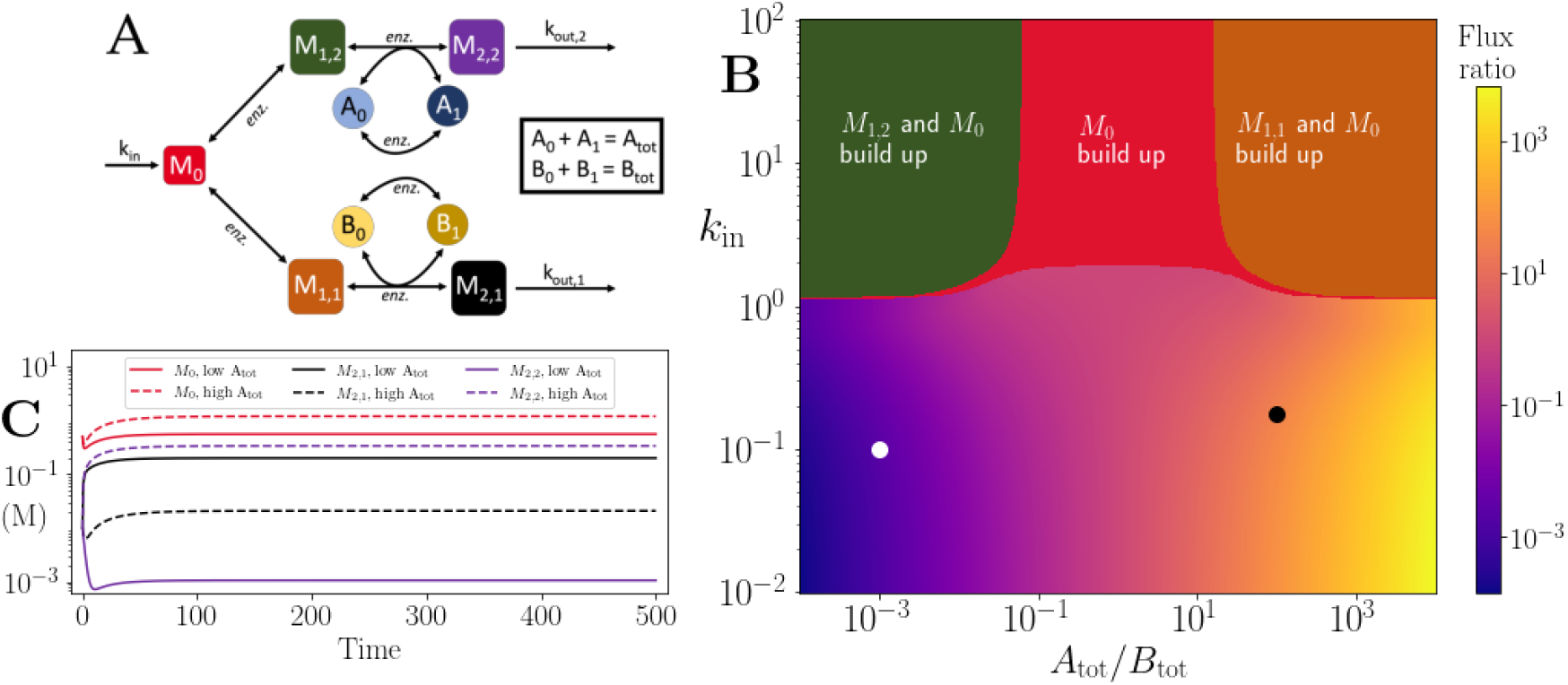
**(A)** Cartoon representation of two branching pathways from the same upstream metabolite. The two branches are linked to separate co-substrate pools, *A* and *B*. Note that pathway independent turnover of the co-substrates is included in the model (see *Supplementary File 2*). **(B)** The pathways’ flux ratio (i.e. flux into *M*_*2,2*_ divided by flux into *M*_*2,1*_) shown in colour mapping, against the ratio of co-substrate pool sizes, *A*_*tot*_ and *B*_*tot*_, and the influx rate, *k*_*in*_, into the upstream metabolite. In the block colour areas, the system has no steady state and the indicated metabolite(s) *M*_*0*_ and one of the metabolites *M*_*1,2*_ or *M*_*1,1*_ accumulate towards infinity. **(C)** Concentrations of upstream and branch-endpoint metabolites over time, coloured as shown in the inset of the panel. The solid lines show results using parameters indicated by the white dot in panel (B), where *B*_*tot*_ > *A*_*tot*_, while the dashed lines show results using parameters indicated by the black dot in panel (B), where *A*_*tot*_ > *B*_*tot*_. For both simulations, all kinetic parameters are arbitrarily set to 1, apart from the pathway-independent co-substrate recycling (*V*_*max,Ea*_) that is set to 10 (see *Supplementary File 2*).

We hypothesised that regulating the two co-substrate pool sizes, *A*_*tot*_ and *B*_*tot*_, could allow regulation of the fluxes on the two branches. To test this hypothesis, we run numerical simulations with different co-substrate pool sizes and influx rates into the branch point. We found that the ratio of fluxes across the two branches can be regulated by changing the ratio of *A*_*tot*_ to *B*_*tot*_ (Fig. 4B). The regulation effect is seen with a large range of *k*_*in*_ values, but the threshold effect is still present with high enough *k*_*in*_ values leading to loss of steady state and metabolite build up. In that case, the resulting metabolite build-up can affect either branch depending on which co-substrate has the lower pool size (see upper corner regions on Fig. 4B). There is also a regime of only the upstream, branch point metabolite building-up, but this happens only when all reactions are considered as reversible and the extent of it depends on turnover rates of the two co-substrates (Fig S7 and *SI* section 8).

In the no-build-up, steady state regime, changing the pool size ratio of the two co-substrates causes a change in fluxes and metabolite levels, The change in flux ratio is of the same order as the change in pool size ratio (Fig. 4C & D), while the change in the ratio of metabolite levels is in general less. This relation between pool size ratio and flux ratio on each branch is unaffected by the value of *k*_*in*_. We also evaluated the level of regulation that can be achieved by varying the turnover rates of *A* and *B*. The flux regulation effect in this case is weaker, unless the difference in the turnover rates is large and the influx rate is close to the threshold (Fig. S8).

### Inter-pathway co-substrate cycling limits maximum influx difference and allows for correlating pathway outfluxes despite influx noise

We next considered a simplified model of two independent pathways interconnected by a single co-substrate pool (Fig. 5A and *SI* section 9). This model can represent several different processes in metabolism, for example the coupling between the TCA cycle and the respiratory electron transfer chain, through NADH generation and consumption respectively, or the coupling between the pentose phosphate pathway and some amino acid biosynthesis pathways (notably Methionine), through NADPH generation and consumption respectively (S1F). We hypothesised that such inter-pathway co-substrate cycling might cause the co-substrate related limit to relate to dfference in pathway influxes, rather than input into one pathway, and also balance the pathway output fluxes against influx fluctuations.

**Figure 5:**
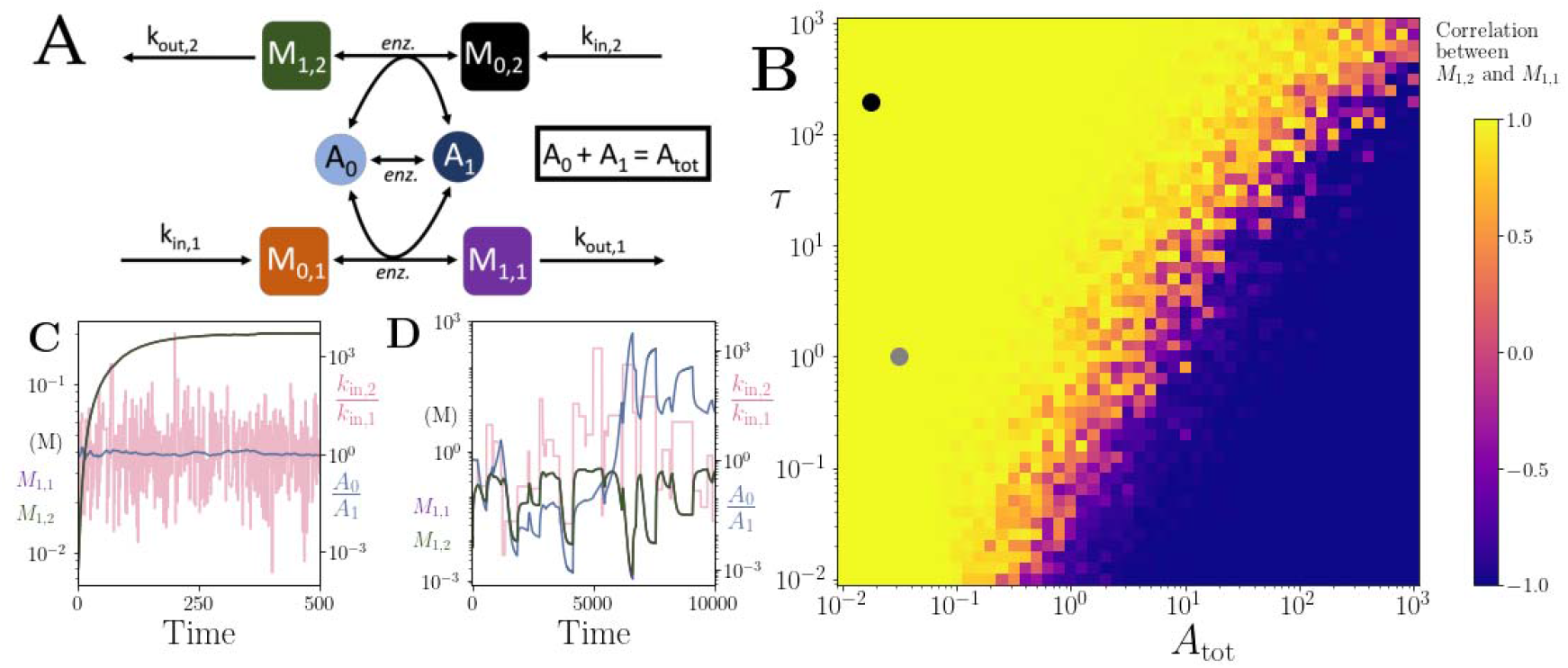
**(A)** Cartoon representation of two pathways coupled via the same co-substrate cycling. The two forms of the co-substrate are indicated as *A*_*0*_ and *A*_*1*_. It is converted from *A*_*0*_ to *A*_*1*_ on the lower pathway, and from *A*_*1*_ to *A*_*0*_ in the upper pathway. The presented results are for a model with reversible enzyme kinetics, while the results from a model with irreversible enzyme kinetics are shown in Fig. S9. **(B)** Correlation coefficient of the two pathway product metabolites, *M*_*1,2*_ and *M*_*1,1*_, as a function of the total amount of co-substrate (*A*_*tot*_) and the extent of fluctuations in the two pathway influxes, *k*_*in,1*_ and *k*_*in,2*_. The influx fluctuation is characterised by a waiting time that is exponentially distributed with mean τ, after which the log ratio of the *k*_*in*_ values is drawn from a standard normal distribution. The mean of the *k*_*in*_ values is set to be 0.1 and the pathway-independent cycling occurs at a much lower rate compared to the other reactions (see *Supplementary File 3*). **(C)** Concentrations of metabolites *M*_*1,2*_ (green) and *M*_*1,1*_ (magenta), pathway influx ratio (pink), and *A*_0_/*A*_1_ ratio (blue) as a function of time. The simulation is run with parameters corresponding to the grey dot in (B) where the products are correlated, and the rate of *k*_*in*_ fluctuations is on a similar timescale to the other reactions. The system is largely unresponsive to the noise. **(D)** Concentrations of metabolites *M*_*1,2*_ (green) and *M*_*1,1*_ (magenta), pathway influx ratio (pink), and *A*_0_/*A*_1_ ratio (blue) as a function of time. The simulation is run with parameters corresponding to the black dot in (B) where the products are correlated, but the fluctuations in *k*_*in*_ values occur at a much lower rate than the other reactions. For both simulations, all kinetic parameters are arbitrarily set to 1, apart from the pathway-independent co-substrate recycling (*V*_*max,Ea*_) that is set to 0.01 (see *Supplementary File 3*).

To address the first hypothesis, we used analytical methods and explored the relation between the systems’ ability to reach steady state and system parameters. We found that, indeed, for this coupled system, the ability to reach steady state depends on the influx difference between two pathways (Fig. S9). This dependence is given by a composite function of the total pool size and the kinetic parameters relating to pathway-independent turn-over of the co-substrate (see *SI*, section 10).

To test the second hypothesis about the output balancing, we considered the correlation of the steady-state outputs of the pathways with random fluctuations in their influx (Fig. 5B). As the pool size decreases, the system reaches a point where there is a transition from anti-correlation to high correlation in product output (blue to yellow region in Fig. 5B). At low pool sizes, pathway outputs are fully correlated despite significant fluctuation in pathway influx (Fig. 5C, D). Within this correlated regime, we identified two different sub-regimes. The first is a regime where the metabolite concentrations stay relatively constant despite the influx noise (Fig 5C). This regime arises because the influx fluctuations are occurring at a much faster rate than the pathway kinetics and the system is rather non-responsive to influx noise. In a second regime, the influx noise is at time scales comparable to pathway kinetics. Here, the metabolite concentrations can readily change with the influx changes, and the system is ‘responsive’, yet the output levels are still correlated (Fig. 5D). This regime is directly a result of co-substrate cycling dynamics. Because the turnover of co-substrate is essentially coupling the two pathways, their outputs become directly correlated. This effect does not depend on whether pathway reactions are modelled as reversible or irreversible, but on the rate of the assumed background, i.e. pathway-independent turnover of the co-substrate (Fig. S10).

These results show that coupling by co-substrate cycling can introduce a limit on influxes of independent pathways or metabolic processes. Furthermore, such coupling can allow high correlation in the pathway outputs, despite significant noise in the inputs of those pathways. These effects are most readily seen where the turnover of the coupling co-substrate by other processes is low. We note that an example case for such a scenario is the coupling of respiration and oxidative phosphorylation, where transmembrane proton cycling takes the role of the cycled co-substrate (33).

## CONCLUSIONS

We presented a mathematical analysis of metabolic systems featuring co-substrate cycling and showed that such cycling introduces a threshold effect on system dynamics. As the pathway’s influx rate, *k*_*in*_, approaches a threshold value, the steady state concentrations of metabolites that are upstream of a reaction linked to co-substrate cycling, increase towards infinity and the system cannot reach steady state. Specifically, for reactions involving co-substrates, there are two thresholds on influx rate, one relating to the kinetics of the enzyme directly mediating that reaction, and another relating to the kinetics of the enzymes mediating the turnover of the co-substrate and the pool size of that co-substrate.

This second, additional constraint arising from co-substrate cycling can be directly relevant for cell physiology. We particularly note that this threshold can be highly dynamic, and condition- and cell-dependent. When cells have a permanently or occasionally lowered total co-substrate pool size (i.e. lower *A*_*tot*_), or when they are placed under challenging conditions (e.g. high carbon- or nitrogen-source concentrations) causing higher *k*_*in*_ values across various pathways, their metabolic systems can be close to the threshold presented here. While both *k*_*in*_ and *A*_*tot*_ can be adjusted in the long term, for example by reducing substrate transporter expression or increasing co-substrate biosynthesis, there can be short term impact on cells experiencing significant flux limitations and metabolite accumulations.

These results could contribute to our understanding of two commonly observed metabolic dynamics that arise under increasing or high substrate concentrations, and that are shown to cause either ‘substrate-induced death’ (24) or ‘overflow metabolism’. The latter usually refers to a respiration-to-fermentation switch under respiratory conditions (e.g. the Warburg and Crabtree effects (3, 4, 12, 34)), but other types of overflow metabolism, involving excretion of amino acids and vitamins, has also been observed (6, 35). Several arguments have been put forward to explain these observations, including osmotic effects arising from high substrate concentrations causing cell death and limitations in respiratory pathways or cell’s protein resources causing a respiration-to-fermentation switch (4, 12, 13).

Notwithstanding the possible roles of these processes, the presented theory leads to the hypothesis that both substrate-induced death and metabolite excretions could relate to increasing substrate influx rate reaching close to the limits imposed by co-substrate dynamics. There is experimental support for this hypothesis in the case of both observations. Substrate-induced death and associated mutant phenotypes are linked to the dynamics associated with ATP regeneration in glycolysis (22-24). Based on that finding, it has been argued that cells aim to avoid the threshold dynamics through allosteric regulation of those steps of the glycolysis that involve ATP consumption (23). In the case of respiration-to-fermentation switch, it has been shown that the glucose influx threshold, at which fermentative overflow starts, changes upon introducing additional NADH conversion reactions in both yeast and *E. coli* populations (36, 37). In another supportive case, sulfur-compound excretions are linked to alterations in NAD(P)H pool through changes in the amino acid metabolism (38, 39).

Dynamical thresholds relating to co-substrate pools would be relevant for all co-substrates, and not just for ATP or NADH, which have been the focus of most experimental studies to date. We would expect that altering kinetics of enzymes involved in co-substrate cycling can have direct impact on cell physiology, and in particular on metabolic excretions. This prediction can be tested by exploring the effect of mutations on enzymes linked to co-substrate consumption and production, or by altering co-substrate pool sizes and assessing effects of such perturbations on the dynamics of metabolic excretions. These tests can be experimentally implemented by introducing additional enzymes specialising in co-substrate consumption or production (e.g. ATPases, oxidases, or other) and controlling their expression. It would also be possible to monitor co-substrate pool sizes in cells in real time by using fluorescent sensors on key metabolites such as ATP or glutamate, or by measuring autofluorescence of certain pool metabolites, such as NAD(P)H, under alterations to influx rate of glucose or ammonium.

Besides acting as a flux constraint, we find that co-substrate pools can also allow for regulation of pathway fluxes through regulation of pool size or turnover dynamics. We find that such regulation can take the form of balancing inter-connected pathways, thereby ensuring correlation between outputs of different metabolic processes, or regulating flux across branch points. Regulation of fluxes through co-substrate pools can act to adjust metabolic fluxes at time scales shorter than possible via gene regulation, and possibly at similar time scales as with allosteric regulation – especially when considering pool size alterations through exchange among connected pools. Possibility of such a regulatory role has been indicated experimentally, where total ATP pool size is found to change when yeast cells are confronted with a sudden increase in glucose influx rate (30). In that study, the change in the ATP pool is found to link to the purine metabolism pathways, which are linked to several conserved moieties; GTP, ATP, NAD, NADP, S-adenosylmethionine, and Coenzyme A. These findings suggest that cells could dynamically alter pool sizes associated with different parts of metabolism, limiting flux through some pathways, while allowing higher flux in others, and thereby shifting the metabolites from the latter to the former. This could provide a dynamic self-regulation and the pool sizes of key co-substrates could be seen as ‘tuning points’ controlling a more complex metabolic system. We thus propose further experimental analyses focusing on co-substrate pool sizes and turnover dynamics to understand and manipulate cell physiology.

## METHODS

### Model of a single reaction with co-substrate cycling

The metabolic system shown in Fig. 1A, comprises the following biochemical reactions:

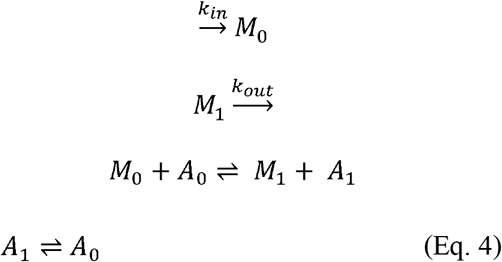

where metabolites are denoted by *M*_*i*_ and the different forms of the co-substrate are denoted by *A*_*i*_. We assume additional conversion between *A*_*1*_ and *A*_*0*_, mediated through other enzymatic reactions. The parameters *k*_*in*_, and *k*_*out*_ denote the in- and out-flux of *M*_*0*_ and *M*_*1*_ respectively, from and to other pathways or across cell boundary. The ordinary differential equations (ODEs) for the system shown in Eq. 4 (and Fig. 1A), using irreversible Michaelis-Menten enzyme kinetics would be:

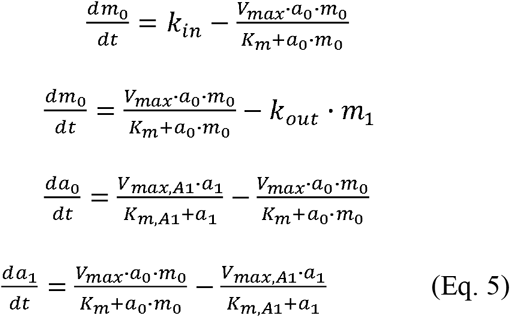

where *m*_*0*_ and *a*_*0*_ denote the concentrations of *M*_*0*_ and *A*_*0*_ respectively, *K*_*m*_ denotes a composite parameter of the Michaelis-Menten coefficients of the enzyme for its substrates, and *V*_*max*_ is the total enzyme concentration times its catalytic rate (i.e. *E* = *k*_*cat*_· *E*_*tot*_). We further have the conservation relation *a*_*0*_ + *a*_*1*_ = *A*_*tot*_, where *A*_*tot*_ is a constant. This assumption would be justified when influx of any form of the cycled metabolite into the system is independent of the rest of the metabolic system (see further discussion and analysis in *SI* section 2). The steady states of Eq. 5 can be found by setting the left side equal to zero and performing algebraic re-arrangements to isolate each of the variables (see *SI*). The resulting analytical expressions for steady state metabolite concentration are shown in Eq. 1, and in the *SI* for this model with reversible enzyme kinetics, as well as for other models.

### Symbolic and numerical computations

For all symbolic computations, utilised in finding steady state solutions and deriving mathematical conditions on rate parameters, we used the software Maple 2021, as well as theoretical results presented in (40). To run numerical simulations of select systems, we used Python packages with the standard solver functions. All numerical simulations were performed in the Python environment. The main model simulation files relating to Figures 4 and 5 are provided as *Supplementary Files 2* and *3*, while all remaining simulation and analysis scripts are made available at a dedicated Github page: https://github.com/OSS-Lab/CoSubstrateDynamics.

### Reaction fluxes and enzyme kinetic parameters

To support the model findings on co-substrate pools acting as a possible limitation on reaction fluxes, we analysed measured and FBA-derived flux data collated previously (31, 32). We focussed our analyses on reactions involving co-substrates only. We compared measured (or FBA-derived) fluxes to flux thresholds based on enzyme kinetics (i.e., first condition in Eq. 2). To calculate the latter, we used data on enzyme kinetics and levels as collated in (31), which is based on the BRENDA database (41) and proteomics-based measurements (42). We note that most available kinetic constants for enzymes have been obtained under *in vitro* conditions, which can be very different from those of the cytosol (43). When comparing flux levels against co-substrate pool sizes, we used the matching, measured pool-sizes from (32). All the data used in this analysis is provided in the *Supplementary File 1*, and through a dedicated Github page, which contains additional analysis scripts: https://github.com/OSS-Lab/CoSubstrateDynamics.

## Acknowledgements

We would like to thank Wenying Shou for constructive comments on an earlier version of this manuscript, and Dan Davidi for his help with datasets of reaction fluxes and enzyme abundances.

## SUPPLEMENTARY FILES

**Supplementary Information**. This file contains all of the supplementary figures, the mathematical analyses of the reaction systems and corresponding analytical solutions, and the descriptions of the simulated models.

**Supplementary File 1**. Enzyme kinetics, flux, metabolite concentration, and enzyme abundance data associated with flux analyses.

**Supplementary File 2**. Python implementation of branched pathway model, presented in Figure 4.

**Supplementary File 3**. Python implementation of connected pathway model, presented in Figure 5.

