## Supplementary information for "Co-substrate pools can constrain and regulate pathway fluxes in cell metabolism"

### for

**6 Different stoichiometries for co-substrate cycling along a single pathway – mimicking the case seen in upper glycolysis** **19**

**7 Co-substrate cycling along with metabolite cycling – mimicking the case seen in nitrogen assimilation** **21**

**8 Analysis of existing flux data against predicted limits** **24**

**9 Pathway branching into two pathways with independent co-substrates** **24**

**10 Independent pathways coupled by co-substrate cycling** **26**

#### 1 The mathematical approach and modeling setting

As explained in the main text, we are interested in understanding the effect of co-substrate cycling on the flux through metabolic pathways, such as those shown in Figure S1.

In this supplementary text, we use a generic co-substrate pair, denoted as $A_0$, $A_1$. We consider the synthesis or degradation of the co-substrate pair, or consider it as a conserved moiety, i.e. having a fixed total concentration. Our generic co-substrate pair, $A_0$ and $A_1$, can be taken as representing a specific co-substrate, such as NAD(H), but note that the mathematical analyses presented would be applicable to any co-substrate pair in natural metabolic pathways (as discussed in the main text).

For our analyses, we consider a generalised model of a linear metabolic pathway, as well as additional metabolic pathway structures. Throughout the presented analyses, we consider reactions to be either enzyme mediated or not, and when they are enzyme mediated, we consider them either to be **reversible** or **irreversible**. In the former case, the enzymatic conversions are shown as $M_{i-1} \rightleftharpoons M_i$, while in the latter case, they are shown as $M_{i-1} \longrightarrow M_i$. These notations do not show enzyme complexes explicitly, but we use enzymatic rates derived from reaction schemes accounting for enzyme complexes (see below).

In certain models, we consider some, or all, cycling reactions of the co-substrate to occur independently of the enzymatic reactions involved in the metabolic pathway, e.g. due to hydrolysis reactions. We refer to this type of recycling as **free conversion**, e.g. in the case of a generic co-substrate considered here, we have:

$$\mathrm{A}_0 \underset{k_6}{\overset{k_5}{\rightleftharpoons}} \mathrm{A}_1.$$

We talk about **irreversible co-substrate conversion**, if $k_5 = 0$ or $k_6 = 0$, that is, only conversion in one direction is considered. We talk about **no free co-substrate conversion**, if $k_5 = k_6 = 0$, that is, the co-substrate cycling is only related through the reactions in the metabolic pathway.

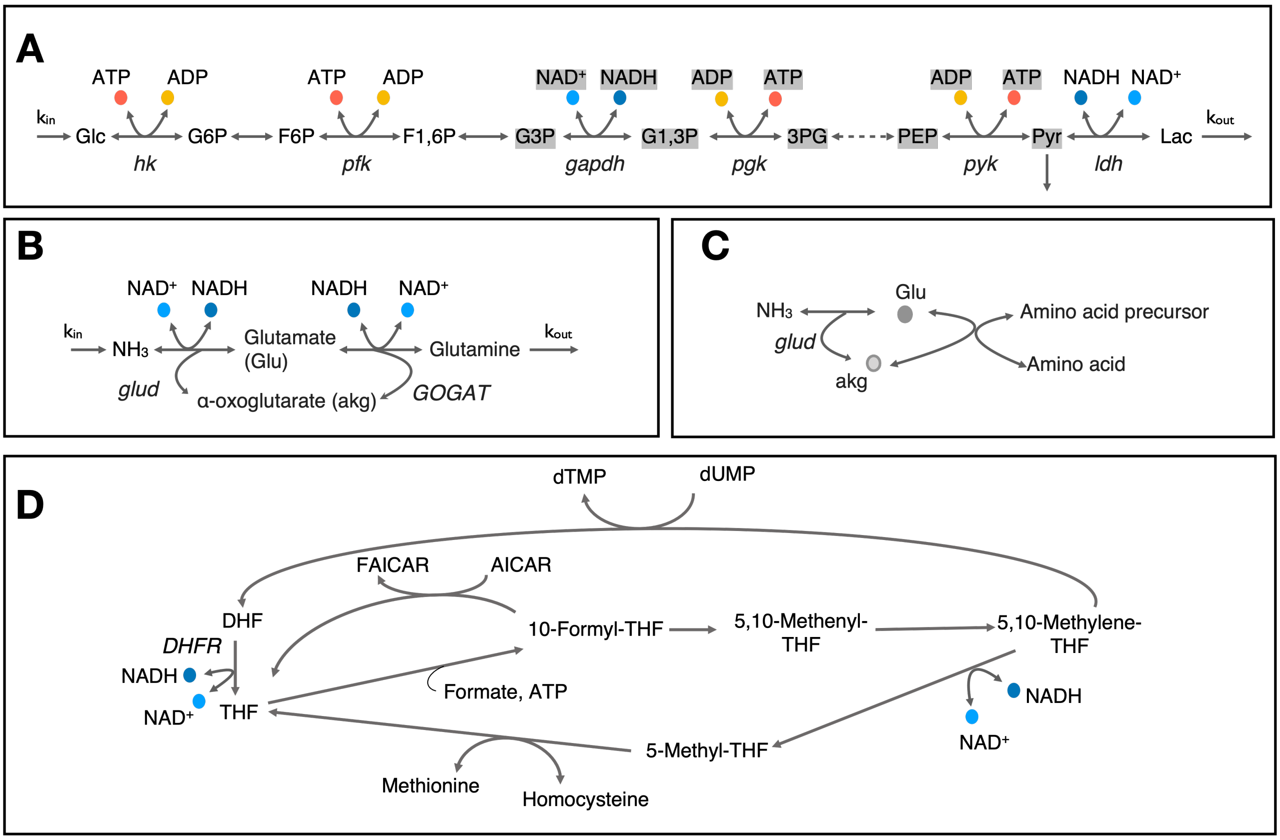

Figure S1: **(A)** Cartoon representation of upper glycolysis pathway. Note that stoichiometric balance across the pathway changes after F1,6P and metabolites and co-substrates highlighted with gray background have a stoichiometry of 2. **(B & C)** Cartoon representation of nitrogen assimilation via glutamate and involvement of glutamate cycling in amino acid biosynthesis. **(D)** Cartoon representation of metabolite cycles involved in one-carbon metabolism around tetrahydrofolate. Enzyme and metabolite name abbreviations are: Glc – Glucose, G6P – Glucose-6-phosphate, F6P – Fructose-6-phosphate, F1,6P – Fructose-1,6-biphosphate, G3P – glyceraldehyde-3-phosphate, G1,3P – 1,3-biphospho-D-glycerate, 3PG – 3-phospho-D-glycerate, PEP – Phosphoenolpyruvate, Pyr – Pyruvate, Lac – Lactate, DHF – Dihydrofolate, THF - Tetrahydrofolate, AICAR - 5-amino-4-imidazolecarboxamide ribotide, FAICAR - 5'-phosphoribosyl-formamido-carboxamide, hk – hexokinase, pfk – phosphofructokinase, gapdh – glyceraldehyde-3-phosphate dehydrogenase, pgk – phosphoglycerate kinase, pyk – phophoenolpyruvate kinase, ldh – lactate dehydrogenase, glud – glutamate dehydrogenase, GOGAT – glutamate synthase, DHFR – dihydrofolate reducatase.

#### 1.1 Enzyme kinetics

Each metabolic pathway is modelled using either Michaelis-Menten (irreversible case) or Haldane (reversible case) enzyme kinetics, for the individual reactions it comprises. The general kinetics can be expressed as follows, where we let $a_0, a_1$ denote respectively the concentrations of the co-substrate pair $A_0$ and $A_1$, and $m_i$ to denote the concentration of $M_i$, the $i$-th metabolite in the pathway.

In the case of a reversible, enzymatic reaction involving a co-substrate and assuming simultaneous binding of both substrates to the enzyme, we have the following reaction scheme:

$$\mathrm{M}_{i-1} + \mathrm{A}_0 + E \xrightleftharpoons[k_{\mathrm{off}}]{k_{\mathrm{on}}} E\mathrm{M}_{i-1}\mathrm{A}_0 \xrightleftharpoons[k_{\mathrm{off},1}]{k_{\mathrm{on},1}} E\mathrm{M}_i\mathrm{A}_1 \xrightleftharpoons[k_{\mathrm{off},2}]{k_{\mathrm{on},2}} \mathrm{M}_i + \mathrm{A}_1 + E.$$

For this reversible reaction scheme, the rate of production of $M_i$ takes the form

$$v = \frac{E_i L_{i-1}\, m_{i-1} a_0 - F_i K_i\, m_i a_1}{K_i L_{i-1} + K_i\, m_i a_1 + L_{i-1}\, m_{i-1} a_0}.$$

Likewise, for the reversible enzymatic conversion $M_{i-1} \rightleftharpoons M_i$, we have the following reaction scheme:

$$\mathrm{M}_{i-1} + E \xrightleftharpoons[k_{\mathrm{off}}]{k_{\mathrm{on}}} E\mathrm{M}_{i-1} \xrightleftharpoons[k_{\mathrm{off},1}]{k_{\mathrm{on},1}} E\mathrm{M}_i \xrightleftharpoons[k_{\mathrm{off},2}]{k_{\mathrm{on},2}} \mathrm{M}_i + E.$$

The rate of production of $M_i$ is given by

$$v = \frac{E_i L_{i-1}\, m_{i-1} - F_i K_i\, m_i}{K_i L_{i-1} + K_i\, m_i + L_{i-1}\, m_{i-1}}.$$

In both of these reversible rate equations, the parameters $K$ and $L$ are equivalent to the Haldane coefficients $K_S$ and $K_P$, respectively and are given by

$$K_i = \frac{k_{\mathrm{on},1}k_{\mathrm{on},2} + k_{\mathrm{off}}k_{\mathrm{on},2} + k_{\mathrm{off}}k_{\mathrm{off},1}}{k_{\mathrm{on}}(k_{\mathrm{on},2} + k_{\mathrm{off},1} + k_{\mathrm{on},1})} \quad \text{and} \quad L_{i-1} = \frac{k_{\mathrm{on},1}k_{\mathrm{on},2} + k_{\mathrm{off}}k_{\mathrm{on},2} + k_{\mathrm{off}}k_{\mathrm{off},1}}{k_{\mathrm{off},2}(k_{\mathrm{off},1} + k_{\mathrm{on},1} + k_{\mathrm{off}})}. \tag{1.1}$$

When there are two substrates that take part in the reaction, the $k_{\mathrm{on}}$ and $k_{\mathrm{off},2}$ parameters are composite parameters composed of two binding coefficients, one for each substrate. This does not affect the derivations, so for convenience we use $K_S$ and $K_P$.

The parameters $E$ and $F$ correspond to the Haldane coefficients $k_{\mathrm{cat}+}$ and $k_{\mathrm{cat}-}$, multiplied by the total enzyme concentration (denoted $E_{\mathrm{tot}}$, below), and are given by

$$E_i = E_{\mathrm{tot}} \frac{k_{\mathrm{on},1}k_{\mathrm{on},2}}{k_{\mathrm{on},2} + k_{\mathrm{off},1} + k_{\mathrm{on},1}} \quad \text{and} \quad F_i = E_{\mathrm{tot}} \frac{k_{\mathrm{off}}k_{\mathrm{off},1}}{k_{\mathrm{off}} + k_{\mathrm{off},1} + k_{\mathrm{on},1}}. \tag{1.2}$$

For the irreversible enzymatic reaction, the reaction schemes simplify to:

$$\mathrm{M}_{i-1} + \mathrm{A}_0 + E \xrightleftharpoons[k_{\mathrm{off}}]{k_{\mathrm{on}}} E\mathrm{M}_{i-1}\mathrm{A}_0 \xrightarrow{k_{\mathrm{cat}}} \mathrm{M}_i + \mathrm{A}_1 + E.$$

$$\mathrm{M}_{i-1} + E \xrightleftharpoons[k_{\mathrm{off}}]{k_{\mathrm{on}}} E\mathrm{M}_{i-1} \xrightarrow{k_{\mathrm{cat}}} \mathrm{M}_i + E.$$

And the rate of production for the two cases are given by

$$v = \frac{E_i\, m_{i-1} a_0}{K_i + m_{i-1} a_0}, \qquad \text{and} \qquad v = \frac{E_i\, m_{i-1}}{K_i + m_{i-1}},$$

where $E_i$ is the catalytic rate coefficient of the $i$-th enzyme multiplied by its total concentration, and $K_i$ is its Michaelis-Menten coefficient. Again, when there are two substrates, the $k_{\mathrm{on}}$ parameter is a composite parameter composed of two binding coefficients, one for each substrate. As in the reversible case, this does not affect the derivations, so we use $K_i$ for convenience. Influx and outflux follow non-enzymatic reaction kinetics, with reaction rate constants as indicated by the labels of the reactions.

##### 1.2 Analytical approach

Our mathematical analysis is primarily concerned with finding conditions on the kinetic parameters, if any, that imply that the system has a positive steady state. This is different than system reduction, e.g. as done in the analyses leading to Michaelis-Menten kinetics. Our analysis distinctively solves the entire system for steady states and determines conditions on kinetic parameters to satisfy the steady state equations.

Thus, for each of the metabolic pathway motifs we consider, we build the ODEs defining the rates of change of variables, find the conservation laws among variables, and consider a system of equations whose solutions are the steady states of the ODEs constrained by the conservation laws. We then follow one of two strategies. We first attempt to solve all equations for all concentrations. For some systems, we readily get an expression in terms of the parameters of the system. For other systems, this approach is not possible. In this case, using all equations in the system but one, we solve for the steady states of all concentrations but one. This gives all concentrations in terms of the remaining concentration, say $x$. Plugging these expressions in the remaining equation of the system, we obtain a final equation whose solutions characterize the steady states of the system. We need then to study when the solutions obtained this way are positive.

We are also interested in proving if a given system has a positive steady state for all parameter combinations, and that this steady state is stable. When there is one positive steady state, we find the Hurwtiz determinants associated with the characteristic polynomial of the Jacobian of the system of ODEs, evaluated at the steady state. If these are all positive, then the steady state is asymptotically stable [1].

To decide on the existence of a steady state, throughout the analysis, we will use repeatedly the following lemma, which is a consequence of the well-known Descartes' rule of signs.

**Lemma 1.** *Let $p(x)$ be a univariate polynomial of degree two, with negative leading term. If at some value $T$, we have $p(T) > 0$, then $p$ has a root in the interval $(0, T)$ if and only if the independent term of $p$ is negative.*

*Proof.* The Descartes' rule of signs establishes that the number of positive roots of a polynomial cannot exceed the number $\tau$ of sign changes in the sequence of coefficients ignoring zero coefficients, and the difference between $\tau$ and the number of positive roots is an even number. As the polynomial $p$ in the statement attains positive values, it must have some positive coefficient. Furthermore, as the degree two polynomial has negative leading term, the sequence of the sign of terms (when terms are ordered from lowest exponent to highest) is one of the following $++-$, $+--$, $-+-$, $+0-$, $0+-$.

If the independent term is positive or zero, then the sign sequence is one of $++-$, $+--$, $+0-$, $0+-$. In this case, there is one sign change in the sequence, and it follows that the polynomial has exactly one positive root. As $p(0) > 0$, $p(T) > 0$ and $p$ becomes negative as $x$ goes to $+\infty$, the root must be in the interval $(T, +\infty)$.

If the independent term is negative, then the sign sequence is $-+-$. The polynomial is negative both at 0 and at $+\infty$. As $p(T) > 0$, there must be a positive root in $(0, T)$ and one in $(T, +\infty)$, and there cannot be more by the Descartes' rule of signs. From this, the statement of the lemma follows. □

#### 2 Considering co-substrate pool size

We first consider cycling of a generic co-substrate $A_0$ and $A_1$, with biosynthesis and degradation of both forms.

$$0 \underset{k_2}{\overset{k_1}{\rightleftharpoons}} A_0 \qquad 0 \underset{k_4}{\overset{k_3}{\rightleftharpoons}} A_1 \qquad A_0 \underset{k_6}{\overset{k_5}{\rightleftharpoons}} A_1.$$

We suppose that the biosynthesis occurs at a constant rate, while degradation and cycling are proportional to the concentration of the relevant chemical species. Writing $a_0 = [A_0]$ and $a_1 = [A_1]$, the differential equations for these concentrations are:

$$\frac{da_0}{dt} = k_1 - (k_2 + k_5)a_0 + k_6 a_1$$
$$\frac{da_1}{dt} = k_3 - (k_4 + k_6)a_1 + k_5 a_0.$$

Since all the terms are linear or constant, the steady state values are the solutions of the linear equation:

$$\begin{pmatrix} -(k_2 + k_5) & k_6 \\ k_5 & -(k_4 + k_6) \end{pmatrix} \begin{pmatrix} a_0 \\ a_1 \end{pmatrix} = \begin{pmatrix} -k_1 \\ -k_3 \end{pmatrix}.$$

The steady states are then found to be:

$$a_0 = \frac{k_1 k_4 + k_1 k_6 + k_3 k_6}{k_4 k_5 + k_2 k_4 + k_2 k_6}, \qquad a_1 = \frac{k_2 k_3 + k_3 k_5 + k_1 k_5}{k_4 k_5 + k_2 k_4 + k_2 k_6}.$$

If we consider the case where the synthesis and degradation rates of the different forms of the co-substrate (i.e. cycled metabolite) are the same, i.e. $k_1 = k_3 = k_s$ and $k_2 = k_4 = k_d$, these equations simplify to:

$$a_0 = \frac{k_s(k_d + 2k_6)}{k_d(k_d + k_5 + k_6)}, \qquad a_1 = \frac{k_s(k_d + 2k_5)}{k_d(k_d + k_5 + k_6)},$$

and the eigenvalues of the Jacobian of the system evaluated at this steady state are always real and negative. When $k_d$ is sufficiently small compared to co-substrate conversion rates, it can be safely neglected in the brackets, resulting in the expression of steady state formulas as:

$$a_0 = \frac{2k_6 k_s / k_d}{k_5 + k_6}, \qquad a_1 = \frac{2k_5 k_s / k_d}{k_5 + k_6},$$

We can compare the above expressions with those obtained from the case, where we assume a constant pool size of the cycled metabolite (i.e. $k_1 = k_2 = k_3 = k_4 = 0$). In that case, the steady states are $a_0 = Tk_6/(k_5 + k_6)$ and $a_1 = Tk_5/(k_5 + k_6)$, where $T$ is the total pool size. Thus, under the limit of degradation rates being much smaller than conversion rates, the two cases will be identical and co-substrates will act as a conserved moiety for the rest of the metabolic system.

If we now assume that the cycling of co-substrates is an enzymatic reaction and make the same simplifying assumptions as above that $k_1 = k_3 = k_s$ and $k_2 = k_4 = k_d$, the ODEs for the system are:

$$\frac{da_0}{dt} = k_s - k_d a_0 - \frac{a_0 E_a L_a - a_1 F_a K_a}{a_0 L_a + a_1 K_a + K_a L_a},$$
$$\frac{da_1}{dt} = k_s - k_d a_1 + \frac{a_0 E_a L_a - a_1 F_a K_a}{a_0 L_a + a_1 K_a + K_a L_a}.$$

The only real and positive steady state is now found to be:

$$a_0 = \Big(K_a L_a k_d + K_a(F_a + 3k_s) + L_a(E_a - k_s)$$
$$- \sqrt{-4K_a k_s(K_a - L_a)(2(F_a + k_s) + k_d L_a) + (F_a K_a + 3K_a k_s + E_a L_a + K_a k_d L_a - k_s L_a)^2}\Big)/(2k_d(K_a - L_a)),$$
$$a_1 = \Big( - K_a L_a k_d - K_a(F_a - 3k_s) - L_a(E_a + 3k_s)$$
$$+ \sqrt{-4K_a k_s(K_a - L_a)(2(F_a + k_s) + k_d L_a) + (F_a K_a + 3K_a k_s + E_a L_a + K_a k_d L_a - k_s L_a)^2}\Big)/(2k_d(K_a - L_a)).$$

This is stable as long as all parameters are positive. Note that in the case of $K_a = L_a$, the steady state solutions converge to a real number less than infinity by l'Hopital's Rule. Also, note that the sum $a_0 + a_1$ is constant as in the non-enzymatic case presented above. Thus, whether the metabolite cycling is considered as a non-enzymatic or enzymatic (i.e. following Michaelis-Menten kinetics) reaction, the co-substrates will act as a conserved moiety for the rest of the metabolic system in both cases.

### 3 Single reaction models

In this section we derive results for a single reaction of two metabolites, involving co-substrate cycling or not, as presented in the main text.

##### 3.1 Enzymatic reaction with co-substrate cycling

We first reconsider the case where all reactions, including the off-pathway cycling, are enzymatic (hence are modelled with Michaelis-Menten kinetics). The reactions are:

$$0 \xrightarrow{k_{\text{in}}} M_0 \qquad M_0 + \text{A}_0 \rightleftharpoons M_1 + \text{A}_1 \qquad M_1 \xrightarrow{k_{\text{out}}} 0 \qquad \text{A}_0 \rightleftharpoons \text{A}_1.$$

This corresponds to the motif depicted in Fig. 1A of the main text, and the resulting ODEs are:

$$\begin{aligned}
\frac{dm_0}{dt} &= k_{\text{in}} - \frac{E_1 L_0 m_0 a_0 - F_1 K_1 m_1 a_1}{K_1 m_1 a_1 + L_0 a_0 m_0 + K_1 L_0}\\
\frac{dm_1}{dt} &= \frac{E_1 L_0 m_0 a_0 - F_1 K_1 m_1 a_1}{K_1 m_1 a_1 + L_0 a_0 m_0 + K_1 L_0} - k_{\text{out}} m_1\\
\frac{da_0}{dt} &= -\frac{E_1 L_0 m_0 a_0 - F_1 K_1 m_1 a_1}{K_1 m_1 a_1 + L_0 a_0 m_0 + K_1 L_0} - \frac{E_a L_a a_0 - F_a K_a a_1}{K_a a_1 + L_a a_0 + K_a L_a}\\
\frac{da_1}{dt} &= \frac{E_1 L_0 m_0 a_0 - F_1 K_1 m_1 a_1}{K_1 m_1 a_1 + L_0 a_0 m_0 + K_1 L_0} + \frac{E_a L_a a_0 - F_a K_a a_1}{K_a a_1 + L_a a_0 + K_a L_a}.
\end{aligned}$$

This ODE system has one conservation law, namely the sum of $a_0$ and $a_1$ is constant:

$$a_0 + a_1 = A_{\text{tot}}.$$

The steady states of the system are:

$$\begin{aligned}
m_0 &= \frac{K_1 k_{\text{in}}}{(E_1 - k_{\text{in}})\,(F_a A_{\text{tot}} - k_{\text{in}}(L_a + A_{\text{tot}}))}\alpha,\\
a_0 &= \frac{K_a\,(F_a A_{\text{tot}} - k_{\text{in}}(L_a + A_{\text{tot}}))}{K_a(F_a - k_{\text{in}}) + L_a(E_a + k_{\text{in}})},\\
a_1 &= \frac{L_a\,(K_a k_{\text{in}} + T(E_a + k_{\text{in}}))}{K_a(F_a - k_{\text{in}}) + L_a(E_a + k_{\text{in}})}.
\end{aligned}$$

where, we introduced the composite parameter alpha, as follows:

$$\alpha = \frac{k_{\text{out}} L_0 \big(K_a(F_a - k_{\text{in}}) + L_a(E_a + k_{\text{in}})\big) + (F_1 + k_{\text{in}}) L_a (K_a k_{\text{in}} + k_{\text{in}} A_{\text{tot}} + E_a A_{\text{tot}})}{K_a L_0 k_{\text{out}}}. \tag{3.1}$$

For the steady state equations given above to be positive, the following conditions must be satisfied:

$$k_{\text{in}} < E_1 \quad \text{and} \quad k_{\text{in}} < \frac{F_a A_{\text{tot}}}{L_a + A_{\text{tot}}}. \tag{3.2}$$

Note that as $\frac{F_a A_{\text{tot}}}{L_a + A_{\text{tot}}} < F_a$, the second condition readily implies $k_{\text{in}} < F_a$ and $\alpha$ is positive. When there is a positive steady state, then it is asymptotically stable.

If the **main pathway is irreversible**, but the co-substrate reaction is still reversible, the ODEs describing the system dynamics are:

$$\begin{aligned}
\frac{dm_0}{dt} &= k_{\text{in}} - \frac{E_1 m_0 a_0}{K_1 + m_0 a_0},\\
\frac{dm_1}{dt} &= \frac{E_1 m_0 a_0}{K_1 + m_0 a_0} - k_{\text{out}} m_1,\\
\frac{da_0}{dt} &= -\frac{E_1 m_0 a_0}{K_1 + m_0 a_0} - \frac{E_a L_a a_0 - F_a K_a a_1}{K_a a_1 + L_a a_0 + K_a L_a},\\
\frac{da_1}{dt} &= \frac{E_1 m_0 a_0}{K_1 + m_0 a_0} + \frac{E_a L_a a_0 - F_a K_a a_1}{K_a a_1 + L_a a_0 + K_a L_a}.
\end{aligned}$$

The steady state of the system is:

$$\begin{aligned}
m_0 &= \frac{K_1 k_{\text{in}}}{(E_1 - k_{\text{in}})(F_a A_{\text{tot}} - k_{\text{in}}(L_a + A_{\text{tot}}))}\beta,\\
m_1 &= \frac{k_{\text{in}}}{k_{\text{out}}},\\
a_0 &= \frac{K_a(F_a A_{\text{tot}} - k_{\text{in}}(L_a + A_{\text{tot}}))}{K_a\,(F_a - k_{\text{in}}) + L_a\,(E_a + k_{\text{in}})},\\
a_1 &= \frac{L_a\big(K_a k_{\text{in}} + A_{\text{tot}}(E_a + k_{\text{in}})\big)}{K_a\,(F_a - k_{\text{in}}) + L_a\,(E_a + k_{\text{in}})}.
\end{aligned}$$

Where the composite parameter beta is defined as:

$$\beta = \frac{K_a\left(F_a - k_{\text{in}}\right) + L_a\left(E_a + k_{\text{in}}\right)}{K_a}. \tag{3.3}$$

For this to be positive the same conditions as in the reversible case must be satisfied:

$$k_{\text{in}} < E_1 \quad \text{and} \quad k_{\text{in}} < \frac{F_a A_{\text{tot}}}{L_a + A_{\text{tot}}}. \tag{3.4}$$

When these are satisfied, $\beta$ is positive, and the positive steady state is asymptotically stable.

If the **main pathway is irreversible**, and the co-substrate reaction only flows from $A_1$ to $A_0$, the ODEs describing the system dynamics are:

$$\begin{aligned}
\frac{dm_0}{dt} &= k_{\text{in}} - \frac{E_1 m_0 a_0}{K_1 + m_0 a_0},\\
\frac{dm_1}{dt} &= \frac{E_1 m_0 a_0}{K_1 + m_0 a_0} - k_{\text{out}} m_1,\\
\frac{da_0}{dt} &= -\frac{E_1 m_0 a_0}{K_1 + m_0 a_0} + \frac{F_a a_1}{L_a + a_1},\\
\frac{da_1}{dt} &= \frac{E_1 m_0 a_0}{K_1 + m_0 a_0} - \frac{F_a a_1}{L_a + a_1}.
\end{aligned}$$

The steady state of the system is:

$$\begin{aligned}
m_0 &= \frac{K_1 k_{\text{in}}\left(F_a - k_{\text{in}}\right)}{(E_1 - k_{\text{in}})(F_a A_{\text{tot}} - k_{\text{in}}(L_a + A_{\text{tot}}))},\\
m_1 &= \frac{k_{\text{in}}}{k_{\text{out}}},\\
a_0 &= A_{\text{tot}} - \frac{k_{\text{in}} L_a}{F_a - k_{\text{in}}},\\
a_1 &= \frac{k_{\text{in}} L_a}{F_a - k_{\text{in}}}.
\end{aligned}$$

For these steady states to be positive the same conditions as in the other two cases must be satisfied:

$$k_{\text{in}} < E_1 \quad \text{and} \quad k_{\text{in}} < \frac{F_a A_{\text{tot}}}{L_a + A_{\text{tot}}}. \tag{3.5}$$

When these are satisfied, the positive steady state is asymptotically stable.

#### 3.2 Enzymatic reaction with co-substrate cycling, biosynthesis and degradation

To extend the previous analysis we consider the same simple scenario where a pathway involves a single co-substrate consuming reaction and a back reaction re-generating the co-substrate, with the addition of synthesis and degradation of the co-substrate forms. This system is comprised of the following reactions:

$$\begin{gathered}
0 \xrightarrow{k_{\text{in}}} M_0 \qquad M_0 + \mathrm{A}_0 \rightleftharpoons M_1 + \mathrm{A}_1 \qquad M_1 \xrightarrow{k_{\text{out}}} 0\\
\mathrm{A}_0 \underset{k_6}{\overset{k_5}{\rightleftharpoons}} \mathrm{A}_1 \qquad 0 \underset{k_8}{\overset{k_7}{\rightleftharpoons}} \mathrm{A}_0 \qquad 0 \underset{k_{10}}{\overset{k_9}{\rightleftharpoons}} \mathrm{A}_1.
\end{gathered}$$

The resulting system of ODEs is:

$$\begin{aligned}
\frac{dm_0}{dt} &= k_{\text{in}} - \frac{E_1 L_0 m_0 a_0 - F_1 K_1 m_1 a_1}{K_1 m_1 a_1 + L_0 a_0 m_0 + K_1 L_0}\\
\frac{dm_1}{dt} &= \frac{E_1 L_0 m_0 a_0 - F_1 K_1 m_1 a_1}{K_1 m_1 a_1 + L_0 a_0 m_0 + K_1 L_0} - k_{\text{out}} m_1\\
\frac{da_0}{dt} &= -\frac{E_1 L_0 m_0 a_0 - F_1 K_1 m_1 a_1}{K_1 m_1 a_1 + L_0 a_0 m_0 + K_1 L_0} - (k_5 + k_8) a_0 + k_6 a_1 + k_7\\
\frac{da_1}{dt} &= \frac{E_1 L_0 m_0 a_0 - F_1 K_1 m_1 a_1}{K_1 m_1 a_1 + L_0 a_0 m_0 + K_1 L_0} + k_5 a_0 - (k_6 + k_{10}) a_1 + k_9.
\end{aligned}$$

The steady state concentrations are then given by:

$$
\begin{aligned}
a_0 &= \frac{k_{10}(k_7 - k_{\text{in}}) + k_6(k_7 + k_9)}{k_{10}k_5 + k_{10}k_8 + k_6k_8} \\
a_1 &= \frac{k_8(k_9 + k_{\text{in}}) + k_5(k_7 + k_9)}{k_{10}k_5 + k_{10}k_8 + k_6k_8} \\
m_0 &= \frac{K_1k_{\text{in}}\left((F_1 + k_{\text{in}})(k_8(k_9 + k_{\text{in}}) + k_5(k_7 + k_9)) + k_{\text{out}}L_0(k_{10}k_5 + k_{10}k_8 + k_6k_8)\right)}{L_0k_{\text{out}}\left(E_1 - k_{\text{in}}\right)\left(k_{10}(k_7 - k_{\text{in}}) + k_6(k_7 + k_9)\right)} \\
m_1 &= \frac{k_{\text{in}}}{k_{\text{out}}}.
\end{aligned}
$$

These expressions are positive if and only if

$$
k_{\text{in}} < k_7 + \frac{k_6}{k_{10}}(k_7 + k_9) \qquad \text{and} \qquad k_{\text{in}} < E_1.
$$

If the **main path is irreversible**, similarly to the previous section, the ODEs describing the system's dynamics, are:

$$
\begin{aligned}
\frac{dm_0}{dt} &= k_{\text{in}} - \frac{E_1m_0a_0}{K_1 + m_0a_0} \\
\frac{dm_1}{dt} &= \frac{E_1m_0a_0}{K_1 + m_0a_0} - k_{\text{out}}m_1 \\
\frac{da_0}{dt} &= -\frac{E_1m_0a_0}{K_1 + m_0a_0} - (k_5 + k_8)a_0 + k_6a_1 + k_7 \\
\frac{da_1}{dt} &= \frac{E_1m_0a_0}{K_1 + m_0a_0} + k_5a_0 - (k_6 + k_{10})a_1 + k_9.
\end{aligned}
$$

The steady state in this case is:

$$
\begin{aligned}
a_0 &= \frac{k_{10}(k_7 - k_{\text{in}}) + k_6(k_7 + k_9)}{k_{10}k_5 + k_{10}k_8 + k_8k_6} \\
a_1 &= \frac{k_8(k_9 + k_{\text{in}}) + k_5(k_7 + k_9)}{k_{10}k_5 + k_{10}k_8 + k_8k_6} \\
m_0 &= \frac{K_1k_{\text{in}}(k_{10}k_5 + k_{10}k_8 + k_8k_6)}{(E_1 - k_{\text{in}})(k_{10}(k_7 - k_{\text{in}}) + k_6(k_7 + k_9))} \\
m_1 &= \frac{k_{\text{in}}}{k_{\text{out}}}.
\end{aligned}
$$

These expressions are positive if and only if

$$
k_{\text{in}} < k_7 + \frac{k_6}{k_{10}}(k_7 + k_9) \qquad \text{and} \qquad k_{\text{in}} < E_1.
$$

Comparing these results with those of Subsection 3.1, we see some similar themes. Firstly, the total amount of co-substrate $a_0 + a_1$ at steady state is a constant, even when it is not explicitly considred to be a conserved moiety. Secondly, one of the conditions for a positive steady state is $k_{\text{in}} < E_1$, and any other conditions take the form $k_{\text{in}} < f$, where $f$ is a function of the parameters controlling synthesis, degradation and the turnover of the co-substrate. Thus, in the analysis that follows, we only consider the case of conserved co-substrate, as adding synthesis and degradation only affects the quantitative results, not the qualitative behaviour.

##### 3.3 Enzymatic reaction without co-substrate cycling

Considering an enzymatic reaction without co-substrates, the reactions are:

$$
0 \xrightarrow{k_{\text{in}}} M_0 \qquad\qquad M_0 \underset{k_2}{\overset{k_1}{\rightleftharpoons}} M_1 \qquad\qquad M_1 \xrightarrow{k_{\text{out}}} 0.
$$

The resulting system of ODEs is:

$$\begin{aligned}\frac{dm_0}{dt} &= k_{\text{in}} - \frac{E_1L_0m_0 - F_1K_1m_1}{K_1m_1 + L_0m_0 + K_1L_0}\\ \frac{dm_1}{dt} &= \frac{E_1L_0m_0 - F_1K_1m_1}{K_1m_1 + L_0m_0 + K_1L_0} - k_{\text{out}}m_1.\end{aligned}$$

In this case there is no conservation law, and the steady states of the system are:

$$m_0 = \frac{K_1k_{\text{in}}\,(F_1 + k_{\text{in}} + k_{\text{out}}L_0)}{k_{\text{out}}L_0\,(E_1 - k_{\text{in}})}, \qquad m_1 = \frac{k_{\text{in}}}{k_{\text{out}}}.$$

These expressions are positive if and only if $k_{\text{in}} < E_1$ and the steady state is asymptotically stable when this holds.

If the **main path is irreversible**, the ODEs describing system dynamics are:

$$\begin{aligned}\frac{dm_0}{dt} &= k_{\text{in}} - \frac{E_1m_0}{K_1 + m_0}\\ \frac{dm_1}{dt} &= \frac{E_1m_0}{K_1 + m_0} - k_{\text{out}}m_1,\end{aligned}$$

and the steady state is:

$$m_0 = \frac{K_1k_{\text{in}}}{E_1 - k_{\text{in}}}, \qquad m_1 = \frac{k_{\text{in}}}{k_{\text{out}}}.$$

Again, expressions are positive if and only if $k_{\text{in}} < E_1$ and the steady state is asymptotically stable when this holds.

From this we see that the condition $k_{\text{in}} < E_1$ for stability in the enzymatic system with co-substrate cycling is a result of the reaction being enzymatic.

#### 3.4 Non-Enzymatic reaction with co-substrate cycling

Consider a theoretical case of non-enzymatic reactions for the main reaction:

$$0 \xrightarrow{k_{\text{in}}} M_0 \qquad M_0 + \mathrm{A}_0 \underset{k_2}{\overset{k_1}{\rightleftharpoons}} M_1 + \mathrm{A}_1 \qquad M_1 \xrightarrow{k_{\text{out}}} 0 \qquad \mathrm{A}_0 \underset{k_4}{\overset{k_3}{\rightleftharpoons}} \mathrm{A}_1.$$

The resulting system of ODEs, describing system dynamics, is:

$$\begin{aligned}\frac{dm_0}{dt} &= k_{\text{in}} - k_1m_0a_0 + k_2m_1a_1\\ \frac{dm_1}{dt} &= k_1m_0a_0 - k_2m_1a_1 - k_{\text{out}}m_1\\ \frac{da_0}{dt} &= -k_1m_0a_0 + k_2a_1m_1 - k_3a_0 + k_4a_1\\ \frac{da_1}{dt} &= k_1m_0a_0 - k_2m_1a_1 + k_3a_0 - k_4a_1\end{aligned}$$

There is the conservation law

$$a_0 + a_1 = A_{\text{tot}}.$$

The steady state of the system is:

$$\begin{aligned}m_0 &= \frac{k_{\text{in}}(k_2(k_{\text{in}} + k_3A_{\text{tot}}) + k_{\text{out}}(k_3 + k_4))}{k_1k_{\text{out}}(k_4A_{\text{tot}} - k_{\text{in}})}\\ m_1 &= \frac{k_{\text{in}}}{k_{\text{out}}},\\ a_0 &= \frac{k_4A_{\text{tot}} - k_{\text{in}}}{k_3 + k_4},\\ a_1 &= \frac{k_3A_{\text{tot}} + k_{\text{in}}}{k_3 + k_4}.\end{aligned}$$

These expressions are positive if and only if $k_{\text{in}} < k_4A_{\text{tot}}$, and when this is satisfied the steady state is asymptotically stable.

If the **system reactions are irreversible**, the ODEs describing the system dynamics are:

$$\begin{aligned}
\frac{dm_0}{dt} &= k_{\text{in}} - k_1 m_0 a_0 \\
\frac{dm_1}{dt} &= k_1 m_0 a_0 - k_{\text{out}} m_1 \\
\frac{da_0}{dt} &= -k_1 m_0 a_0 + k_4 a_1 \\
\frac{da_1}{dt} &= k_1 m_0 a_0 - k_4 a_1
\end{aligned}$$

and the steady state is:

$$\begin{aligned}
m_0 &= \frac{k_4 k_{\text{in}}}{k_1 (k_4 A_{\text{tot}} - k_{\text{in}})}, \\
m_1 &= \frac{k_{\text{in}}}{k_{\text{out}}}, \\
a_0 &= \frac{k_4 A_{\text{tot}} - k_{\text{in}}}{k_4}, \\
a_1 &= \frac{k_{\text{in}}}{k_4}.
\end{aligned}$$

As in the reversible case, these expressions are positive when $k_{\text{in}} < k_4 A_{\text{tot}}$ and the steady state is asymptotically stable when this holds.

Hence, we see that introducing co-substrate cycling always introduces a new constraint into the system, even in this simple case where the cycling is mass-action based and not enzymatic.

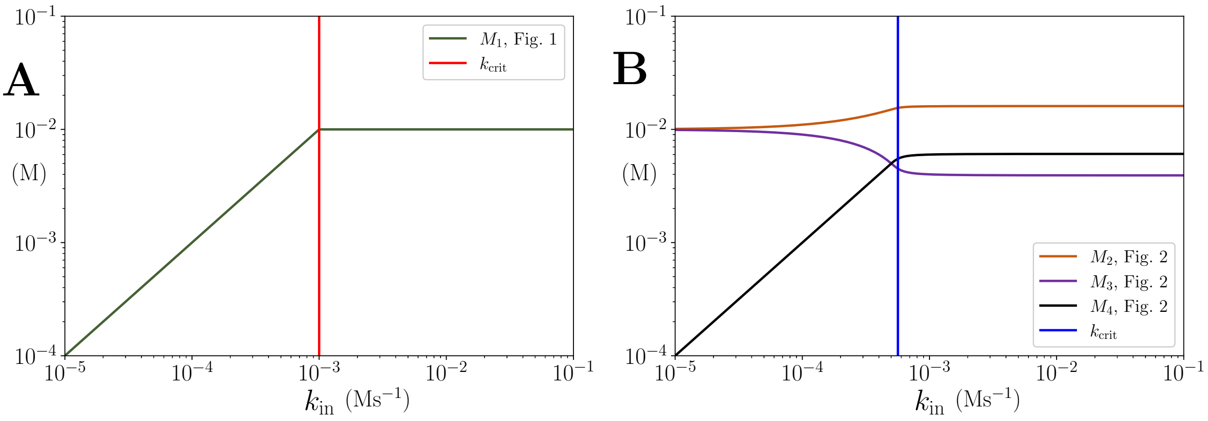

Figure S2: Concentrations of non-accumulating metabolites vs $k_{\text{in}}$ in the models presented in Figures 1 and 2 of the main text. **(A)** Concentration of non-accumulating metabolite $M_1$ from Figure 1. Red line shows the critical $k_{\text{in}}$ value, after which the concentration of $M_1$ remains constant **(B)** Concentrations of non-accumulating metabolites from the models presented in Figure 2. Blue line shows the threshold value for $k_{\text{in}}$. Once the threshold value for $k_{\text{in}}$ is reached the concentrations of the non-accumulating metabolites do not change. $A_{\text{tot}} = 10^{-1}$ in panel (A) and $10^{-6}$ in panel (B). All other parameters are the same as their counterparts in the main text.

### 4 Linear, arbitrary length, pathway model with co-substrate cycling

We consider next a linear, generic pathway of $n+1$ metabolites, and comprising the following reaction mechanism:

$$\begin{array}{llll}
0 \xrightarrow{k_{\text{in}}} M_0 & M_0 + \text{A}_0 \rightleftharpoons M_1 + \text{A}_1 & & M_{n-1} + \text{A}_1 \rightleftharpoons M_n + \text{A}_0 \\
M_1 \rightleftharpoons M_2 \;\cdots\; M_{i-1} \rightleftharpoons M_i \rightleftharpoons M_{i+1} & \cdots \rightleftharpoons M_{n-1} & & M_n \xrightarrow{k_{\text{out}}} 0.
\end{array} \tag{4.1}$$

#### 4.1 Steady states of the linear pathway model. The case with $n = 3$

We first consider the dynamics of model (4.1) with $n = 3$, as this is the first case where the system makes biochemical sense. This system has the form:

$$\begin{aligned} 0 \xrightarrow{k_{\text{in}}} M_0 \qquad & M_0 + \mathrm{A}_0 \rightleftharpoons M_1 + \mathrm{A}_1 \quad M_1 \rightleftharpoons M_2 \\ M_3 \xrightarrow{k_{\text{out}}} 0 \qquad & M_2 + \mathrm{A}_1 \rightleftharpoons M_3 + \mathrm{A}_0. \end{aligned} \tag{4.2}$$

Considering **reversible reaction kinetics**, the evolution of the concentrations of the species in time is modelled by the following ODE system:

$$\begin{aligned}
\frac{dm_0}{dt} &= k_{\text{in}} - \frac{E_1L_0m_0a_0 - F_1K_1m_1a_1}{K_1m_1a_1 + L_0m_0a_0 + K_1L_0} \\
\frac{dm_1}{dt} &= \frac{E_1L_0m_0a_0 - F_1K_1m_1a_1}{K_1m_1a_1 + L_0m_0a_0 + K_1L_0} - \frac{E_2L_1m_1 - F_2K_2m_2}{K_2m_2 + L_1m_1 + K_2L_1} \\
\frac{dm_2}{dt} &= \frac{E_2L_1m_1 - F_2K_2m_2}{K_2m_2 + L_1m_1 + K_2L_1} - \frac{E_3L_2m_2a_1 - F_3K_3m_3a_0}{K_3m_3a_0 + L_2m_2a_1 + K_3L_2} \\
\frac{dm_3}{dt} &= \frac{E_3L_2m_2a_1 - F_3K_3m_3a_0}{K_3m_3a_0 + L_2m_2a_1 + K_3L_2} - k_{\text{out}}m_3 \\
\frac{da_0}{dt} &= -\frac{E_1L_0m_0a_0 - F_1K_1m_1a_1}{K_1m_1a_1 + L_0m_0a_0 + K_1L_0} + \frac{E_3L_2m_2a_1 - F_3K_3m_3a_0}{K_3m_3a_0 + L_2m_2a_1 + K_3L_2} \\
\frac{da_1}{dt} &= \frac{E_1L_0m_0a_0 - F_1K_1m_1a_1}{K_1m_1a_1 + L_0m_0a_0 + K_1L_0} - \frac{E_3L_2m_2a_1 - F_3K_3m_3a_0}{K_3m_3a_0 + L_2m_2a_1 + K_3L_2}.
\end{aligned}$$

The system has two conservation laws:

$$a_0 + a_1 = A_{\text{tot}}, \qquad m_1 + m_2 + a_0 = W. \tag{4.3}$$

Solving recursively the steady state equations given by $\frac{dm_0}{dt} + \cdots + \frac{dm_3}{dt} = 0$, $\frac{dm_3}{dt} = 0$, $\frac{dm_2}{dt} + \frac{dm_3}{dt} = 0$, $\frac{dm_1}{dt} + \frac{dm_2}{dt} + \frac{dm_3}{dt} = 0$ for $m_3, m_2, m_1, m_0$ and the first conservation law, we obtain

$$\begin{aligned}
m_0 &= \frac{K_1\left((F_1 + k_{\text{in}})m_1a_1 + L_0k_{\text{in}}\right)}{a_0L_0\left(E_1 - k_{\text{in}}\right)}, \\
m_1 &= \frac{K_2\left((F_2 + k_{\text{in}})m_2 + L_1k_{\text{in}}\right)}{L_1\left(E_2 - k_{\text{in}}\right)}, \\
m_2 &= \frac{k_{\text{in}}K_3\left((F_3 + k_{\text{in}})a_0 + L_2k_{\text{out}}\right)}{k_{\text{out}}a_1L_2\left(E_3 - k_{\text{in}}\right)}, \\
m_3 &= \frac{k_{\text{in}}}{k_{\text{out}}}, \qquad\qquad a_1 = A_{\text{tot}} - a_0.
\end{aligned} \tag{4.4}$$

By substituting recursively $m_2$ in $m_1$ and $m_1$ in $m_0$, we obtain expressions of all variables at steady state in terms of $a_0$. For these to be positive, it needs to hold that $0 < a_0 < A_{\text{tot}}$, and further

$$k_{\text{in}} < \min(E_1, E_2, E_3). \tag{4.5}$$

We substitute (4.4) into the remaining conservation law, and obtain a polynomial in $a_0$ whose roots in the interval $(0, T)$ are in one-to-one correspondence with the positive steady states, provided (4.5) holds. The polynomial is

$$\begin{aligned}
p(a_0) = &-L_1L_2k_{\text{out}}\left(E_3 - k_{\text{in}}\right)\left(E_2 - k_{\text{in}}\right)a_0^2 + \big(L_1L_2k_{\text{out}}\left(A_{\text{tot}} + W\right)\left(E_3 - k_{\text{in}}\right)\left(E_2 - k_{\text{in}}\right) \\
&- K_2L_1L_2k_{\text{in}}k_{\text{out}}\left(E_3 - k_{\text{in}}\right) + K_3k_{\text{in}}\left(F_3 + k_{\text{in}}\right)\left(E_2L_1 + F_2K_2\right) \\
&+ K_3k_{\text{in}}^2\left(K_2 - L_1\right)\left(F_3 + k_{\text{in}}\right)\big)a_0 + L_2K_2\,\alpha,
\end{aligned}$$

where

$$\alpha = k_{\text{in}}(k_{\text{in}} + F_2)K_2K_3 + k_{\text{in}}A_{\text{tot}}(E_3 - k_{\text{in}})K_2L_1 + k_{\text{in}}(E_2 - k_{\text{in}})K_3L_1 - \ A_{\text{tot}}\,W(E_3 - k_{\text{in}})(E_2 - k_{\text{in}})L_1. \tag{4.6}$$

As $a_0 = A_{\text{tot}}$, we have $p(A_{\text{tot}}) > 0$, and hence, by Lemma 1, the system has positive steady states if and only if $\alpha < 0$.

Note that at $k_{\text{in}} = 0$, $\alpha = -\ A_{\text{tot}}\, W(E_3 - k_{\text{in}})(E_2 - k_{\text{in}})L_1 < 0$. Hence, for $k_{\text{in}}$ small enough, $\alpha < 0$ and also (4.5) holds, and the system has a positive steady state.

The steady state value of $a_0$ is found by solving the quadratic equation $p(a_0) = 0$ and considering the smallest root. We note that $\alpha$ is a polynomial in $k_{\text{in}}$ of degree 2 with negative independent term. The sign of the leading term depends on the choice of parameters. If the minimum in (4.5) is attained at $k_{\text{in}} = E_2$ or $k_{\text{in}} = E_3$, then at this value of $k_{\text{in}}$, $\alpha$ is positive independently of the rest of the parameters. Hence, in the region of interest $\alpha$ is negative only in an interval of the form $(0, B)$, where $B$ is the smallest positive root of $\alpha$. The bound for positive steady states is now given as

$$k_{\text{in}} < \min(E_1, B).$$

If the minimum is attained at $B$, then, as $k_{\text{in}}$ approaches $B$, the smallest positive root of $p(a_0)$ approaches 0 as $\alpha$ approaches zero. Using this in (4.4), we obtain that

$$\begin{aligned} &m_0 \xrightarrow[k_{\text{in}}\to B]{} \infty, && m_1 \xrightarrow[k_{\text{in}}\to B]{} \frac{Bk_{\text{out}}\,(F_2K_3 + L_1A_{\text{tot}}\,(E_3 - B) + BK_3)}{L_1A_{\text{tot}}\,(E_3 - B)\,(E_2 - B)}, \\ &m_2 \xrightarrow[k_{\text{in}}\to B]{} \frac{BK_3}{A_{\text{tot}}\,(E_3 - B)} && m_3 \xrightarrow[k_{\text{in}}\to B]{} \tfrac{B}{k_{\text{out}}}. \end{aligned} \tag{4.7}$$

If $E_1 < B$, then $k_{\text{in}}$ needs to be smaller than $E_1$. When $k_{\text{in}}$ approaches $E_1$, $a_0$ approaches the root of $p$ when $k_{\text{in}} = E_1$, which is not zero. Then $m_0$ still approaches $\infty$ as it has $E_1 - k_{\text{in}}$ in the denominator, while $m_1, m_2$ converge to some positive value.

##### 4.2 Steady states of the linear pathway model. The general case.

We consider the dynamics of model (4.1) with $n$ taking any positive value strictly larger than 1, and positioning the co-substrate recycling in arbitrary places.

$$\begin{aligned} 0 &\xrightarrow{k_{\text{in}}} M_0 \\ M_0 &\rightleftharpoons M_1 \rightleftharpoons \cdots \rightleftharpoons M_{\ell-1} \rightleftharpoons M_\ell && M_\ell + \mathrm{A}_0 \rightleftharpoons M_{\ell+1} + \mathrm{A}_1 \\ M_{\ell+1} &\rightleftharpoons \cdots \rightleftharpoons M_{\rho-1} \rightleftharpoons M_\rho && M_\rho + \mathrm{A}_1 \rightleftharpoons M_{\rho+1} + \mathrm{A}_0 \\ M_{\rho+1} &\rightleftharpoons \cdots \rightleftharpoons M_{n-1} \rightleftharpoons M_n \\ M_n &\xrightarrow{k_{\text{out}}} 0. \end{aligned} \tag{4.8}$$

For the system to make sense biochemically, we require $0 \leq \ell < \rho - 1 \leq n - 1$.

To write the equations in a simple format, we write

$$x_i = m_i, \qquad i \neq \ell, \rho, \qquad x_\ell = m_\ell a_0, \quad x_\rho = m_\rho a_1$$

and

$$y_i = m_i, \qquad i \neq \ell + 1, \rho + 1, \qquad y_{\ell+1} = m_{\ell+1}a_1, \quad y_{\rho+1} = m_{\rho+1}a_0.$$

We start with the **reversible reaction kinetics**. Then the associated ODE system becomes:

$$\begin{aligned}
\frac{dm_0}{dt} &= k_{\text{in}} - \frac{E_1 L_0\, x_0 - F_1 K_1\, y_1}{K_1 L_0 + L_0 x_0 + K_1 y_1} \\
\vdots \quad & \quad \vdots \\
\frac{dm_i}{dt} &= \frac{E_i L_{i-1}\, x_{i-1} - F_i K_i\, y_i}{K_i L_{i-1} + K_i\, y_i + L_{i-1}\, x_{i-1}} - \frac{E_{i+1} L_i\, x_i - F_{i+1} K_{i+1}\, y_{i+1}}{K_{i+1} L_i + K_{i+1}\, y_{i+1} + L_i\, x_i} \qquad i = 1, \dots, n-1 \\
\vdots \quad & \quad \vdots \\
\frac{dm_n}{dt} &= \frac{E_n L_{n-1}\, x_{n-1} - F_n K_n\, y_n}{K_n L_{n-1} + K_n\, y_n + L_{n-1}\, x_{n-1}} - k_{\text{out}}\, m_n \\
\frac{da_0}{dt} &= -\frac{E_{\ell+1} L_\ell\, x_\ell - F_{\ell+1} K_{\ell+1}\, y_{\ell+1}}{K_{\ell+1} L_\ell + L_\ell x_\ell + K_{\ell+1} y_{\ell+1}} + \frac{E_{\rho+1} L_\rho\, x_\rho - F_{\rho+1} K_{\rho+1}\, y_{\rho+1}}{K_{\rho+1} L_\rho + L_\rho x_\rho + K_{\rho+1} y_{\rho+1}} \\
\frac{da_1}{dt} &= \frac{E_{\ell+1} L_\ell\, x_\ell - F_{\ell+1} K_{\ell+1}\, y_{\ell+1}}{K_{\ell+1} L_\ell + L_\ell x_\ell + K_{\ell+1} y_{\ell+1}} - \frac{E_{\rho+1} L_\rho\, x_\rho - F_{\rho+1} K_{\rho+1}\, y_{\rho+1}}{K_{\rho+1} L_\rho + L_\rho x_\rho + K_{\rho+1} y_{\rho+1}}.
\end{aligned}$$

This ODE system has precisely two conservation laws:

$$a_0 + a_1 = A_{\text{tot}}, \qquad m_{\ell+1} + \cdots + m_\rho + a_0 = W.$$

Note that we have the following equalities:

$$\frac{dm_0}{dt} + \cdots + \frac{dm_i}{dt} = k_{\text{in}} - \frac{E_{i+1} L_i\, x_i - F_{i+1} K_{i+1}\, y_{i+1}}{K_{i+1} L_i + K_{i+1}\, y_{i+1} + L_i\, x_i}, \qquad i = 0, \dots, n-1 \tag{4.9}$$

$$\frac{dm_0}{dt} + \cdots + \frac{dm_n}{dt} = k_{\text{in}} - k_{\text{out}}\, m_n. \tag{4.10}$$

For $i = 1, \dots, n-2$, by solving (4.9) for $x_i$, we obtain the following recursive formulas:

$$x_i = \frac{K_{i+1}(k_{\text{in}} L_i + (k_{\text{in}} + F_{i+1})\, y_{i+1})}{(E_{i+1} - k_{\text{in}}) L_i}, \qquad i = 0, \dots, n-1. \tag{4.11}$$

Finally, from (4.10) and the conservation law $a_0 + a_1 = A_{\text{tot}}$, we obtain

$$x_n = \frac{k_{\text{in}}}{k_{\text{out}}}, \qquad a_1 = A_{\text{tot}} - a_0. \tag{4.12}$$

These expressions are all positive if and only if $0 < a_0 < A_{\text{tot}}$ and $k_{\text{in}} < E_i$ for $i = 1, \dots, n$. Note that the value of $m_i$ can now be found recursively from $m_n$ using (4.11), as we show next.

Recall that $0 \le \ell < \rho - 1 \le n - 1$. Then, it holds

$$y_i = x_i \epsilon_i, \qquad \epsilon_i = 1, i \neq \ell, \ell+1, \rho, \rho+1, \qquad \epsilon_\ell = \epsilon_{\rho+1}^{-1} = \tfrac{1}{a_0}, \quad \epsilon_{\ell+1} = \epsilon_\rho^{-1} = a_1.$$

We write for shortness

$$x_i = \frac{z_i + b_i\, x_{i+1}}{c_i}, \qquad i = 0, \dots, n-1,$$

where for $i = 0, \dots, n-1$,

$$z_i = k_{\text{in}} K_{i+1} L_i \qquad b_i = \epsilon_{i+1}(k_{\text{in}} + F_{i+1}) K_{i+1} \qquad c_i = (E_{i+1} - k_{\text{in}}) L_i. \tag{4.13}$$

Then we claim that

$$x_i = \frac{\left(\sum_{j=i}^{n-1} z_j (b_i \cdots b_{j-1})(c_{j+1} \cdots c_{n-1})\right) + (b_i \cdots b_{n-1}) x_n}{c_i \cdots c_{n-1}}, \qquad i = 1, \dots, n-1. \tag{4.14}$$

We prove this by descending induction on $i$. Note that products over an empty index equal 1. For instance, $b_j \cdots b_{j-1} = 1$. For $i = n-1$, (4.14) agrees with (4.11). Assume the formula is true for some $i$, and we prove it for $i-1$. To do so, we use (4.11) and the induction hypothesis for $x_i$. For $n-1 > i-1 \geq 0$, we have

$$
\begin{aligned}
x_{i-1} &= \frac{z_{i-1} + b_{i-1}x_i}{c_{i-1}} = \frac{z_{i-1} + b_{i-1}\frac{\left(\sum_{j=i}^{n-1} z_j(b_i\cdots b_{j-1})(c_{j+1}\cdots c_{n-1})\right)+(b_i\cdots b_{n-1})x_n}{c_i\cdots c_{n-1}}}{c_{i-1}} \\
&= \frac{z_{i-1}c_i\cdots c_{n-1} + b_{i-1}\left(\left(\sum_{j=i}^{n-1} z_j(b_i\cdots b_{j-1})(c_{j+1}\cdots c_{n-1})\right) + (b_i\cdots b_{n-1})x_n\right)}{c_{i-1}c_i\cdots c_{n-1}} \\
&= \frac{z_{i-1}c_i\cdots c_{n-1} + \left(\sum_{j=i}^{n-1} z_j(b_{i-1}\cdots b_{j-1})(c_{j+1}\cdots c_{n-1})\right) + (b_{i-1}\cdots b_{n-1})x_n}{c_{i-1}\cdots c_{n-1}} \\
&= \frac{\left(\sum_{j=i-1}^{n-1} z_j(b_{i-1}\cdots b_{j-1})(c_{j+1}\cdots c_{n-1})\right) + (b_{i-1}\cdots b_{n-1})x_n}{c_{i-1}\cdots c_{n-1}}.
\end{aligned}
$$

This is (4.14) for $i-1$. Hence, the formula holds. Finally, we can use that $x_n = m_n = \frac{k_{\text{in}}}{k_{\text{out}}}$ to obtain:

$$
x_i = \frac{k_{\text{out}}\left(\sum_{j=i}^{n-1} z_j(b_i\cdots b_{j-1})(c_{j+1}\cdots c_{n-1})\right) + (b_i\cdots b_{n-1})k_{\text{in}}}{k_{\text{out}}\ c_i\cdots c_{n-1}}, \qquad i = 0,\ldots,n-1. \tag{4.15}
$$

Let

$$
\bar{b}_{u,j} := \prod_{i=u}^{j}(k_{\text{in}} + F_{i+1})K_{i+1}, \qquad \bar{c}_{u,j} := c_u\cdots c_j = \prod_{i=u}^{j}(E_{i+1} - k_{\text{in}})L_i.
$$

This gives:

$$
m_i = \frac{k_{\text{out}}\left(\sum_{j=i}^{n-1} z_j(b_i\cdots b_{j-1})\bar{c}_{j+1,n-1})\right) + (b_i\cdots b_{n-1})k_{\text{in}}}{k_{\text{out}}\ \bar{c}_{i,n-1}}, \qquad i \neq \ell,\rho, \tag{4.16}
$$

$$
m_\ell = \frac{k_{\text{out}}\left(\sum_{j=i}^{n-1} z_j(b_i\cdots b_{j-1})\bar{c}_{j+1,n-1}\right) + (b_i\cdots b_{n-1})k_{\text{in}}}{a_0 k_{\text{out}}\ \bar{c}_{\ell,n-1}}, \tag{4.17}
$$

$$
m_\rho = \frac{k_{\text{out}}\left(\sum_{j=i}^{n-1} z_j(b_i\cdots b_{j-1})\bar{c}_{j+1,n-1}\right) + (b_i\cdots b_{n-1})k_{\text{in}}}{(T-a_0)k_{\text{out}}\ \bar{c}_{\rho,n-1}}. \tag{4.18}
$$

Remember that $b_\ell, b_{\ell+1}, b_\rho, b_{\rho+1}$ depend on $a_0, a_1$, while the rest of $b$'s do not. For a product of the form $b_u\cdots b_j$ with $u \leq j$, we have the following:

If $u \leq \ell - 1$:

$$
b_u\cdots b_j = \begin{cases} \prod_{i=u}^{j}(k_{\text{in}} + F_{i+1})K_{i+1} & j < \ell - 1, \text{ or } \rho \leq j \\ \frac{1}{a_0}\prod_{i=u}^{j}(k_{\text{in}} + F_{i+1})K_{i+1} & j = \ell - 1, \text{ or } j = \rho - 1 \\ \frac{a_1}{a_0}\prod_{i=u}^{j}(k_{\text{in}} + F_{i+1})K_{i+1} & \ell - 1 < j < \rho - 1. \end{cases}
$$

If $u = \ell$:

$$
b_u\cdots b_j = \begin{cases} a_1\prod_{i=u}^{j}(k_{\text{in}} + F_{i+1})K_{i+1} & j < \rho - 1 \\ \prod_{i=u}^{j}(k_{\text{in}} + F_{i+1})K_{i+1} & j = \rho - 1 \\ a_0\prod_{i=u}^{j}(k_{\text{in}} + F_{i+1})K_{i+1} & \rho \leq j. \end{cases}
$$

If $\ell < u \leq \rho - 1$:

$$
b_u\cdots b_j = \begin{cases} \prod_{i=u}^{j}(k_{\text{in}} + F_{i+1})K_{i+1} & j < \rho - 1 \\ \frac{1}{a_1}\prod_{i=u}^{j}(k_{\text{in}} + F_{i+1})K_{i+1} & j = \rho - 1 \\ \frac{a_0}{a_1}\prod_{i=u}^{j}(k_{\text{in}} + F_{i+1})K_{i+1} & \rho \leq j. \end{cases}
$$

If $u = \rho$:

$$
b_u\cdots b_j = a_0\prod_{i=u}^{j}(k_{\text{in}} + F_{i+1})K_{i+1}.
$$

If $u > \rho$:

$$b_u \cdots b_j = \prod_{i=u}^{j} (k_{\text{in}} + F_{i+1}) K_{i+1}.$$

Summarising, $b_u \cdots b_j$ equals

$$\begin{cases} \prod_{i=u}^{j}(k_{\text{in}} + F_{i+1})K_{i+1} & u \leq \ell - 1, j < \ell - 1, \text{ or } u \leq \ell - 1, \rho \leq j, \text{ or } u = \ell, j = \rho - 1, \\ & \text{or } \ell < u \leq \rho - 1, j < \rho - 1, \text{ or } u > \rho \\ \frac{1}{a_0} \prod_{i=u}^{j}(k_{\text{in}} + F_{i+1})K_{i+1} & j = \ell - 1, \text{ or } u \leq \ell - 1, j = \rho - 1 \\ \frac{a_1}{a_0} \prod_{i=u}^{j}(k_{\text{in}} + F_{i+1})K_{i+1} & u \leq \ell - 1, \ell - 1 < j < \rho - 1 \\ a_1 \prod_{i=u}^{j}(k_{\text{in}} + F_{i+1})K_{i+1} & u = \ell, j < \rho - 1 \\ a_0 \prod_{i=u}^{j}(k_{\text{in}} + F_{i+1})K_{i+1} & u = \ell, \rho \leq j, \text{ or } u = \rho \\ \frac{1}{a_1} \prod_{i=u}^{j}(k_{\text{in}} + F_{i+1})K_{i+1} & \ell < u \leq \rho - 1, j = \rho - 1 \\ \frac{a_0}{a_1} \prod_{i=u}^{j}(k_{\text{in}} + F_{i+1})K_{i+1} & \ell < u \leq \rho - 1, \rho \leq j. \end{cases}$$

Observe that for all $i$, the summand of $\sum_{j=i}^{n-1} z_j(b_i \cdots b_{j-1})(c_{j+1} \cdots c_{n-1})$ corresponding to $i = j$ is

$$z_i(c_{i+1} \cdots c_{n-1}),$$

which does not depend either on $a_0$ or $a_1$.

In particular, we deduce that $m_i$ for $i \leq \ell - 1$, the term $(b_i \cdots b_{n-1})$ does not depend on $a_0, a_1$, and in the sum $\sum_{j=i}^{n-1} z_j(b_i \cdots b_{j-1})(c_{j+1} \cdots c_{n-1})$ there are summands involving $\frac{1}{a_0}$, e.g. when $j = \ell$. Hence $m_i$, for $i \leq \ell - 1$, goes to infinity as $a_0$ goes to zero. Note that $a_1$ goes to $T$ when $a_0$ goes to zero. When $i = \ell$, the denominator of $m_i$ is multiplied by $a_0$. As the numerator has at least one term that is not multiple of $a_0$, $m_\ell$ goes to infinity as well. We conclude that

$$m_i \xrightarrow[a_0 \to 0]{} \infty, \qquad 0 \leq i \leq \ell.$$

When $i \geq \ell$, then no summand in the numerator of $m_i$ in (4.16)-(4.18) that involves $\frac{1}{a_0}$, and neither the denominator has $a_0$ as factor. As the numerator has at least one term that is not multiple of $a_0$, $m_i$ goes to a finite value as $a_0$ goes to zero.

$$m_i \xrightarrow[a_0 \to 0]{} \text{number}, \qquad i \geq \ell.$$

The number can be found using (4.16)-(4.18) and is a function of the parameters of the system, not involving $W$.

We consider now the remaining equation, namely the conservation law $m_{\ell+1} + \cdots + m_\rho + a_0 = W$, to determine the value of $a_0$. We have

$$\sum_{i=\ell+1}^{\rho} m_i + a_0 - W = \sum_{i=\ell+1}^{\rho-1} \frac{k_{\text{out}} \left( \sum_{j=i}^{n-1} z_j(b_i \cdots b_{j-1}) \bar{c}_{j+1,n-1} \right) + (b_i \cdots b_{n-1}) k_{\text{in}}}{k_{\text{out}}\ \bar{c}_{i,n-1}}$$

$$+ \frac{k_{\text{out}} \left( \sum_{j=\rho}^{n-1} z_j(b_\rho \cdots b_{j-1}) \bar{c}_{j+1,n-1} \right) + (b_\rho \cdots b_{n-1}) k_{\text{in}}}{a_1 k_{\text{out}}\ \bar{c}_{\rho,n-1}} + a_0 - W = (*).$$

We have $(b_i \cdots b_{n-1}) = \frac{a_0}{a_1} \bar{b}_{i,n-1}$ when $\ell < i < \rho$ and $(b_\rho \cdots b_{n-1}) = a_0 \bar{b}_{\rho,n-1}$. Also

$$\sum_{j=\rho}^{n-1} z_j(b_\rho \cdots b_{j-1}) \bar{c}_{j+1,n-1} = z_\rho \bar{c}_{\rho+1,n-1} + \sum_{j=\rho+1}^{n-1} a_0 z_j \bar{b}_{\rho,j-1} \bar{c}_{j+1,n-1}.$$

Finally, for $\ell+1 \le i \le \rho-1$,

$$\begin{aligned}\sum_{j=i}^{n-1} z_j(b_i\cdots b_{j-1})\bar{c}_{j+1,n-1} &= \sum_{j=i}^{\rho-1} z_j(b_i\cdots b_{j-1})\bar{c}_{j+1,n-1} + z_\rho(b_i\cdots b_{\rho-1})\bar{c}_{\rho+1,n-1}\\ &\quad + \sum_{j=\rho+1}^{n-1} z_j(b_i\cdots b_{j-1})\bar{c}_{j+1,n-1}\\ &= \sum_{j=i}^{\rho-1} z_j\bar{b}_{i,j-1}\bar{c}_{j+1,n-1} + z_\rho\tfrac{1}{a_1}\bar{b}_{i,\rho-1}\bar{c}_{\rho+1,n-1} + \tfrac{a_0}{a_1}\sum_{j=\rho+1}^{n-1} z_j\bar{b}_{i,j-1}\bar{c}_{j+1,n-1}.\end{aligned}$$

This gives

$$\begin{aligned}(*) &= \sum_{i=\ell+1}^{\rho-1}\frac{k_{\text{out}}\left(\sum_{j=i}^{\rho-1} z_j\bar{b}_{i,j-1}\bar{c}_{j+1,n-1} + z_\rho\frac{1}{a_1}\bar{b}_{i,\rho-1}\bar{c}_{\rho+1,n-1} + \frac{a_0}{a_1}\sum_{j=\rho+1}^{n-1} z_j\bar{b}_{i,j-1}\bar{c}_{j+1,n-1}\right) + \frac{a_0}{a_1}\bar{b}_{i,n-1}k_{\text{in}}}{k_{\text{out}}\ \bar{c}_{i,n-1}}\\ &\quad + \frac{k_{\text{out}}\left(z_\rho\bar{c}_{\rho+1,n-1} + \sum_{j=\rho+1}^{n-1} a_0 z_j\bar{b}_{\rho,j-1}\bar{c}_{j+1,n-1}\right) + a_0\bar{b}_{\rho,n-1}k_{\text{in}}}{a_1 k_{\text{out}}\ \bar{c}_{\rho,n-1}} + a_0 - W\\ &= \sum_{i=\ell+1}^{\rho-1}\frac{k_{\text{out}}\left(a_1\sum_{j=i}^{\rho-1} z_j\bar{b}_{i,j-1}\bar{c}_{j+1,n-1} + z_\rho\bar{b}_{i,\rho-1}\bar{c}_{\rho+1,n-1} + a_0\sum_{j=\rho+1}^{n-1} z_j\bar{b}_{i,j-1}\bar{c}_{j+1,n-1}\right) + a_0\bar{b}_{i,n-1}k_{\text{in}}}{a_1 k_{\text{out}}\ \bar{c}_{i,n-1}}\\ &\quad + \frac{k_{\text{out}}\left(z_\rho\bar{c}_{\rho+1,n-1} + \sum_{j=\rho+1}^{n-1} a_0 z_j\bar{b}_{\rho,j-1}\bar{c}_{j+1,n-1}\right) + a_0\bar{b}_{\rho,n-1}k_{\text{in}}}{a_1 k_{\text{out}}\ \bar{c}_{\rho,n-1}} + a_0 - W.\end{aligned}$$

Hence $(*)$ vanishes if and only if

$$\begin{aligned}0 = \sum_{i=\ell+1}^{\rho-1} k_{\text{out}}&\left(a_1\sum_{j=i}^{\rho-1} z_j\bar{b}_{i,j-1}\bar{c}_{\ell+1,i-1}\bar{c}_{j+1,n-1} + z_\rho\bar{b}_{i,\rho-1}\bar{c}_{\ell+1,i-1}\bar{c}_{\rho+1,n-1} + a_0\sum_{j=\rho+1}^{n-1} z_j\bar{b}_{i,j-1}\bar{c}_{\ell+1,i-1}\bar{c}_{j+1,n-1}\right)\\ &+ a_0\bar{b}_{i,n-1}\bar{c}_{\ell+1,i-1}k_{\text{in}} + k_{\text{out}}\left(z_\rho\bar{c}_{\ell+1,\rho-1}\bar{c}_{\rho+1,n-1} + \sum_{j=\rho+1}^{n-1} a_0 z_j\bar{b}_{\rho,j-1}\bar{c}_{\ell+1,\rho-1}\bar{c}_{j+1,n-1}\right)\\ &+ a_0\bar{b}_{\rho,n-1}\bar{c}_{\ell+1,\rho-1}k_{\text{in}} + (a_0 - W)k_{\text{out}}\,\bar{c}_{\ell+1,n-1}\,a_1.\end{aligned} \tag{4.19}$$

As $a_1 = A_{\text{tot}} - a_0$, this is a degree 2 polynomial in $a_0$. The leading term comes from the term $(a_0 - W)k_{\text{out}}\,\bar{c}_{\ell+1,n-1}\,a_1$, and is

$$-k_{\text{out}}\,\bar{c}_{\ell+1,n-1}a_0^2,$$

which is negative. The independent term is

$$\begin{aligned}\sum_{i=\ell+1}^{\rho-1} k_{\text{out}}&\left(A_{\text{tot}}\sum_{j=i}^{\rho-1} z_j\bar{b}_{i,j-1}\bar{c}_{\ell+1,i-1}\bar{c}_{j+1,n-1} + z_\rho\bar{b}_{i,\rho-1}\bar{c}_{\ell+1,i-1}\bar{c}_{\rho+1,n-1}\right)\\ &+ k_{\text{out}}z_\rho\bar{c}_{\ell+1,\rho-1}\bar{c}_{\rho+1,n-1} - A_{\text{tot}}W k_{\text{out}}\,\bar{c}_{\ell+1,n-1}.\end{aligned}$$

We divide by $k_{\text{out}}$ and define

$$\alpha = \sum_{i=\ell+1}^{\rho-1}\bar{c}_{\ell+1,i-1}\left(A_{\text{tot}}\sum_{j=i}^{\rho-1} z_j\bar{b}_{i,j-1}\bar{c}_{j+1,n-1} + z_\rho\bar{b}_{i,\rho-1}\bar{c}_{\rho+1,n-1}\right) + z_\rho\bar{c}_{\ell+1,\rho-1}\bar{c}_{\rho+1,n-1} - A_{\text{tot}}W\,\bar{c}_{\ell+1,n-1},$$

where recall from (4.13) that

$$z_i = k_{\text{in}}K_{i+1}L_i \qquad b_i = \epsilon_{i+1}(k_{\text{in}} + F_{i+1})K_{i+1} \qquad c_i = (E_{i+1} - k_{\text{in}})L_i.$$

When $a_0 = A_{\text{tot}}$, all terms multiplying $a_1$ vanish, and then the polynomial is a sum of positive terms, hence positive. By Lemma 1, the system has a positive steady state if and only if

$$\alpha < 0.$$

Note that when $k_{\text{in}} = 0$ this inequality holds, as all terms with $z_j$ vanish. When $k_{\text{in}}$ approaches one of $E_i$ with $\ell + 1 < i \leq n$, the negative term of $\alpha$ approaches zero, and the inequality does not hold.

For example, for $n = 3$, $\ell = 0$ and $\rho = 2$, $\alpha$ was found in (4.6). For $n = 4$, $\ell = 0$ and $\rho = 2$, we have

$$\begin{aligned}\alpha = &-A_{\text{tot}}W(E_2 - k_{\text{in}})L_1(E_3 - k_{\text{in}})L_2(E_4 - k_{\text{in}}) + k_{\text{in}}K_4(E_2 - k_{\text{in}})L_1(E_3 - k_{\text{in}})L_2\\ &+ A_{\text{tot}}k_{\text{in}}k_{\text{out}}L_1(E_4 - k_{\text{in}})(E_3 - k_{\text{in}})L_2 + A_{\text{tot}}k_{\text{in}}K_3L_2(E_4 - k_{\text{in}})(F_2 + k_{\text{in}})k_{\text{out}}\\ &+ k_{\text{in}}K_4(F_3 + k_{\text{in}})K_3(F_2 + k_{\text{in}})k_{\text{out}} + k_{\text{in}}K_3L_2(E_2 - k_{\text{in}})L_1(E_4 - k_{\text{in}})A_{\text{tot}}\\ &+ k_{\text{in}}K_4(E_2 - k_{\text{in}})L_1(F_3 + k_{\text{in}})K_3.\end{aligned}$$

Let $B$ be the smallest positive root of $\alpha = 0$ as a polynomial in $k_{\text{in}}$, if it exists, or take $B = \infty$ if not. Similarly, let $B'$ be the second such root, if it exists, or $B' = \infty$ if not. If the smallest of $E_j$ is attained for some $E_i$ with $\ell + 1 < i \leq n$, then $\alpha$ is positive at $k_{\text{in}} = E_i$. In that case, as $\alpha$ is negative at $k_{\text{in}} = 0$, there is at least one value of $k_{\text{in}} < E_i$ for all $i$ such that $\alpha = 0$ and hence $B$ is finite. This shows that $\min(E_1, \ldots, E_n, B) = \min(E_1, \ldots, E_{\ell+1}, B)$.

Putting it all together, we have shown that for all

$$k_{\text{in}} < \min(E_1, \ldots, E_{\ell+1}, B) \tag{4.20}$$

the system has positive steady states, and if

$$k_{\text{in}} \geq \min(E_1, \ldots, E_{\ell+1}, B), \qquad \text{or} \qquad k_{\text{in}} < \min(E_1, \ldots, E_{\ell+1}), \quad B < k_{\text{in}} < B',$$

then the system has no positive steady state.

Taking condition (4.20), if the minimum is attained at $B$, then when $k_{\text{in}}$ approaches $B$, the first positive root of the polynomial in (4.19) approaches zero (as $\alpha$ goes to zero). As this determines the steady state value of $a_0$, we see that $a_0$ approaches zero, and the $m_i$ converge to the values given above. Specifically,

$$m_j \xrightarrow[k_{\text{in}} \to B]{} \infty, \qquad \text{for } 0 \leq j \leq \ell, \qquad m_j \xrightarrow[k_{\text{in}} \to B]{} \text{number}, \qquad \text{for } j \geq \ell.$$

By the comment above, (4.20) cannot be attained at $E_i$ with $\ell + 1 < i \leq n$. If (4.20) is attained at $E_i$ with $i \leq \ell + 1$, then as $k_{\text{in}}$ approaches this minimum, $a_0$ converges to a number. In this case, the concentrations that tend to infinity are those with $E_i - k_{\text{in}}$ in the denominator:

$$m_j \xrightarrow[k_{\text{in}} \to E_i]{} \infty, \qquad \text{for } 0 \leq j < i, \qquad m_j \xrightarrow[k_{\text{in}} \to E_i]{} \text{number}, \qquad \text{for } j \geq i.$$

### 5 Multiple co-substrate cycling along a single pathway – mimicking the case seen in glycolysis, combined with fermentation

In this section, we consider a scenario of intra-pathway cycling with two different co-substrates. This is a simplified version of the case seen in the combined pathways of glycolysis and fermentation. The reaction scheme we consider comprises:

$$\begin{array}{ccc} 0 \xrightarrow{k_{\text{in}}} M_0 & M_0 + \text{ATP} \rightleftharpoons M_1 + \text{ADP} & M_1 + \text{NAD} \rightleftharpoons M_2 + \text{NADH} \\ M_4 \xrightarrow{k_{\text{out}}} 0 & M_2 + \text{ADP} \rightleftharpoons M_3 + \text{ATP} & M_3 + \text{NADH} \rightleftharpoons M_4 + \text{NAD}. \end{array} \tag{5.1}$$

We write $A_0$ =ATP, $A_1$ =ADP, $A_2$ =NAD, $A_3$ =NADH, for simplicity. The ODE system governing the dynamics of the network is:

$$
\begin{aligned}
\frac{dm_0}{dt} &= k_{\text{in}} - \frac{E_1 L_0 a_0 m_0 - F_1 K_1 a_1 m_1}{K_1 a_1 m_1 + L_0 a_0 m_0 + K_1 L_0} \\
\frac{dm_1}{dt} &= \frac{E_1 L_0 a_0 m_0 - F_1 K_1 a_1 m_1}{K_1 a_1 m_1 + L_0 a_0 m_0 + K_1 L_0} - \frac{E_2 L_1 a_2 m_1 - F_2 K_2 a_3 m_2}{K_2 a_3 m_2 + L_1 a_2 m_1 + K_2 L_1} \\
\frac{dm_2}{dt} &= \frac{E_2 L_1 a_2 m_1 - F_2 K_2 a_3 m_2}{K_2 a_3 m_2 + L_1 a_2 m_1 + K_2 L_1} - \frac{E_3 L_2 a_1 m_2 - F_3 K_3 a_0 m_3}{K_3 a_0 m_3 + L_2 a_1 m_2 + K_3 L_2} \\
\frac{dm_3}{dt} &= \frac{E_3 L_2 a_1 m_2 - F_3 K_3 a_0 m_3}{K_3 a_0 m_3 + L_2 a_1 m_2 + K_3 L_2} - \frac{E_4 L_3 a_3 m_3 - F_4 K_4 a_2 m_4}{K_4 a_2 m_4 + L_3 a_3 m_3 + K_4 L_3} \\
\frac{dm_4}{dt} &= \frac{E_4 L_3 a_3 m_3 - F_4 K_4 a_2 m_4}{K_4 a_2 m_4 + L_3 a_3 m_3 + K_4 L_3} - k_{\text{out}} m_4 \\
\frac{da_0}{dt} &= -\frac{E_1 L_0 a_0 m_0 - F_1 K_1 a_1 m_1}{K_1 a_1 m_1 + L_0 a_0 m_0 + K_1 L_0} - \frac{E_3 L_2 a_1 m_2 - F_3 K_3 a_0 m_3}{K_3 a_0 m_3 + L_2 a_1 m_2 + K_3 L_2} \\
\frac{da_1}{dt} &= \frac{E_1 L_0 a_0 m_0 - F_1 K_1 a_1 m_1}{K_1 a_1 m_1 + L_0 a_0 m_0 + K_1 L_0} - \frac{E_3 L_2 a_1 m_2 - F_3 K_3 a_0 m_3}{K_3 a_0 m_3 + L_2 a_1 m_2 + K_3 L_2} \\
\frac{da_2}{dt} &= -\frac{E_2 L_1 a_2 m_1 - F_2 K_2 a_3 m_2}{K_2 a_3 m_2 + L_1 a_2 m_1 + K_2 L_1} + \frac{E_4 L_3 a_3 m_3 - F_4 K_4 a_2 m_4}{K_4 a_2 m_4 + L_3 a_3 m_3 + K_4 L_3} \\
\frac{da_3}{dt} &= \frac{E_2 L_1 a_2 m_1 - F_2 K_2 a_3 m_2}{K_2 a_3 m_2 + L_1 a_2 m_1 + K_2 L_1} - \frac{E_4 L_3 a_3 m_3 - F_4 K_4 a_2 m_4}{K_4 a_2 m_4 + L_3 a_3 m_3 + K_4 L_3}.
\end{aligned}
$$

There are four conservation laws:

$$a_0 + a_1 = A_{\text{tot}}, \qquad a_2 + a_3 = B_{\text{tot}}, \qquad m_1 + m_2 + a_0 = W, \qquad m_2 + m_3 + a_2 = M.$$

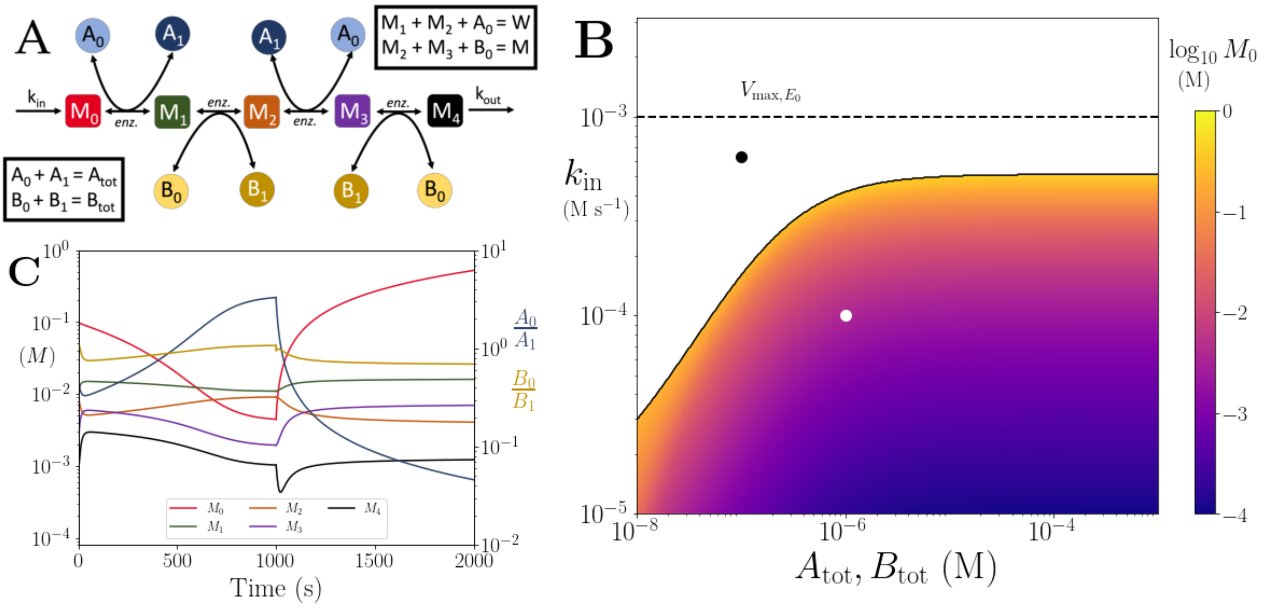

Figure S3: **(A)** Cartoon representation of a chain of reversible reactions with metabolite cycling of two different metabolites, as shown in Eq. 5.1. Each cycled metabolite has two forms, and there is no pathway independent cycling. **(B)** Heatmap of of the steady state concentration of $M_0$ as a function of the metabolite pool sizes ($A_{\text{tot}} = B_{\text{tot}}$) and the inflow flux ($k_{\text{in}}$). White area shows the region where there is no steady state. **(C)** Concentrations of $M_{0-4}$, $A_0/A_1$ and $B_0/B_1$ ratios as a function of time. At $t = 1000$s, parameters are switched from the white dot in panel (B) (where a steady state exists) to the black dot (where we see build up of $M_0$), and continued decline of $A_0$. The $B_0/B_1$ ratio remains constant however as these are still cycled by reactions after the build up. In (B) and (C) the other parameters are $k_{\text{cat}} = 100s^{-1}$, $E_{\text{tot}} = 0.01mM$, $K_m = 50\mu M$ and $k_{\text{out}} = 0.1s^{-1}$.

We consider the equations $\frac{dm_0}{dt} + \cdots + \frac{dm_4}{dt} = 0$, $\frac{dm_4}{dt} = 0$, $\frac{dm_3}{dt} + \frac{dm_4}{dt} = 0$, $\frac{dm_2}{dt} + \frac{dm_3}{dt} + \frac{dm_4}{dt} = 0$, $\frac{dm_1}{dt} + \frac{dm_2}{dt} + \frac{dm_3}{dt} + \frac{dm_4}{dt} = 0$, and solve them iteratively for $m_4, m_3, m_2, m_1, m_0$ and obtain:

$$m_4 = \frac{k_{\text{in}}}{k_{\text{out}}}, \qquad m_3 = \frac{k_{\text{in}} K_4 \left((F_4 + k_{\text{in}})a_2 + L_3 k_{\text{out}}\right)}{k_{\text{out}} a_3 L_3 \left(E_4 - k_{\text{in}}\right)},$$
$$m_2 = \frac{K_3 \left((F_3 + k_{\text{in}})a_0 m_3 + L_2 k_{\text{in}}\right)}{a_1 L_2 \left(E_3 - k_{\text{in}}\right)}, \qquad m_1 = \frac{K_2 \left((F_2 + k_{\text{in}})a_3 m_2 + L_1 k_{\text{in}}\right)}{a_2 L_1 \left(E_2 - k_{\text{in}}\right)},$$
$$m_0 = \frac{K_1 \left((F_1 + k_{\text{in}})a_1 m_1 + L_0 k_{\text{in}}\right)}{a_0 L_0 \left(E_1 - k_{\text{in}}\right)},$$

and $a_1 = A_{\text{tot}} - a_0$, $a_3 = B_{\text{tot}} - a_2$. As usual, a necessary condition for positive steady states is

$$k_{\text{in}} < \min(E_1, E_2, E_3, E_4). \tag{5.2}$$

For this system, we are left with two conservation laws that we have not used, and two steady state values that are still free. By plugging the expressions above into $m_1 + m_2 + a_0 = W$, and $m_2 + m_3 + a_2 = M$, we obtain a system of two polynomial equations in two variables. For some combinations of parameter values, there will be positive steady states, and for others, not, as the following examples shows.

By choosing

$$\begin{aligned} &M = 1, \quad W = 1, \quad F_1 = 1 \quad F_2 = 1, \quad F_3 = 1, \quad F_4 = 2, \quad K_1 = 1 \quad K_2 = 0.1, \quad K_3 = 0.1,\\ &K_4 = 2, \quad L_0 = 1 \quad L_1 = 1 \quad L_2 = 2, \quad L_3 = 10, \quad A_{\text{tot}} = 3, \quad B_{\text{tot}} = 2, \quad E_1 = 3, \quad E_2 = 3,\\ &E_3 = 12, \quad E_4 = 12, \quad k_{\text{in}} = 2, \quad k_{\text{out}} = 10, \end{aligned}$$

the polynomial system becomes

$$0 = 20000a_2^2a_0 - 60000a_2^2 + 180560a_2 - 60315.2a_2a_0 - 95200 + 32120a_0$$
$$0 = 20000a_2^2a_0^2 - 40000a_2a_0^2 - 79996.64a_2^2a_0 + 164086.88a_2a_0 - 7928a_0 + 59720a_2^2 - 131680a_2 + 24480$$

and the solutions are:

$$a_0 \sim 0.69, \qquad a_2 \sim 0.679,$$

which are in the desired interval.

If instead we replace $L_3$ by 1, we obtain a system whose solutions are

$$a_0 \sim 2.978, \qquad a_2 \sim -0.323,$$

and hence there are no positive steady states.

This shows that there is a condition for positive steady states to exist, and although (10.3) is necessary, it is not sufficient.

### 6 Different stoichiometries for co-substrate cycling along a single pathway – mimicking the case seen in upper glycolysis

In this section, we consider a scenario of intra-pathway cycling with varying stoichiometry in the pathway. In particular, we consider a simplified version of the case seen in upper glycolysis. The reaction scheme we consider comprises:

$$\begin{aligned} &0 \xrightarrow{k_{\text{in}}} M_0 \qquad M_0 + \text{ATP} \rightleftharpoons M_1 + \text{ADP} \qquad M_1 + \text{ATP} \rightleftharpoons 2M_2 + \text{ADP}\\ &\qquad\qquad M_2 + \text{ADP} \rightleftharpoons M_3 + \text{ATP} \qquad M_3 \xrightarrow{k_{\text{out}}} 0. \end{aligned} \tag{6.1}$$

We let $A_0 = \text{ATP}$ and $A_1 = \text{ADP}$ as usual.

With Michaelis-Menten kinetics, the ODE system is

$$\frac{dm_0}{dt} = k_{\text{in}} - \frac{E_1L_0m_0a_0 - F_1K_1m_1a_1}{K_1m_1a_1 + L_0m_0a_0 + K_1L_0}$$
$$\frac{dm_1}{dt} = \frac{E_1L_0m_0a_0 - F_1K_1m_1a_1}{K_1m_1a_1 + L_0m_0a_0 + K_1L_0} - \frac{E_2L_1m_1a_0 - F_2K_2m_2^2a_1}{K_2m_2^2a_1 + L_1m_1a_0 + K_2L_1}$$
$$\frac{dm_2}{dt} = \frac{2(E_2L_1m_1a_0 - F_2K_2m_2^2a_1)}{K_2m_2^2a_1 + L_1m_1a_0 + K_2L_1} - \frac{E_3L_2m_2a_1 - F_3K_3m_3a_0}{K_3m_3a_0 + L_2m_2a_1 + K_3L_2}$$
$$\frac{dm_3}{dt} = \frac{E_3L_2m_2a_1 - F_3K_3m_3a_0}{K_3m_3a_0 + L_2m_2a_1 + K_3L_2} - k_{\text{out}}m_3$$
$$\frac{da_0}{dt} = -\frac{E_1L_0m_0a_0 - F_1K_1m_1a_1}{K_1m_1a_1 + L_0m_0a_0 + K_1L_0} - \frac{E_2L_1m_1a_0 - F_2K_2m_2^2a_1}{K_2m_2^2a_1 + L_1m_1a_0 + K_2L_1} + \frac{E_3L_2m_2a_1 - F_3K_3m_3a_0}{K_3m_3a_0 + L_2m_2a_1 + K_3L_2}$$
$$\frac{da_1}{dt} = \frac{E_1L_0m_0a_0 - F_1K_1m_1a_1}{K_1m_1a_1 + L_0m_0a_0 + K_1L_0} + \frac{E_2L_1m_1a_0 - F_2K_2m_2^2a_1}{K_2m_2^2a_1 + L_1m_1a_0 + K_2L_1} - \frac{E_3L_2m_2a_1 - F_3K_3m_3a_0}{K_3m_3a_0 + L_2m_2a_1 + K_3L_2}.$$

The system has two conservation laws:

$$a_0 + a_1 = A_{\text{tot}}, \qquad m_1 + m_2 + a_0 = W.$$

By considering the equation $0 = 2\frac{dm_0}{dt} + 2\frac{dm_1}{dt} + \frac{dm_2}{dt} + \frac{dm_3}{dt} = k_{\text{out}}m_3 - 2k_{\text{in}}$, we obtain

$$m_3 = \frac{2k_{\text{in}}}{k_{\text{out}}}.$$

Upon substitution of this value of $m_3$ into $\frac{dm_3}{dt} = 0$, $\frac{dm_2}{dt} + \frac{dm_3}{dt} = 0$ and $2\frac{dm_1}{dt} + \frac{dm_2}{dt} + \frac{dm_3}{dt} = 0$, and solving recursively for $M_2, m_1, m_0$, we obtain the following steady state relations:

$$m_2 = \frac{2k_{\text{in}}K_3\left((F_3 + 2k_{\text{in}})a_0 + L_2k_{\text{out}}\right)}{k_{\text{out}}a_1L_2\left(E_3 - 2k_{\text{in}}\right)},$$
$$m_1 = -\frac{K_2\left((F_2 + k_{\text{in}})a_1m_2^2 + L_1k_{\text{in}}\right)}{L_1a_0\left(E_2 - k_{\text{in}}\right)},$$
$$m_0 = -\frac{K_1\left((F_1 + k_{\text{in}})a_1m_1 + L_0k_{\text{in}}\right)}{a_0L_0\left(E_1 - k_{\text{in}}\right)},$$

and recall $a_1 = A_{\text{tot}} - a_0$. We see that if $0 < a_0 < A_{\text{tot}}$, these expressions are all positive if and only if

$$k_{\text{in}} < \min(E_1, E_2, E_3/2). \tag{6.2}$$

As usual, we consider the remaining conservation law, $m_1 + m_2 + a_0 = W$, together with these expressions, to find a polynomial $p(a_0)$ whose roots in $(0, A_{\text{tot}})$ are in one-to-one correspondence with positive steady states, provided (10.4) holds. The polynomial has degree 3, positive leading term, negative independent term and term of degree 2, and is positive at $a_0 = A_{\text{tot}}$. With this information, we cannot immediately conclude that there is a positive root in $(0, A_{\text{tot}})$. But for positive steady states to exist, we need the term of degree 1 to be positive (this follows from Descartes' rule of signs, as usual), and this sets an extra condition on the parameters. When this happens, there will be two positive steady states.

Specifically, the term of degree 1 is

$$\begin{aligned}\alpha = L_2k_{\text{out}}\Big(&L_1L_2\left(E_3 - 2k_{\text{in}}\right)^2k_{\text{out}}\left(A_{\text{tot}}W(E_2 - k_{\text{in}}) + K_2k_{\text{in}}\right)\\ &- 2K_3L_1L_2\left(E_2 - k_{\text{in}}\right)\left(E_3 - 2k_{\text{in}}\right)k_{\text{in}}k_{\text{out}}\\ &- 8K_2K_3^2k_{\text{in}}^2\left(F_3 + 2k_{\text{in}}\right)\left(F_2 + k_{\text{in}}\right)\Big),\end{aligned}$$

which depends on $A_{\text{tot}}$ and $W$. To summarize for positive steady states to exist, we need

$$k_{\text{in}} < E_1, \qquad k_{\text{in}} < E_2, \qquad k_{\text{in}} < \tfrac{E_3}{2}, \qquad \alpha > 0.$$

If $k_{\text{in}}$ is small enough, then the conditions hold as $\alpha > 0$ at $k_{\text{in}} = 0$.

However, these conditions are not sufficient. To see this, by inspecting the term $\alpha$, one can see that if $K_2, K_3$ are small enough, then there will be two positive steady states, while if they are larger, there will be none. We have verified that both scenarios occur. For example, fix the parameter values to be

$$\begin{array}{ccccccc} A_{\text{tot}} = 1, & W = 2, & F_1 = 1, & F_2 = 2, & F_3 = 3, & L_0 = 1 & L_1 = 1, \\ L_2 = 2, & K_1 = 1 & k_{\text{in}} = 1, & k_{\text{out}} = 1, & E_1 = 2, & E_2 = 2, & E_3 = 3, \end{array}$$

and note that (10.4) holds. With $K_2 = K_3 = 0.1$, the polynomial of interest becomes $p(a_0) = 4a_0^3 - 14.300a_0^2 + 7.360a_0 - 0.448$, and so $\alpha > 0$. The polynomial has two positive roots under $A_{\text{tot}} = 1$, namely 0.07 and 0.537. For $K_2 = 0.2, K_3 = 0.3$, the polynomial is $p(a_0) = 4a_0^3 - 23.400a_0^2 + 2.080a_0 - 1.664$, and although $\alpha > 0$, it has no root in the interval $(0, 1)$.

An analogous reasoning holds for the irreversible system.

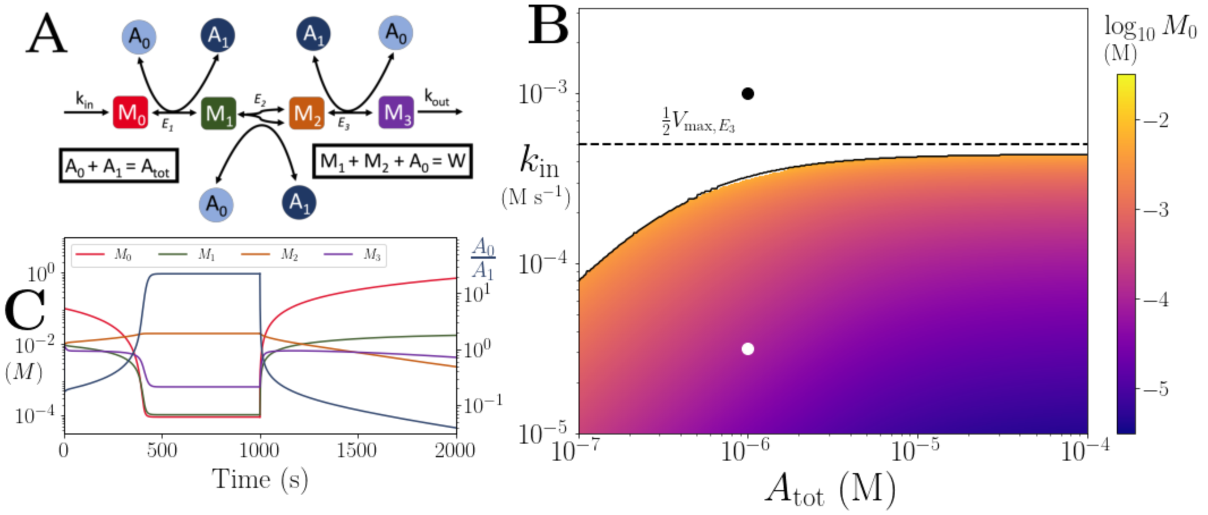

Figure S4: **(A)** Reaction system modelled in Sec. 6. Branching arrowhead from $M_1$ to $M_2$ indicates that two $M_2$ molecules are produced/used in the forward/backward reactions. **(B)** Heatmap of the steady state $M_0$ concentration. In the white area there is no steady state due to continual build up of $M_0$. Dashed line shows the smallest limit imposed by Eq. 10.4. **(C)** Time series of $M_{0\to 4}$ and $A_0/A_1$. At $t = 1000$s, parameters are switched from the white dot in panel (B) (where a steady state exists) to the black dot (where we see build up of $M_0$). Note that in the build up regime, $M_2$ reduces as well as $A_0/A_1$, as $M_2$ production is dependent on the presence of $A_0$. In (B) and (C) the other parameters are $k_{\text{cat}} = 100s^{-1}$, $E_{\text{tot}} = 0.01mM$, $K_m = 50\mu M$ and $k_{\text{out}} = 0.1s^{-1}$.

#### 7 Co-substrate cycling along with metabolite cycling – mimicking the case seen in nitrogen assimilation

In this section, we consider a scenario of intertwined, co-substrate cycling within a pathway. In particular, we consider a simplified version of the case seen in nitrogen assimilation where NAD(P)H cycling co-occurs together with a cycling from alpha-ketoglutarate to glutamate (mediated by glutamate dehydrogenase) and back from glutamate to alpha-ketoglutarate and glutamine (mediated by glutamate synthase). The simplified reaction scheme we consider comprises:

$$\begin{array}{ll} 0 \xrightarrow{k_{\text{in}}} \text{NH3} & M_0 + \text{NADH} + \text{NH3} \rightleftharpoons M_1 + \text{NAD} \\ M_2 \xrightarrow{k_{\text{out}}} 0 & M_1 + \text{NAD} \rightleftharpoons M_0 + M_2 + \text{NADH}. \end{array} \tag{7.1}$$

Denoting $A_0$ =NADH, $A_1$ =NAD, $A_2$ =NH3, the ODE system is:

$$
\begin{aligned}
\frac{dm_0}{dt} &= -\frac{E_1 L_0 a_0 a_2 m_0 - F_1 K_1 a_1 m_1}{L_0 a_0 a_2 m_0 + K_1 a_1 m_1 + K_1 L_0} + \frac{E_2 L_1 a_1 m_1 - F_2 K_2 a_0 m_0 m_2}{K_2 a_0 m_0 m_2 + L_1 a_1 m_1 + K_2 L_1} \\
\frac{dm_1}{dt} &= \frac{E_1 L_0 a_0 a_2 m_0 - F_1 K_1 a_1 m_1}{L_0 a_0 a_2 m_0 + K_1 a_1 m_1 + K_1 L_0} - \frac{-F_2 K_2 a_0 m_0 m_2 + E_2 L_1 a_1 m_1}{K_2 a_0 m_0 m_2 + L_1 a_1 m_1 + K_2 L_1} \\
\frac{dm_2}{dt} &= \frac{-F_2 K_2 a_0 m_0 m_2 + E_2 L_1 a_1 m_1}{K_2 a_0 m_0 m_2 + L_1 a_1 m_1 + K_2 L_1} - k_{\text{out}} m_2 \\
\frac{da_0}{dt} &= -\frac{E_1 L_0 a_0 a_2 m_0 - F_1 K_1 a_1 m_1}{L_0 a_0 a_2 m_0 + K_1 a_1 m_1 + K_1 L_0} + \frac{-F_2 K_2 a_0 m_0 m_2 + E_2 L_1 a_1 m_1}{K_2 a_0 m_0 m_2 + L_1 a_1 m_1 + K_2 L_1} \\
\frac{da_1}{dt} &= \frac{E_1 L_0 a_0 a_2 m_0 - F_1 K_1 a_1 m_1}{L_0 a_0 a_2 m_0 + K_1 a_1 m_1 + K_1 L_0} - \frac{-F_2 K_2 a_0 m_0 m_2 + E_2 L_1 a_1 m_1}{K_2 a_0 m_0 m_2 + L_1 a_1 m_1 + K_2 L_1} \\
\frac{da_2}{dt} &= k_{\text{in}} - \frac{E_1 L_0 a_0 a_2 m_0 - F_1 K_1 a_1 m_1}{L_0 a_0 a_2 m_0 + K_1 a_1 m_1 + K_1 L_0}.
\end{aligned}
$$

There are three conservation laws:

$$a_0 + m_1 = M, \qquad a_0 + a_1 = A_{\text{tot}}, \qquad m_0 + a_1 = W.$$

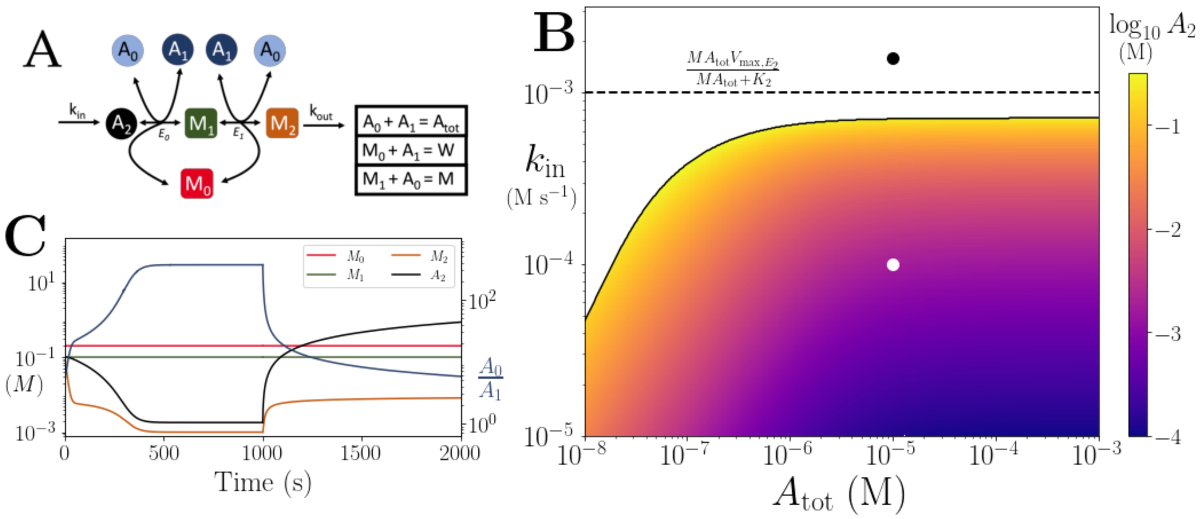

Figure S5: **(A)** Reaction system modelled in Sec. 7. **(B)** Heatmap of the steady state $A_2$ concentration. In the white area there is no steady state due to continual build up of $A_2$. Dashed line shows the limit $k_{\text{in}} < \frac{MTE_2}{MT+K_2}$, which is the smallest limit for these parameters. **(C)** Time series of $M_{0\to 2}$, $A_2$ and $A_0/A_1$. At $t = 1000$s, parameters are switched from the white dot in panel (B) (where a steady state exists) to the black dot (where there is continual build up of $A_2$). In (B) and (C) the other parameters are $k_{\text{cat}} = 100s^{-1}$, $E_{\text{tot}} = 0.01mM$, $K_m = 50\mu M$ and $k_{\text{out}} = 0.1s^{-1}$.

From $\frac{da_2}{dt} + \frac{dm_1}{dt} + \frac{dm_2}{dt} = 0$ we get

$$m_2 = \frac{k_{\text{in}}}{k_{\text{out}}}$$

as expected. From $\frac{dm_2}{dt} = 0$ and $\frac{da_2}{dt} = 0$ we get

$$m_1 = \frac{k_{\text{in}} K_2 \left(F_2 a_0 m_0 + k_{\text{in}} a_0 m_0 + L_1 k_{\text{out}}\right)}{k_{\text{out}} a_1 L_1 \left(E_2 - k_{\text{in}}\right)}, \qquad a_2 = \frac{K_1 \left(F_1 a_1 m_1 + a_1 k_{\text{in}} m_1 + L_0 k_{\text{in}}\right)}{m_0 a_0 L_0 \left(E_1 - k_{\text{in}}\right)}.$$

From the second and third conservation laws, we get

$$a_1 = A_{\text{tot}} - a_0, \qquad m_0 = W - a_1 = W - A_{\text{tot}} + a_0.$$

This gives that for a positive steady state, we need $k_{\text{in}} < E_1, k_{\text{in}} < E_2$ and $\max(A_{\text{tot}} - W, 0) < a_0 < A_{\text{tot}}$. Note that $A_{\text{tot}} - W$ is the constant value of $a_0 - m_0$ along trajectories.

Plugging these expressions into the first conservation law, we have that steady states are in one-to-one correspondence with the solutions to

$$M = a_0 + \frac{k_{\text{in}} K_2 \left(F_2 a_0 \left(a_0 - A_{\text{tot}} + W\right) + a_0 k_{\text{in}} \left(a_0 - A_{\text{tot}} + W\right) + L_1 k_{\text{out}}\right)}{k_{\text{out}} \left(-a_0 + A_{\text{tot}}\right) L_1 (E_2 - k_{\text{in}})}. \tag{7.2}$$

We first note that the derivative of the right hand side of (7.2) with respect to $a_0$ is:

$$1 + \frac{k_{\text{in}} K_2 \left(F_2 \left(a_0 - A_{\text{tot}} + W\right) + F_2 a_0 + k_{\text{in}} \left(a_0 - A_{\text{tot}} + W\right) + a_0 k_{\text{in}}\right)}{k_{\text{out}} \left(-a_0 + A_{\text{tot}}\right) L_1 (E_2 - k_{\text{in}})}$$
$$+ \frac{k_{\text{in}} K_2 \left(F_2 a_0 \left(a_0 - A_{\text{tot}} + W\right) + a_0 k_{\text{in}} \left(a_0 - A_{\text{tot}} + W\right) + L_1 k_{\text{out}}\right)}{k_{\text{out}} \left(-a_0 + A_{\text{tot}}\right)^2 L_1 (E_2 - k_{\text{in}})}$$

As $A_{\text{tot}} - a_0 > 0$ and $a_0 - A_{\text{tot}} - W > 0$, this function is positive in the interval of interest. Therefore, the right hand side of (7.2) is an increasing function of $a_0$ when $\max(A_{\text{tot}} - W, 0) < a_0 < A_{\text{tot}}$. It follows that if (7.2) has a solution, it has exactly one.

Rewriting (7.2) as a polynomial equation, steady states are in one-to-one correspondence with the roots in the interval $\max(A_{\text{tot}} - W, 0) < a_0 < A_{\text{tot}}$ of the following polynomial

$$p(a_0) = \left(-L_1 (E_2 - k_{\text{in}}) k_{\text{out}} + K_2 k_{\text{in}} (F_2 + k_{\text{in}})\right) a_0^2 + \left(L_1 (E_2 - k_{\text{in}}) k_{\text{out}} (M + T)\right.$$
$$\left. + (W - A_{\text{tot}}) K_2 k_{\text{in}} (F_2 + k_{\text{in}})\right) a_0 - L_1 k_{\text{out}} \left(M A_{\text{tot}} (E_2 - k_{\text{in}}) - K_2 k_{\text{in}}\right).$$

When $a_0 = A_{\text{tot}}$, the polynomial is positive.

**Case 1: $A_{\text{tot}} - W \leq 0$.** In this case we want $0 < a_0 < A_{\text{tot}}$. If the independent term of $p$ is negative, then $p$ has exactly one solution in $(0, A_{\text{tot}})$. So, if $MT(E_2 - k_{\text{in}}) - K_2 k_{\text{in}} < 0$, we have one positive steady state. This is equivalent to

$$k_{\text{in}} < \frac{M A_{\text{tot}} E_2}{M A_{\text{tot}} + K_2}.$$

If the independent term is positive or zero, then we note that the degree 1 term is also positive. So either $p$ has all coefficients positive and no positive roots, or the leading term is negative, in which case there is one root larger than $A_{\text{tot}}$. Therefore, no positive steady states in this case.

**Case 2: $A_{\text{tot}} - W > 0$.** In this case we want $A_{\text{tot}} - W < a_0 < A_{\text{tot}}$. We find that $p(A_{\text{tot}} - W)$ is

$$p(A_{\text{tot}} - W) = -L_1 k_{\text{out}} \left(W (M - A_{\text{tot}} + W)(E_2 - k_{\text{in}}) - K_2 k_{\text{in}}\right).$$

If this is negative, then there is one positive steady state, as $p$ is positive at $A_{\text{tot}}$. The condition is

$$k_{\text{in}} < \frac{W E_2 \left(M - A_{\text{tot}} + W\right)}{W (M - A_{\text{tot}} + W) + K_2}.$$

Note that $M + W - A_{\text{tot}} = a_0 + m_1 + m_0 + a_1 - a_0 - a_1 = m_0 + m_1$, and hence needs to be positive. This is assumed here.

If $p(A_{\text{tot}} - W) \geq 0$, then we are in the situation where $p$ is nonnegative both at $A_{\text{tot}} - W$ and $A_{\text{tot}}$, so, if there are roots in the interval $(A_{\text{tot}} - W, A_{\text{tot}})$, there must be two. This contradicts that we already showed that there was at most one. So, in this case, no steady states.

To summarize, there is one positive steady state if and only if $k_{\text{in}} < E_1$ and

$$k_{\text{in}} < \frac{M A_{\text{tot}} E_2}{M A_{\text{tot}} + K_2} \qquad \text{and} \qquad A_{\text{tot}} - W \leq 0,$$

or

$$k_{\text{in}} < \frac{W E_2 \left(M - A_{\text{tot}} + W\right)}{W (M - A_{\text{tot}} + W) + K_2}, \qquad A_{\text{tot}} - W > 0, \qquad \text{and} \qquad M + W - A_{\text{tot}} > 0.$$

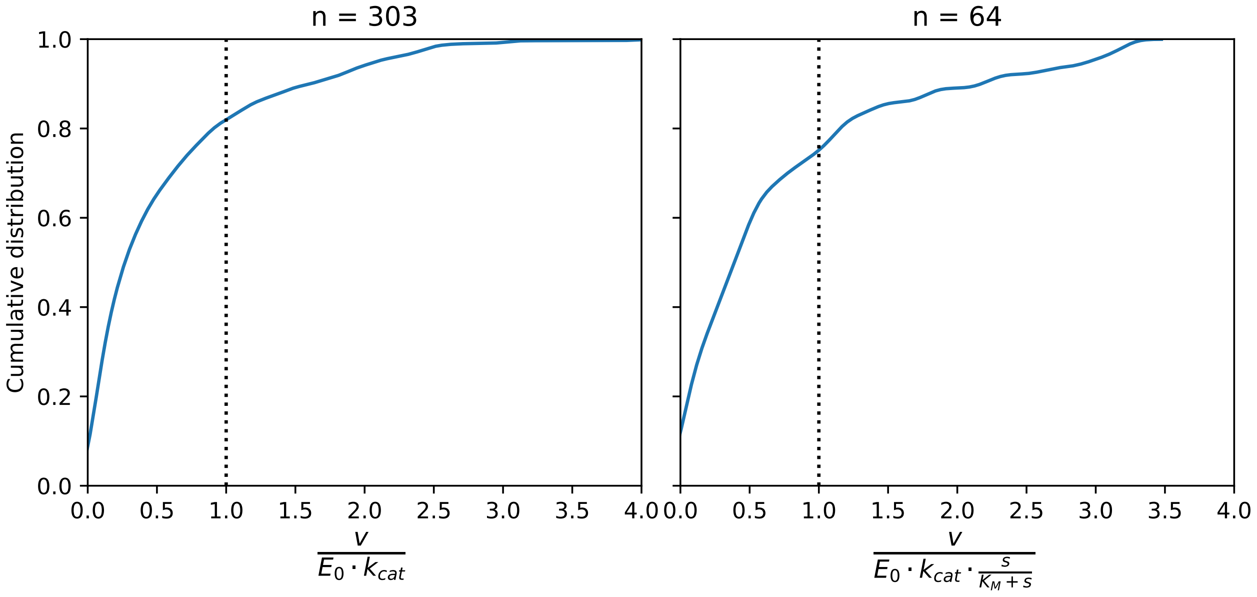

Figure S6: Cumulative distribution of the ratio of observed flux (measured or FBA-predicted) to enzyme kinetic based limit ($V_{\mathrm{max}}$) (left panel) or to enzyme kinetic based limit accounting for substrate affinity (right panel).

### 8 Analysis of existing flux data against predicted limits

To support the presented theory, we have analysed existing flux data - compiled from experiments and using flux balance analysis modelling - against predicted limits. The details and main results of this analysis are presented in the main text, under the results and methods sections. Here, we show the distribution of the ratio between collated flux data and flux limit based on enzyme kinetics (Fig. S6) and the correlation between co-substrate pool size and flux (Table S1).

### 9 Pathway branching into two pathways with independent co-substrates

We consider a scenario where two pathways share a common upstream metabolite, as shown in the motif in Fig. 4A in the main text. Each branch has its own conserved moiety that is cycled and all reactions are reversible. The reaction system is as follows:

$$\begin{array}{ccc} 0 \xrightarrow{k_{\mathrm{in}}} M_0 & M_0 \rightleftharpoons M_{1,2} & M_0 \rightleftharpoons M_{1,1} \\ M_{1,2} + A_0 \rightleftharpoons M_{2,2} + A_1 & M_{1,1} + B_0 \rightleftharpoons M_{2,1} + B_1 & A_1 \rightleftharpoons A_0 \\ B_1 \rightleftharpoons B_0 & M_{2,2} \xrightarrow{k_{\mathrm{out},2}} 0 & M_{2,1} \xrightarrow{k_{\mathrm{out},1}} 0. \end{array} \tag{9.1}$$

The ODE system is:

$$
\begin{aligned}
\frac{dm_0}{dt} &= k_{\text{in}} - \frac{E_{1,1}L_0m_0 - F_{1,1}K_{1,1}m_{1,1}}{K_{1,1}L_0 + K_{1,1}m_{1,1} + L_0m_0} - \frac{E_{1,2}L_0m_0 - F_{1,2}K_{1,2}m_{1,2}}{K_{1,2}L_0 + K_{1,2}m_{1,2} + L_0m_0} \\
\frac{dm_{1,2}}{dt} &= \frac{E_{1,2}L_0m_0 - F_{1,2}K_{1,2}m_{1,2}}{K_{1,2}L_0 + K_{1,2}m_{1,2} + L_0m_0} - \frac{E_{2,2}L_{1,2}m_{1,2}a_0 - F_{2,2}K_{2,2}m_{2,2}a_1}{K_{2,2}L_{1,2} + K_{2,2}m_{2,2}a_1 + L_{1,2}m_{1,2}a_0} \\
\frac{dm_{1,1}}{dt} &= \frac{E_{1,1}L_0m_0 - F_{1,1}K_{1,1}m_{1,1}}{K_{1,1}L_0 + K_{1,1}m_{1,1} + L_0m_0} - \frac{E_{2,1}L_{1,1}m_{1,1}b_0 - F_{2,1}K_{2,1}m_{2,1}b_1}{K_{2,1}L_{1,1} + K_{2,1}m_{2,1}b_1 + L_{1,1}m_{1,1}b_0} \\
\frac{dm_{2,2}}{dt} &= \frac{E_{2,2}L_{1,2}m_{1,2}a_0 - F_{2,2}K_{2,2}m_{2,2}a_1}{K_{2,2}L_{1,2} + K_{2,2}m_{2,2}a_1 + L_{1,2}m_{1,2}a_0} - m_{2,2}k_{\text{out},2} \\
\frac{dm_{2,1}}{dt} &= \frac{E_{2,1}L_{1,1}m_{1,1}b_0 - F_{2,1}K_{2,1}m_{2,1}b_1}{K_{2,1}L_{1,1} + K_{2,1}m_{2,1}b_1 + L_{1,1}m_{1,1}b_0} - m_{2,1}k_{\text{out},1} \\
\frac{da_0}{dt} &= -\frac{E_aL_aa_0 - F_aK_aa_1}{K_aL_a + K_aa_1 + L_aa_0} - \frac{E_{2,2}L_{1,2}m_{1,2}a_0 - F_{2,2}K_{2,2}m_{2,2}a_1}{K_{2,2}L_{1,2} + K_{2,2}m_{2,2}a_1 + L_{1,2}m_{1,2}a_0} \\
\frac{da_1}{dt} &= \frac{E_aL_aa_0 - F_aK_aa_1}{K_aL_a + K_aa_1 + L_aa_0} + \frac{E_{2,2}L_{1,2}m_{1,2}a_0 - F_{2,2}K_{2,2}m_{2,2}a_1}{K_{2,2}L_{1,2} + K_{2,2}m_{2,2}a_1 + L_{1,2}m_{1,2}a_0} \\
\frac{db_0}{dt} &= -\frac{E_bL_bb_0 - F_bK_bb_1}{K_bL_b + K_bb_1 + L_bb_0} - \frac{E_{2,1}L_{1,1}m_{1,1}b_0 - F_{2,1}K_{2,1}m_{2,1}b_1}{K_{2,1}L_{1,1} + K_{2,1}m_{2,1}b_1 + L_{1,1}m_{1,1}b_0} \\
\frac{db_1}{dt} &= \frac{E_bL_bb_0 - F_bK_bb_1}{K_bL_b + K_bb_1 + L_bb_0} + \frac{E_{2,1}L_{1,1}m_{1,1}b_0 - F_{2,1}K_{2,1}m_{2,1}b_1}{K_{2,1}L_{1,1} + K_{2,1}m_{2,1}b_1 + L_{1,1}m_{1,1}b_0}.
\end{aligned}
\tag{9.2}
$$

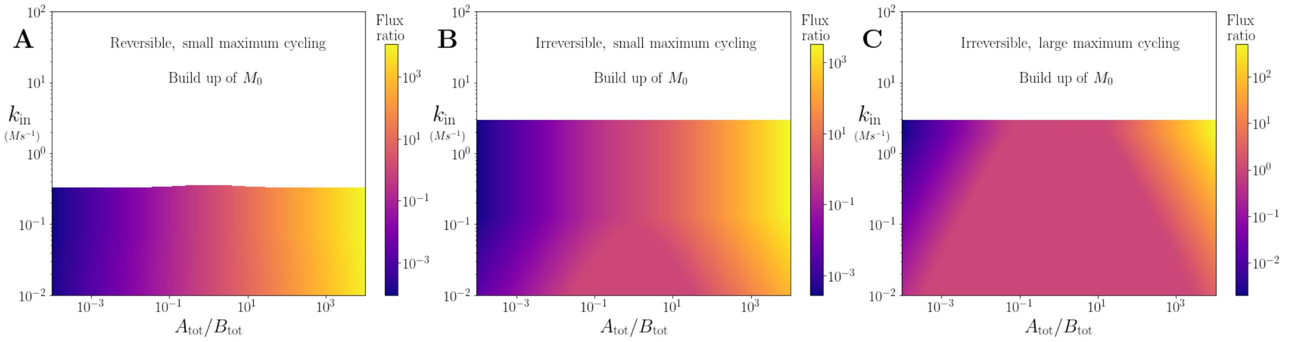

Figure S7: Effect of varying $A_{\text{tot}}/B_{\text{tot}}$ and $k_{\text{in}}$ in the branching system shown in Fig. 4A, on the ratio of flux into $M_{2,2}/M_{2,1}$ for reversible and irreversible dynamics and different $V_{\max,A}$ and $V_{\max,B}$ values. Downstream metabolite has higher flux when it's respective co-substrate has a higher concentration. **(A)** Reversible reactions with $V_{\max,A} = V_{\max,B} = 0.1$ (i.e. one order of magnitude smaller than the other reaction rates). **(B)** Irreversible reactions with $V_{\max,A} = V_{\max,B} = 0.1$ (i.e. one order of magnitude smaller than the other reaction rates). **(C)** Reversible reactions with $V_{\max,A} = V_{\max,B} = 10$ (i.e. one order of magnitude larger than the other reaction rates).

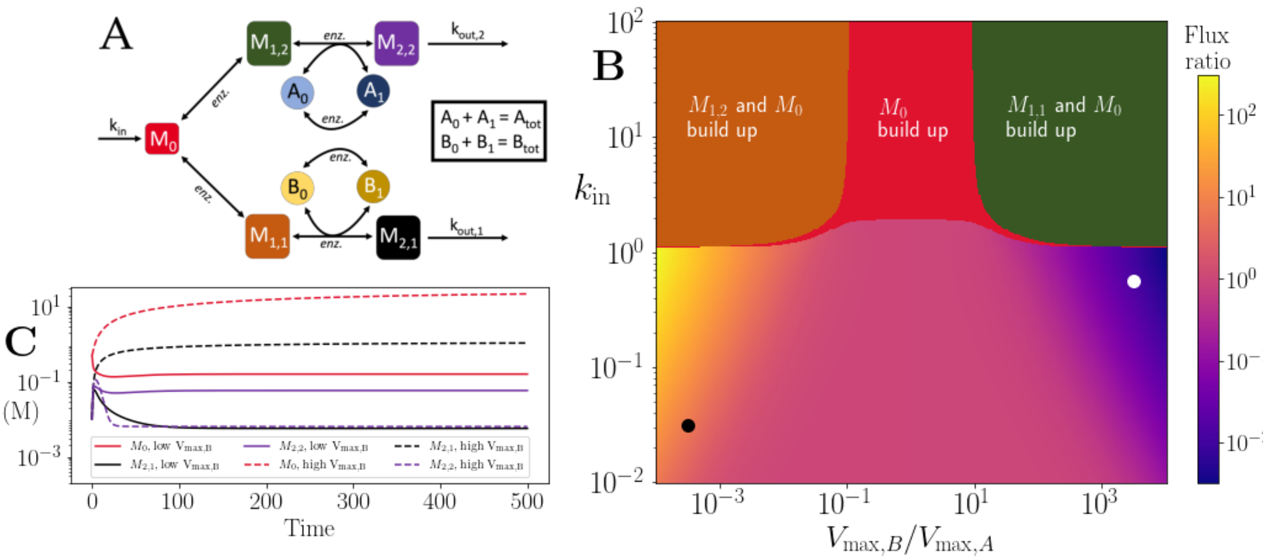

Figure S8: Effect of varying $V_{\mathrm{max},B}/V_{\mathrm{max},A}$ (i.e. maximum back-cycling) ratio and $k_{\mathrm{in}}$ in the branching system shown in Fig. 4A. **(A)** Motif showing the dynamics. **(B)** Heatmap showing the ratio of flux into $M_{2,2}/M_{2,1}$. Downstream metabolite has higher flux when the back-cycling rate of its co-substrate is higher, with a greater effect for higher $k_{\mathrm{in}}$ values. **(C)** Time series of downstream metabolites and shared precursor. Solid/dashed lines show parameters for black/white dot in (B).

While it is not possible to solve this system directly, we examine through simulations the effect of varying the influx of the shared upstream metabolite, $k_{\mathrm{in}}$ along with the ratio of the pool sizes, $A_{\mathrm{tot}}/B_{\mathrm{tot}}$ in the main text (Fig. 4), while varying $k_{\mathrm{in}}$ with the ratio of moiety back-cycling rates $k_a/k_b$ is presented in Fig S8.

#### 10 Independent pathways coupled by co-substrate cycling

We consider a scenario with two pathways that when isolated take the form examined in Sec. 2 of this document. However, we now suppose that they are coupled by the conserved moiety, *i.e.* $A_0 \to A_1$ in the 'forward' direction of the first pathway, while $A_1 \to A_0$ in the 'forward' direction of the other. The metabolite reactions are reversible, and there is pathway-independent cycling of the shared conserved moiety. The motif is shown in Fig. 5A in the main text, and the reactions are as follows:

$$\begin{aligned} &0 \xrightarrow{k_{\mathrm{in},1}} M_{0,1} \qquad M_{0,1} + A_0 \rightleftharpoons M_{1,1} + A_1 \qquad M_{1,1} \xrightarrow{k_{\mathrm{out},1}} 0 \\ &0 \xrightarrow{k_{\mathrm{in},2}} M_{0,2} \qquad M_{0,2} + A_1 \rightleftharpoons M_{1,2} + A_0 \qquad M_{1,2} \xrightarrow{k_{\mathrm{out},2}} 0 \\ &A_0 \rightleftharpoons A_1. \end{aligned} \tag{10.1}$$

The resulting system of ODEs is:

$$\begin{aligned}
\frac{dm_{0,1}}{dt} &= k_{\text{in},1} - \frac{E_{1,1}L_{0,1}m_{0,1}a_0 - F_{1,1}K_{1,1}m_{1,1}a_1}{K_{1,1}L_{0,1} + K_{1,1}m_{1,1}a_1 + L_{0,1}m_{0,1}a_0}\\
\frac{dm_{1,1}}{dt} &= \frac{E_{1,1}L_{0,1}m_{0,1}a_0 - F_{1,1}K_{1,1}m_{1,1}a_1}{K_{1,1}L_{0,1} + K_{1,1}m_{1,1}a_1 + L_{0,1}m_{0,1}a_0} - k_{\text{out},1}m_{1,1}\\
\frac{dm_{0,2}}{dt} &= k_{\text{in},2} - \frac{E_{1,2}L_{0,2}m_{0,2}a_1 - F_{1,2}K_{1,2}m_{1,2}a_0}{K_{1,2}L_{0,2} + K_{1,2}m_{1,2}a_0 + L_{0,2}m_{0,2}a_1}\\
\frac{dm_{1,2}}{dt} &= \frac{E_{1,2}L_{0,2}m_{0,2}a_1 - F_{1,2}K_{1,2}m_{1,2}a_0}{K_{1,2}L_{0,2} + K_{1,2}m_{1,2}a_0 + L_{0,2}m_{0,2}a_1} - k_{\text{out},2}m_{1,2}\\
\frac{da_0}{dt} &= \frac{E_{1,2}L_{0,2}m_{0,2}a_1 - F_{1,2}K_{1,2}m_{1,2}a_0}{K_{1,2}L_{0,2} + K_{1,2}m_{1,2}a_0 + L_{0,2}m_{0,2}a_1}\\
&\quad - \frac{E_{1,1}L_{0,1}m_{0,1}a_0 - F_{1,1}K_{1,1}m_{1,1}a_1}{K_{1,1}L_{0,1} + K_{1,1}m_{1,1}a_1 + L_{0,1}m_{0,1}a_0} - \frac{E_aL_aa_0 - F_aK_aa_1}{K_aL_a + K_aa_1 + L_aa_0}\\
\frac{da_1}{dt} &= -\frac{E_{1,2}L_{0,2}m_{0,2}a_1 - F_{1,2}K_{1,2}m_{1,2}a_0}{K_{1,2}L_{0,2} + K_{1,2}m_{1,2}a_0 + L_{0,2}m_{0,2}a_1}\\
&\quad + \frac{E_{1,1}L_{0,1}m_{0,1}a_0 - F_{1,1}K_{1,1}m_{1,1}a_1}{K_{1,1}L_{0,1} + K_{1,1}m_{1,1}a_1 + L_{0,1}m_{0,1}a_0} + \frac{E_aL_aa_0 - F_aK_aa_1}{K_aL_a + K_aa_1 + L_aa_0}.
\end{aligned} \tag{10.2}$$

This system has one conservation law

$$a_0 + a_1 = A_{\text{tot}},$$

giving that at steady state, $a_1 = A_{\text{tot}} - a_0$. By solving the steady state equations $\frac{dm_{0,1}}{dt} = \frac{dm_{1,1}}{dt} = \frac{dm_{0,2}}{dt} = \frac{dm_{1,2}}{dt} = 0$ for $m_{0,1}, m_{0,2}, m_{1,1}, m_{1,2}$, we obtain

$$\begin{aligned}
m_{0,1} &= \frac{K_{1,1}k_{\text{in},1}\left((F_{1,1} + k_{\text{in},1})a_1 + L_{0,1}k_{\text{out},1}\right)}{k_{\text{out},1}L_{0,1}a_0\left(E_{1,1} - k_{\text{in},1}\right)}, & m_{1,1} &= \frac{k_{\text{in},1}}{k_{\text{out},1}},\\
m_{0,2} &= \frac{K_{1,2}k_{\text{in},2}((F_{1,2} + k_{\text{in},2})a_0 + L_{0,2}k_{\text{out},2})}{k_{\text{out},2}L_{0,2}a_1\left(E_{1,2} - k_{\text{in},2}\right)}, & m_{1,2} &= \frac{k_{\text{in},2}}{k_{\text{out},2}}.
\end{aligned}$$

The expressions are positive provided $E_{1,1} > k_{\text{in},1}$, $E_{1,2} > k_{\text{in},2}$, and $a_0, a_1 > 0$.

Finally, we use $\frac{da_0}{dt} = 0$ to solve for $a_0$ and obtain

$$a_0 = \frac{K_a\left((k_{\text{in},2} - k_{\text{in},1})(L_a + A_{\text{tot}}) + A_{\text{tot}}F_a\right)}{(k_{\text{in},2} - k_{\text{in},1})(K_a - L_a) + E_aL_a + F_aK_a}. \tag{10.3}$$

For all quantities to be positive at steady state, we require $0 < a_0 < A_{\text{tot}}$. By subtracting from $A_{\text{tot}}$ the value of $a_0$ at steady state, we obtain

$$A_{\text{tot}} - a_0 = L_a\big((A_{\text{tot}} + K_a)(k_{\text{in},1} - k_{\text{in},2}) + A_{\text{tot}}E_a\big). \tag{10.4}$$

To summarize, there is a positive steady state if and only if $E_{1,1} > k_{\text{in},1}$, $E_{1,2} > k_{\text{in},2}$, and (10.4) and (10.3) are positive. This leads to (10.4) positive, that is,

$$\frac{A_{\text{tot}}E_a}{A_{\text{tot}} + K_a} > k_{\text{in},2} - k_{\text{in},1}, \qquad \text{that is} \qquad -\frac{A_{\text{tot}}E_a}{A_{\text{tot}} + K_a} < k_{\text{in},1} - k_{\text{in},2}, \tag{10.5}$$

and either

$$\frac{A_{\text{tot}}F_a}{A_{\text{tot}} + L_a} > k_{\text{in},1} - k_{\text{in},2}, \qquad E_aL_a + F_aK_a > (k_{\text{in},1} - k_{\text{in},2})(K_a - L_a) \tag{10.6}$$

or

$$\frac{A_{\text{tot}}F_a}{A_{\text{tot}} + L_a} < k_{\text{in},1} - k_{\text{in},2}, \qquad E_aL_a + F_aK_a < (k_{\text{in},1} - k_{\text{in},2})(K_a - L_a) \tag{10.7}$$

By discussing cases, we obtain all possible scenarios for a positive steady state to exist, and these are dictated by the difference $k_{\text{in},1} - k_{\text{in},2}$.

For example, if $k_{\text{in},1} = k_{\text{in},2}$, then (10.5) and (10.6) hold directly. If $K_a = L_a$, then we require $\frac{A_{\text{tot}}F_a}{A_{\text{tot}}+L_a} > k_{\text{in},1} - k_{\text{in},2} > -\frac{A_{\text{tot}}E_a}{A_{\text{tot}}+K_a}$ to hold.

If $K_a > L_a$, the conditions lead to either

$$\min\left(\frac{A_{\text{tot}}F_a}{A_{\text{tot}}+L_a}, \frac{E_aL_a+F_aK_a}{(K_a-L_a)}\right) > k_{\text{in},1} - k_{\text{in},2} > -\frac{A_{\text{tot}}E_a}{A_{\text{tot}}+K_a}$$

or

$$\max\left(\frac{A_{\text{tot}}F_a}{A_{\text{tot}}+L_a}, \frac{E_aL_a+F_aK_a}{(K_a-L_a)}, -\frac{A_{\text{tot}}E_a}{A_{\text{tot}}+K_a}\right) < k_{\text{in},1} - k_{\text{in},2}.$$

If $K_a < L_a$, the conditions lead to either

$$\frac{A_{\text{tot}}F_a}{A_{\text{tot}}+L_a} > k_{\text{in},1} - k_{\text{in},2} > \max\left(-\frac{A_{\text{tot}}E_a}{A_{\text{tot}}+K_a}, \frac{E_aL_a+F_aK_a}{(K_a-L_a)}\right)$$

or

$$\max\left(\frac{A_{\text{tot}}F_a}{A_{\text{tot}}+L_a}, -\frac{A_{\text{tot}}E_a}{A_{\text{tot}}+K_a}\right) < k_{\text{in},1} - k_{\text{in},2} < \frac{E_aL_a+F_aK_a}{(K_a-L_a)}.$$

A key consequence of these conditions is that coupled pathways can admit higher influx values without upstream metabolite build up, due to the cycling enzyme condition now depending on the difference between the $k_{\text{in}}$ values rather than the values themselves. This occurs when the limit due to the pathway enzyme is larger than that of the cycling enzyme, so there is always a range of $A_{\text{tot}}$ where this applies. An example is shown in Fig. S9.

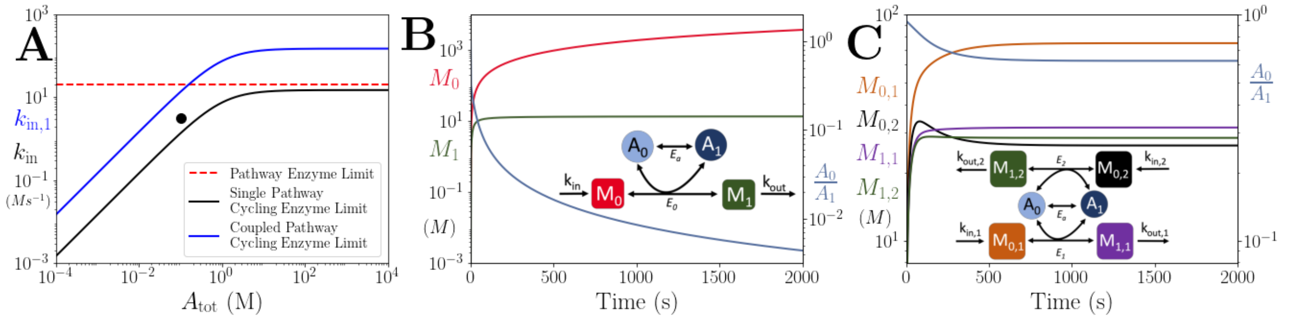

Figure S9: Comparison of the onset of instability in the coupled pathway case with the single pathway case. **(A)** Pathway enzyme and cycling enzyme limits in the single and coupled pathway cases (see legend, pathway enzyme limit is the same for each case). For the coupled pathway, $k_{\text{in},2}$ is set to $0.9k_{\text{in},1}$. Since the limiting factor depends on the difference between $k_{\text{in},1}$ and $k_{\text{in},2}$, higher $k_{\text{in}}$ values are possible without build up of upstream metabolites if all other parameters are kept the same. **(B)** Time series of $M_0$, $M_1$ and $A_0/A_1$ in the single pathway case where the parameters for $A_{\text{tot}}$ and $k_{\text{in}}$ are set by the black dot in (A). Here we see build up $M_0$ because $k_{\text{in}} > A_{\text{tot}}F_a/(A_{\text{tot}} + L_a)$. **(C)** Time series of $M_{0,1}$, $M_{0,2}$, $M_{1,1}$, $M_{1,2}$ and $A_0/A_1$ in the coupled pathway case where the parameters for $A_{\text{tot}}$, $k_{\text{in},1}$ and $k_{\text{in},2}$ are set by the black dot in (A). Here we see the system admits a steady state because $k_{\text{in},1} - k_{\text{in},2} < A_{\text{tot}}F_a/(A_{\text{tot}} + L_a)$. The parameters in all panels are: $E_1 = F_1 = E_{1,1} = E_{1,2} = F_{1,1} = F_{1,2} = 20\ Ms^{-1}$, $K_1 = L_0 = K_{1,1} = K_{1,2} = L_{0,1} = L_{0,2} = 1\ M$, $k_{\text{out}} = k_{\text{out},1} = k_{\text{out},2} = 0.1\ s^{-1}$, $E_a = F_a = 15\ Ms^{-1}$, $K_a = L_a = 1\ M$.

To complement this steady state analysis with dynamics of this system, we use numerical simulations to study the system. In particular, we consider the effect of randomly fluctuating influxes on the downstream metabolites. The analysis is achieved by fixing the average of the influxes, while drawing the log-ratio (*i.e.* $\log_{10} k_{\text{in},1}/k_{\text{in},2}$) from a standard normal distribution with mean $\mu = 0$ and variance $\sigma^2 = 1$. A new log-ratio is drawn after waiting a time that is drawn from an exponential distribution with mean $\tau$. Example time-series are shown in Fig. 5(C,D) of the main text. The log-ratio is chosen as the variable instead of simply the ratio as it allows us to examine large variations, while keeping the effect on each pathway symmetric. This random process can be thought of as the discrete-time analogue of the Ornstein-Uhlenbeck process: it has the same steady state distribution, mean, variance and correlation function as its continuous-time counterpart, while being much easier and faster to implement as part of a larger system where the other equations are deterministic.

The effect of different values of $\tau$ and total pool size $A_{\text{tot}}$ is shown in Fig. 5 of the main text, and is further explored in Fig. S10. These analyses show that the effect of increasing the pathway independent moiety cycling rate is to reduce the critical pool size above which the downstream metabolites are anti-correlated. Furthermore, this behaviour is observed whether the metabolite reactions are irreversible or reversible. Since the transition from correlated to anti-correlated is so sharp, relatively small changes in the total amount of cycled co-substrate, combined with noisy influx rates, could lead to the downstream metabolites changing from being correlated to anti-correlated, or vice-versa.

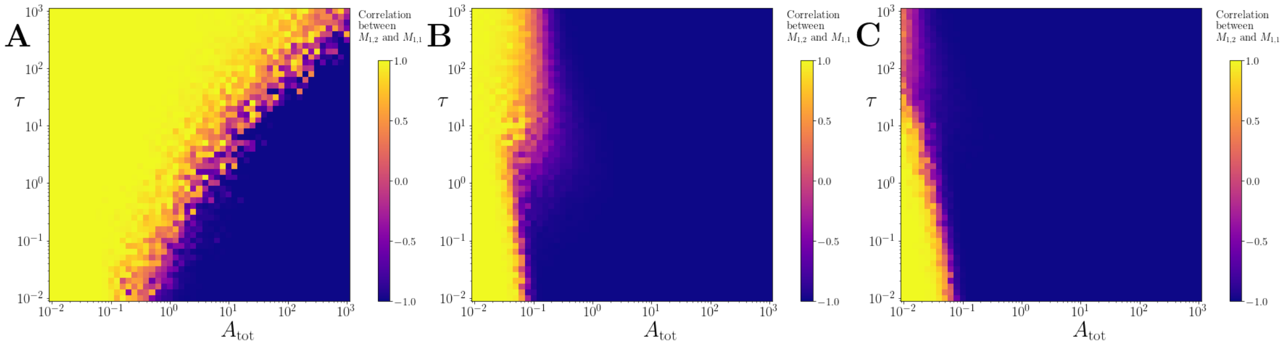

Figure S10: Correlation between products of two coupled pathways, where the metabolite reactions are irreversible. (A), (B) and (C) show results for $V_{\max,E_a} = 0.01, 1$ and 10 respectively. As in the Fig. 5 of the main text, all other parameters are set to 1, apart from the $k_{\text{out}}$ values that are each 0.5. As $V_{\max,E_a}$ increases, the products become anti-correlated at smaller $A_{\text{tot}}$ values. This is also true for the reversible case.

| | reaction | flux source | number of conditions | pearson-r | p-value |
|---|---|---|---|---|---|
| 25 | MDH | Gerosa | 7 | 0.874310 | 0.010042 |
| 33 | PGK | Gerosa | 4 | 0.973547 | 0.026453 |
| 0 | 3OAR140 | Davidi | 7 | 0.794377 | 0.032850 |
| 7 | DHFS | Davidi | 7 | 0.787719 | 0.035438 |
| 46 | UAMAS | Davidi | 7 | 0.783767 | 0.037025 |
| 45 | UAMAGS | Davidi | 7 | 0.783767 | 0.037025 |
| 44 | UAAGDS | Davidi | 7 | 0.783767 | 0.037025 |
| 48 | UGMDDS | Davidi | 7 | 0.783767 | 0.037025 |
| 2 | AIRC2 | Davidi | 7 | 0.783523 | 0.037125 |
| 39 | PRASCSi | Davidi | 7 | 0.783523 | 0.037125 |
| 27 | NADK | Davidi | 7 | 0.783523 | 0.037125 |
| 19 | HSK | Davidi | 7 | 0.783042 | 0.037321 |
| 4 | CTPS2 | Davidi | 7 | 0.782303 | 0.037624 |
| 41 | PTPATi | Davidi | 7 | 0.781795 | 0.037833 |
| 35 | PNTK | Davidi | 7 | 0.781795 | 0.037833 |
| 8 | DTMPK | Davidi | 7 | 0.780177 | 0.038501 |
| 43 | TMPK | Davidi | 7 | 0.777454 | 0.039643 |
| 34 | PMPK | Davidi | 7 | 0.777454 | 0.039643 |
| 1 | ADSK | Davidi | 7 | 0.777418 | 0.039658 |
| 15 | GLU5K | Davidi | 7 | 0.774461 | 0.040918 |
| 37 | PRAGSr | Davidi | 7 | 0.773900 | 0.041159 |
| 40 | PRFGS | Davidi | 7 | 0.773900 | 0.041159 |
| 38 | PRAIS | Davidi | 7 | 0.773900 | 0.041159 |
| 14 | GK1 | Davidi | 7 | 0.772918 | 0.041584 |
| 6 | DBTS | Davidi | 7 | 0.771900 | 0.042026 |
| 5 | CYTK1 | Davidi | 7 | 0.762125 | 0.046410 |
| 20 | ICDHyr | Davidi | 7 | 0.743003 | 0.055681 |
| 47 | UAPGR | Davidi | 7 | 0.702799 | 0.078196 |
| 9 | G5SD | Davidi | 7 | 0.692830 | 0.084415 |
| 28 | P5CR | Davidi | 7 | 0.692830 | 0.084415 |
| 12 | GAPD | Davidi | 7 | -0.681203 | 0.091988 |
| 11 | G6PDH2r | Gerosa | 7 | 0.629982 | 0.129423 |
| 10 | G6PDH2r | Davidi | 7 | 0.617909 | 0.139204 |
| 16 | GND | Davidi | 7 | 0.617909 | 0.139204 |
| 22 | IMPD | Davidi | 7 | -0.539918 | 0.210941 |
| 23 | IPMD | Davidi | 7 | -0.535973 | 0.214953 |
| 36 | PPND | Davidi | 7 | -0.520516 | 0.231016 |
| 26 | MTHFR2 | Davidi | 7 | -0.519049 | 0.232569 |
| 32 | PGCD | Davidi | 7 | -0.517475 | 0.234241 |
| 3 | AKGDH | Gerosa | 7 | 0.498666 | 0.254643 |
| 17 | GND | Gerosa | 7 | 0.498636 | 0.254676 |
| 42 | PYK | Gerosa | 4 | -0.718821 | 0.281179 |
| 18 | HISTD | Davidi | 7 | -0.469323 | 0.288020 |
| 30 | PFK | Davidi | 5 | 0.537260 | 0.350444 |
| 24 | MDH | Davidi | 7 | 0.311471 | 0.496502 |
| 13 | GAPD | Gerosa | 4 | -0.451817 | 0.548183 |
| 21 | ICDHyr | Gerosa | 7 | -0.169510 | 0.716347 |
| 29 | PDH | Gerosa | 7 | 0.098015 | 0.834403 |
| 31 | PFK | Gerosa | 2 | nan | nan |

Table 1: Correlation coefficients between observed flux (measured or FBA-predicted) and co-substrate pool size. For reaction ID and descriptions, see Supplementary File 1.
