## Supplementary File 2 for "Co-substrate pools can constrain and regulate pathway fluxes in cell metabolism": figure_4_branched_pathway_reversible.py

```
#!/usr/bin/env python3
# -*- coding: utf-8 -*-
"""
Created on Wed Mar  9 14:51:47 2022

@author: robert
"""


import numpy as np                      ### standard math library
import matplotlib.pyplot as plt         ### for plotting
from scipy.integrate import odeint      ### for integrating the ODES
import matplotlib as mpl                ### for global matplotlib settings
import time                             ### for timing the code
from matplotlib.colors import ListedColormap  ### for setting colormap colors
import numba                            ### precompiles numpy functions for speed


#### for using latex in the figures ####
### note - if you get an error about latex, comment out this, the x and y axis 
### labels and the title in the heatmap - lines 326 - 328
mpl.rc('text', usetex = True)
timer0=time.perf_counter()    

### PARAMETERS TO CHANGE IN THE HEATMAP

###     backflow is V_{max, E_a}  -   change to see different behaviour - 
###     reducing will introduce black area where there is build up
###     of a m0, m11 and m12
backflow=10.

###  mean of Atot and Btot, 
cosub_avg=1.

###     number of kin and atot values for the heatmap
###     larger values give a higher resolution but take longer. Run initially with 
###     these values, then check the output for how long it took before you increase.
###     time scales linearly with both numkin and numAtotinit (so increasing both by a 
###     factor of x will increase the time by x^2                                                       
numkin=9
numAtotinit=9


### PARAMETERS TO CHANGE IN THE TIME SERIES

### choose kin and cosub ratio values for the dots and time series
kin1=10**(-1)
kin2=10**(-0.75)
cosub_ratio1=10**-3
cosub_ratio2=10**2

#### defin ecolors for use in the figures later
m0col='#e0132f'
m21col='#000000'
m22col='#7030a0'
m12col='#385723'
m11col='#C55A11'


#### set parameters
E11=1.
E12=1.
E21=1.
E22=1.

L0=1.
L11=1.
L12=1.

F11=1.
F12=1.
F21=1.
F22=1.

K11=1.
K12=1.
K21=1.
K22=1.


kout1=0.5
kout2=0.5        


Ea=backflow
Fa=backflow
Ka=1.
La=1.

Eb=backflow
Fb=backflow
Kb=1.
Lb=1.


### Inital values for metabolites
M0_init=0.5
M11_init=0.1
M12_init=0.1
M21_init=0.01
M22_init=0.01


### define functions for the ODES ####

## michaelis - menten with no co-sub
@numba.njit
def nu(m1,m2,e2,f2,k2,l1):
    return (e2*l1*m1 - f2*k2*m2)/(k2*l1+k2*m2+l1*m1)

## michaelis - menten with co-sub
@numba.njit
def mu(m1,m2,s0,s1,e2,f2,k2,l1):
    return (e2*l1*m1*s0 - f2*k2*m2*s1)/(k2*l1+k2*m2*s1+l1*m1*s0)


####  ODES for m0, m11, m12, m21, m22, a0, a1, b0 and b1 ####
def dm0_dt(P,t):
    m0=P[0]
    m11=P[1]
    m12=P[2]
    return kin - nu(m0,m11,E11,F11,K11,L0)- nu(m0,m12,E12,F12,K12,L0)

def dm11_dt(P,t):
    m0=P[0]
    m11=P[1]
    m21=P[3]
    b0=P[7]
    b1=P[8]
    return nu(m0,m11,E11,F11,K11,L0)- mu(m11,m21,b0,b1,E21,F21,K21,L11)

def dm12_dt(P,t):
    m0=P[0]
    m12=P[2]
    m22=P[4]
    a0=P[5]
    a1=P[6]
    return nu(m0,m12,E12,F12,K12,L0)- mu(m12,m22,a0,a1,E22,F22,K22,L12)

def dm21_dt(P,t):
    m11=P[1]
    m21=P[3]
    b0=P[7]
    b1=P[8]
    return mu(m11,m21,b0,b1,E21,F21,K21,L11) - m21*kout1

def dm22_dt(P,t):
    m12=P[2]
    m22=P[4]
    a0=P[5]
    a1=P[6]
    return mu(m12,m22,a0,a1,E22,F22,K22,L12) - m22*kout2

def da0_dt(P,t):
    m12=P[2]
    m22=P[4]
    a0=P[5]
    a1=P[6]
    return -mu(a0,a1,1.,1.,Ea,Fa,Ka,La)- mu(m12,m22,a0,a1,E22,F22,K22,L12)

def da1_dt(P,t):
    m12=P[2]
    m22=P[4]
    a0=P[5]
    a1=P[6]
    return mu(m12,m22,a0,a1,E22,F22,K22,L12)+mu(a0,a1,1.,1.,Ea,Fa,Ka,La)

def db0_dt(P,t):
    m11=P[1]
    m21=P[3]
    b0=P[7]
    b1=P[8]
    return -mu(b0,b1,1.,1.,Eb,Fb,Kb,Lb) -  mu(m11,m21,b0,b1,E21,F21,K21,L11)

def db1_dt(P,t):
    m11=P[1]
    m21=P[3]
    b0=P[7]
    b1=P[8]
    return  mu(m11,m21,b0,b1,E21,F21,K21,L11) +mu(b0,b1,1.,1.,Eb,Fb,Kb,Lb)


#### general function that collects the above into a vector
def dP_dt(P,t):
    return [dm0_dt(P,t), dm11_dt(P,t), dm12_dt(P,t), dm21_dt(P,t), dm22_dt(P,t), 
            da0_dt(P,t), da1_dt(P,t),db0_dt(P,t), db1_dt(P,t)]


####  functions that calculate the flux from 12 to 22 and 11 to 21
def m12tom22flux(P):
    m12=P[2]
    m22=P[4]
    a0=P[5]
    a1=P[6]
    return E22*L12*m12*a0/(K22*L12+K22*m22*a1+L12*m12*a0)

def m11to21flux(P):
    m11=P[1]
    m21=P[3]
    b0=P[7]
    b1=P[8]
    return E21*L11*m11*b0/(K21*L11+K21*m21*b1+L11*m11*b0)


### generate the kin values and then the corners for the heatmap ###
kin_vals=np.logspace(-2,2,numkin)
intervalkin=np.log10(kin_vals[1]/kin_vals[0])
kin_valsplot=np.logspace(-2-intervalkin/2.,2+intervalkin/2,numkin+1)

### generate the Atot and Btot values and then the corners for the heatmap ###
### first define the ratios ###
cosub_ratios = np.logspace(-4,4,numAtotinit)
### calculate the pool sizes so that the average is constant ####
Bpool_vals= 2*cosub_avg/(1+cosub_ratios)
Apool_vals=cosub_ratios*Bpool_vals
interval_cosubratios=np.log10(cosub_ratios[1]/cosub_ratios[0])
cosub_ratios_plot=np.logspace(-4-interval_cosubratios/2,4+interval_cosubratios/2,numAtotinit+1)


#### initialise matrices that will be used in the heatmap
M21M22fluxratios_vals=np.empty((len(kin_vals),len(cosub_ratios)))
M21M22fluxratios_vals[:]=np.NaN

M12buildup=np.empty((len(kin_vals),len(cosub_ratios)))
M12buildup[:]=np.NaN

M11buildup=np.empty((len(kin_vals),len(cosub_ratios)))
M11buildup[:]=np.NaN

M0buildup=np.empty((len(kin_vals),len(cosub_ratios)))
M0buildup[:]=np.NaN

allbuildup=np.empty((len(kin_vals),len(cosub_ratios)))
allbuildup[:]=np.NaN


####  loop over kin
for i in range(len(kin_vals)):
    kin=kin_vals[i]
    
    #### loop over cosub ratios
    for ii in range(len(cosub_ratios)):
        
        #### get initial values of cosubstrates
        A0_init=Apool_vals[ii]/2.
        A1_init=Apool_vals[ii]/2.
        B0_init=Bpool_vals[ii]/2
        B1_init=Bpool_vals[ii]/2

        ### initialise vector for ODE
        P0=[M0_init, M11_init, M12_init, M21_init, M22_init, A0_init, A1_init, B0_init, B1_init]
        numtpts=10**4
        ts=np.linspace(0,numtpts,1000)
        ### integrate the ODE system
        Ps = odeint(dP_dt, P0, ts)
        
        
        
        M0=Ps[:,0]
        M11=Ps[:,1]
        M12=Ps[:,2]
        M21=Ps[:,3]
        M22=Ps[:,4]
        A0=Ps[:,5]
        A1=Ps[:,6]
        B0=Ps[:,7]
        B1=Ps[:,8]
        

        Pfinal=Ps[-1,:]        
        
        
        ### get the final flux values into the end products
        m12fluxfinal=m12tom22flux(Pfinal)
        m11fluxfinal=m11to21flux(Pfinal)
        
        ### calculate the final gradients for the precursor metabolites
        dm0dtfinal=dm0_dt(Pfinal,0)
        dm11dtfinal=dm11_dt(Pfinal,0)
        dm12dtfinal=dm12_dt(Pfinal,0)
        
        ### set the tolerance - if the gradient is below this value then we say 
        ### the relevant variable has reached a constant value
        tol=10**-1
        
        ### find out which metabolites have reached a constant value, or are 
        ### continuing to build up - note that if m0 is constant then all others are too
        if abs(dm0dtfinal)<tol:
            M21M22fluxratios_vals[i,ii]=m12fluxfinal/m11fluxfinal
        elif abs(dm11dtfinal)<tol and abs(dm12dtfinal)<tol:
            M0buildup[i,ii]=1
        elif abs(dm11dtfinal)<tol:
            M12buildup[i,ii]=1
        elif abs(dm12dtfinal)<tol:
            M11buildup[i,ii]=1
        else:
            allbuildup[i,ii]=1

    ### keep track of how far through generating the heatmap you are    
    print( round(i*100/len(kin_vals),0),'% done')

### calculte the time taken to generate the heatmap
timer1=time.perf_counter()    
print(f"time for runs was {(timer1-timer0)/60: 0.4f} minutes")

### initialise the figure  
plt.figure()
plt.axes([0.05,0.05,0.95,0.9])

### plot the heatmap of flux ratios - only for values where there is no build up
plt.pcolormesh(cosub_ratios_plot,kin_valsplot,np.log10(M21M22fluxratios_vals),
               cmap='plasma')#,shading='gouraud')

plt.yscale('log')
plt.xscale('log')

plt.xlabel('$[A]/[B]$',fontsize=20)
plt.ylabel('$k_{\\rm in}$',fontsize=20,rotation=0)
plt.title('flux into $M_{22}$/flux into $M_{21}$')
cb=plt.colorbar()


### show block colors where there is build up of the relevant metabolite(s)
cmap1=ListedColormap(m12col)
plt.pcolormesh(cosub_ratios_plot,kin_valsplot,(M12buildup),cmap=cmap1)#,alpha=0.3)#,shading='gouraud')
plt.yscale('log')
plt.xscale('log')


cmap2=ListedColormap(m11col)
plt.pcolormesh(cosub_ratios_plot,kin_valsplot,(M11buildup),cmap=cmap2)#,alpha=0.3)#,shading='gouraud')
plt.yscale('log')
plt.xscale('log')

cmap3=ListedColormap(m0col)
plt.pcolormesh(cosub_ratios_plot,kin_valsplot,(M0buildup),cmap=cmap3)#,alpha=0.3)#,shading='gouraud')
plt.yscale('log')
plt.xscale('log')

colarray4=[[0,0,0]]
cmap4=ListedColormap(colarray4)
plt.pcolormesh(cosub_ratios_plot,kin_valsplot,(allbuildup),cmap=cmap4)#,alpha=0.3)#,shading='gouraud')
plt.yscale('log')
plt.xscale('log')


Bpool1= 2*cosub_avg/(1+cosub_ratio1)
Apool1=cosub_ratio1*Bpool1

Bpool2= 2*cosub_avg/(1+cosub_ratio2)
Apool2=cosub_ratio2*Bpool2


plt.plot(Apool1/Bpool1,kin1,'o',markerfacecolor='w',markeredgecolor='w',markersize=6)
plt.plot(Apool2/Bpool2,kin2,'o',markerfacecolor='k',markeredgecolor='k',markersize=6)


### generate the two time series
P0=[M0_init, M11_init, M12_init, M21_init, M22_init, Apool1/2, Apool1/2, Bpool1/2, Bpool1/2]
P0_2=[M0_init, M11_init, M12_init, M21_init, M22_init, Apool2/2, Apool2/2, Bpool2/2, Bpool2/2]


timeint=5*10**2

ts=np.linspace(0,timeint,1000)
kin=kin1
Ps = odeint(dP_dt, P0, ts)

M0=Ps[:,0]
M11=Ps[:,1]
M12=Ps[:,2]
M21=Ps[:,3]
M22=Ps[:,4]
A0=Ps[:,5]
A1=Ps[:,6]
B0=Ps[:,7]
B1=Ps[:,8]

kin=kin2
Ps_new = odeint(dP_dt, P0_2, ts)

M0_new=Ps_new[:,0]
M11_new=Ps_new[:,1]
M12_new=Ps_new[:,2]
M21_new=Ps_new[:,3]
M22_new=Ps_new[:,4]
A0_new=Ps_new[:,5]
A1_new=Ps_new[:,6]
B0_new=Ps_new[:,7]
B1_new=Ps_new[:,8]


### plot the time series of m0, m21 and m22 for the values specified above
plt.axes([0.0,-0.65,0.9,0.5])
plt.title('Concentrations over time, white dot')
plt.yscale('log')

plt.plot(ts,M0,color=m0col)
plt.plot(ts,M21,'-',color=m21col)
plt.plot(ts,M22,'-',color=m22col)

plt.plot(ts,M0_new,'--',color=m0col)#,label='$M_0$')
plt.plot(ts,M21_new,'--',color=m21col)#,label='B product')
plt.plot(ts,M22_new,'--',color=m22col)#,label='A product')
```
