## Supplementary File 3 for "Co-substrate pools can constrain and regulate pathway fluxes in cell metabolism": Figure_5_coupled_pathway_reversible.py

```
#!/usr/bin/env python3
# -*- coding: utf-8 -*-
"""
Created on Fri Apr 15 16:35:27 2022

@author: robert
"""

import numpy as np
import matplotlib.pyplot as plt
from scipy.integrate import odeint
import numba
import time


### heatmap takes a while to generate - to skip this set the following variable
### to 0, to include change back to 1
wantheatmap =1

### PARAMETERS TO CHANGE IN THE HEATMAP

###     bf is V_{max, E_a}  -   change to see different behaviour - 
###     increasing lowers the threshold value of Atot to anti-correlation
###     kav is the mean of kin,1 and kin,2 - increasing leads to breakdown of
###     transition between correlated and anti-correlated becuase both end up above the threshold
###     value and there is build up in all cases
###     var is the variance of the log kin ratios - this has not been extensively 
###     examined - please play around with it!
kav=0.1
bf=0.01
var=1.

###     number of tau and atot values for the heatmap
###     larger values give a higher resolution but take longer. Run initially with 
###     these values, then check the output for how long it took before you increase.
###     time scales linearly with both numkin and numAtotinit (so increasing both by a 
###     factor of x will increase the time by x^2  - with the current parameters 
###     it takes around 10 minutes                                                     
numtauvals=9
numAvals=9
taumax=2 ## max value of log10(tau) - increasing leads to much longer simulation times
atotmax=2 ##max value to log10(Atot) - can increase without time penalty

### PARAMETERS TO CHANGE IN THE TIME SERIES
### tau_ts is the correlation time of the noise, Atot_ts is the cosub concentration
tau_ts=10**(2)
Atot_ts=10**-2

#### define colors for use in the figures later
m01col='#c55a11'
m02col='k'
m12col='#385723'
m11col='#7030a0'
a1col=[(32.+143)/510.,(56.+170)/510.,(100.+220)/510. ]
k_ratio_col='#ecb7c7'

### set parameters
E11=1.
E12=1.
L01=1.
L02=1.
F11=1.
F12=1.
K11=1.
K12=1.

Ka=1.
La=1.

### Ea and Fa are the Vmax_Ea values in both directions
Ea=bf
Fa=bf

kout1=0.5
kout2=0.5

### Inital values for metabolites
M01_init=0.1
M02_init=0.1
M11_init=0.01
M12_init=0.01


### define functions for the ODES ####

### get the kin values 
@numba.njit
def get_kratio_vals(var,kav):
    kratio_log=np.random.normal(0,np.sqrt(var))
    kratio=10**(kratio_log)
    
    kin1=2*kav/(1+kratio)
    kin2=kratio*kin1

    return kin1, kin2

## michaelis - menten with no co-sub
@numba.njit
def mu(m1,m2,s0,s1,e2,f2,k2,l1):
    return (e2*l1*m1*s0 - f2*k2*m2*s1)/(k2*l1+k2*m2*s1+l1*m1*s0)

####  ODES for m01, m02, m11, m12, a0 and a1 ####
@numba.njit
def dm01_dt(P,t,kin1):
    m01=P[0]
    m11=P[2]
    a0=P[4]
    a1=P[5]
    return kin1- mu(m01,m11,a0,a1,E11,F11,K11,L01)

@numba.njit
def dm02_dt(P,t,kin2):
    m02=P[1]
    m12=P[3]
    a0=P[4]
    a1=P[5]
    return kin2- mu(m02,m12,a1,a0,E12,F12,K12,L02)

@numba.njit
def dm11_dt(P,t):
    m01=P[0]
    m11=P[2]
    a0=P[4]
    a1=P[5]
    return mu(m01,m11,a0,a1,E11,F11,K11,L01) - kout1*m11

@numba.njit
def dm12_dt(P,t):
    m02=P[1]
    m12=P[3]
    a0=P[4]
    a1=P[5]
    return mu(m02,m12,a1,a0,E12,F12,K12,L02) -kout2*m12

@numba.njit
def da0_dt(P,t):
    m01=P[0]
    m02=P[1]
    m11=P[2]
    m12=P[3]
    a0=P[4]
    a1=P[5]
    return mu(m02,m12,a1,a0,E12,F12,K12,L02) - mu(m01,m11,a0,a1,E11,F11,K11,L01)\
        -mu(a0,a1,1.,1.,Ea,Fa,Ka,La)

@numba.njit
def da1_dt(P,t):
    m01=P[0]
    m02=P[1]
    m11=P[2]
    m12=P[3]
    a0=P[4]
    a1=P[5]
    return  mu(m01,m11,a0,a1,E11,F11,K11,L01) -mu(m02,m12,a1,a0,E12,F12,K12,L02)\
        +mu(a0,a1,1.,1.,Ea,Fa,Ka,La)

#### general function that collects the above into a vector
@numba.njit
def dP_dt(P,t,kin1,kin2):
    return [dm01_dt(P,t,kin1), dm02_dt(P,t,kin2), dm11_dt(P,t), dm12_dt(P,t), 
            da0_dt(P,t), da1_dt(P,t)]


### calculate the next time to change the kin values
@numba.njit    
def get_new_env_t(t0,tau,rand):
    
    t_add=-np.log(rand)*tau
    return t0 + t_add

### wrapper that generates the time series
def get_kratio_paths(params,time_points,pop_init):  
    
    ### parameters for the noise - var is the variance of the log ratio
    ### kav is the average of the kin values
    ### tau is the correlation time of the noise - i.e. the average time
    ### between changes
    var=float(params_noise[0])
    kav=float(params_noise[1])
    tau=float(params_noise[2])
    
    ### generate the initial kin values
    kin1_init,kin2_init=get_kratio_vals(var,kav)
    
    ### i keeps track of the specified time point
    i=0
    t=time_points[0]
    
    ### get the initial population size from the inputs, generate a matrix 
    ### to store the time series in and then store the initial values in the
    ### first row
    Pinit=pop_init
    P_out=np.zeros((len(time_points),6))
    P_out[0,:]=Pinit
    
    ### find out how many kin changes we will roughly have, then pregenerate
    ### this many uniformly distributed random variables
    num_env_vals=np.max([int(10*Tmax/tau),10])
    env_switch_vals=np.random.rand(num_env_vals)
    env_switch_counter=1
    
    ### find the time until the next switch
    t_env=get_new_env_t(t, tau,env_switch_vals[0])
    
    ### get the kin values from the initial values
    kin1=kin1_init
    kin2=kin2_init
    
    ### vector to store the kkin ratios
    kratio_out=np.zeros((len(time_points)))
    kratio_out[0]=kin2/kin1
    ### loop over the sequential time points from the input
    for i in range(len(time_points)-1):
        
        ### find the next time point 
        t_next=time_points[i+1]
        
        ### while the time for next switch is less than the next time point,
        ### run the ODE system and then find the next kin change time and find
        ### the new kin values
        while t_env< t_next:
            Ps_temp=odeint(dP_dt, Pinit, [t,t_env],args=(kin1,kin2))
            Pinit=Ps_temp[-1,:]
            t=t_env
            
            t_env=get_new_env_t(t, tau,env_switch_vals[env_switch_counter])
            env_switch_counter+=1
            kin1,kin2=get_kratio_vals(var,kav)
        
        ### once the time for the next kin change is greater than the next time
        ### point, run the ODE system until this time point
        Ps_temp=odeint(dP_dt, Pinit, [t,t_next],args=(kin1,kin2))
        
        ### save the final value to be the initial values in the next run
        Pinit=Ps_temp[-1,:]
        ### save the final time to be the initial in the next run
        t=t_next
        ### save the metabolite concentrations in the matrix and the kratio 
        ### values in their vector
        P_out[i+1,:]=Pinit
        kratio_out[i+1]=kin2/kin1
    return P_out, kratio_out


if wantheatmap==1:
    
    ### generate the tau values and then the corners for the heatmap ###
    tauvals=np.logspace(-2,taumax,numtauvals)
    intervaltauvals=np.log10(tauvals[1]/tauvals[0])
    tauvals_plot=np.logspace(np.log10(tauvals[0])-intervaltauvals/2.,
                            np.log10(tauvals[-1])+intervaltauvals/2.,numtauvals+1)
    
    ### generate the atot values and then the corners for the heatmap ###
    Atotvals=np.logspace(-2,atotmax,numAvals)
    intervalAvals=np.log10(Atotvals[1]/Atotvals[0])
    Atotvals_plot=np.logspace(np.log10(Atotvals[0])-intervalAvals/2.,
                              np.log10(Atotvals[-1])+intervalAvals/2.,numAvals+1)
    
    ### skip the first 100 time points when calculating the population
    ### correlations - this is to avoid erroneous results due to initial values
    t_skip=100.
    

    
    ### generate a matrix to store the correlation results in
    corr_tau_conc_mat=np.zeros((numtauvals,numAvals))
    
    ### start timer to see how long the matrix genration takes
    timer0=time.perf_counter()
    
    ### loop over the cosub concentrations
    for ii in range(len(Atotvals)):
        ### get the initial A0 and A1 values
        Atot=Atotvals[ii]
        A0_init1=Atot/2.
        A1_init1=Atot/2.
        
        ### get the initial population vector
        popinit=np.array([M01_init, M02_init, M11_init, M12_init, A0_init1, A1_init1])
       
        ### loop over the tau values
        for i in range(len(tauvals)):
            tau=tauvals[i]
            
            ### set the maximum time so that there will be at least of order 
            ### 100 kin changes
            Tmax=np.max([1000.,100*tau])
            
            ### set the interval at which to record the metabolite concentrations
            ### and then generate these time points
            dt=0.01
            time_points=np.linspace(0,Tmax,int(Tmax/dt) +1)
            print('Starting sim with (log10(tau),log10(Atot)) = ',
                      (np.log10([tau, Atot])),
                      ', and Tmax,dt = ',(Tmax,dt))
            params_noise=np.array([var,kav,tau])
           
            ### get the metabolite time series 
            Pout,kratio_out=get_kratio_paths(params_noise,time_points,popinit)
            
            
            ### get M11 and M12 time series from the output and skip the first
            ### tskip values
            M11=Pout[time_points>t_skip,2]
            M12=Pout[time_points>t_skip,3]
            
            ### find the correlation between them and save it in the matrix
            conc_corr=np.corrcoef(M11, M12)[0,1]
            corr_tau_conc_mat[i,ii]=conc_corr
            

            print(conc_corr)
    
    timer1=time.perf_counter()
    
    ### print how long this took
    print(f"time for simulations was {(timer1-timer0)/60: 0.4f} minutes")
    
    ### plot the heatmap
    plt.figure()
    plt.pcolormesh(Atotvals_plot,tauvals_plot,corr_tau_conc_mat,vmin=-1,vmax=1,cmap='plasma')
    plt.yscale('log')
    plt.xscale('log')
    plt.colorbar()
    plt.xlabel('$T_{A}$')
    plt.ylabel('$\\tau$')
    titstr='$k_{av} = '+str(kav)+', V_{max,E_a} = '+str(Ea)+'$'
    plt.title(titstr)
    
    plt.plot(Atot_ts,tau_ts,'ok',markersize=10)


### tau and atot as defined in the preamble
tau=tau_ts
Atot=Atot_ts

Tmax=np.max([1000.,100*tau])
dt=0.01
time_points=np.linspace(0,Tmax,int(Tmax/dt) +1)

A0_init1=Atot/2.
A1_init1=Atot/2.
popinit=np.array([M01_init, M02_init, M11_init, M12_init, A0_init1, A1_init1])

params_noise=np.array([var,kav,tau])

## get the time series
Pout,kratio_out=get_kratio_paths(params_noise,time_points,popinit)


M01=Pout[:,0]
M02=Pout[:,1]
M11=Pout[:,2]
M12=Pout[:,3]
A0=Pout[:,4]
A1=Pout[:,5]

ts=time_points[:]

plt.figure()
ax1=plt.axes([0.1,0.1,0.9,0.9])
ax1.set_yscale('log')
## plot the metabolite concentrations
ax1.plot(time_points,M01,'-',color=m01col,label = '$M_{0,1}$')
ax1.plot(time_points,M02,'-',color=m02col,label='$M_{0,2}$')
ax1.plot(time_points,M11,'-',color=m11col,label = '$M_{1,1}$')
ax1.plot(time_points,M12,'-',color=m12col,label='$M_{1,2}$')
plt.xlabel('$\\rm{Time}$',fontsize=25)
plt.ylabel('$\\rm (M)$',rotation=0, fontsize=15)

### plot the kin ratios and the cosub ratios on a twin axis (rhs)
ax2=ax1.twinx()
ax2.set_yscale('log')
ax2.plot(time_points,kratio_out,color=k_ratio_col)
ax2.plot(time_points,A0/A1,'-',color=a1col)

### puts the metabolite concentrations in front of the kin and cosub ratios
ax1.set_zorder(ax2.get_zorder()+1)
ax1.patch.set_visible(False)
```
